# Heterochromatin and RNAi act independently to ensure genome stability in Mucorales human fungal pathogens

**DOI:** 10.1101/2022.12.01.518749

**Authors:** María Isabel Navarro-Mendoza, Carlos Pérez-Arques, Joseph Heitman

## Abstract

Chromatin modifications play a fundamental role in controlling transcription and genome stability and yet despite their importance, are poorly understood in early-diverging fungi. We present the first comprehensive study of histone-lysine and DNA methyltransferases across the Mucoromycota, emphasizing heterochromatin formation pathways that rely on the Clr4 complex involved in H3K9-methylation, the Polycomb-repressive complex 2 driving H3K27-methylation, or DNMT1-like methyl-transferases that catalyze 5mC DNA methylation. Our analysis uncovered H3K9-methylated heterochromatin as the major chromatin modification repressing transcription in these fungi, which lack both Polycomb silencing and cytosine methylation. Although small RNAs generated by RNAi pathways facilitate the formation of heterochromatin in many eukaryotic organisms, we show that RNAi is not required to maintain either genomic or centromeric heterochromatin in *Mucor*. H3K9-methylation and RNAi act independently to control centromeric regions, suggesting a functional sub-specialization. Whereas the H3K9 methyltransferase Clr4 and heterochromatin formation are essential for cell viability, RNAi is dispensable for viability yet acts as the main epigenetic, regulatory force repressing transposition of centromeric GremLINE1 elements. Mutations inactivating canonical RNAi lead to rampant transposition and insertional inactivation of targets resulting in antimicrobial drug resistance. This fine-tuned, Rdrp2-dependent RNAi activity is critical for genome stability, restricting GremLINE1 retroelements to the centromeres where they occupy long heterochromatic islands. Taken together, our results suggest that RNAi and heterochromatin formation are independent genome defense and regulatory mechanisms in the Mucorales, contributing to a paradigm shift from the co-transcriptional gene silencing observed in fission yeasts to models in which heterochromatin and RNAi operate independently in early-diverging fungi.

## Introduction

Fungi are a diverse clade of eukaryotic organisms that have successfully adapted to niches across the planet (1). To do so, they have evolved ecological strategies to survive and compete in diverse environments (2). In some cases, these result in pathogenic interactions and host disease, causing economic and human losses (3). Adaptation to hosts is a consequence of different molecular mechanisms modulating gene expression, and understanding this provides insights to lessen the burden associated with fungal diseases (4, 5). Many of these mechanisms rely on epigenetic modifications that do not alter the DNA sequence but rather control chromatin accessibility by producing distinct chromatin states. The repressed chromatin state, known as heterochromatin, is frequently associated with low gene density and highly repetitive DNA content, and heterochromatic regions are marked by methylation of histone H3 lysine 9 (H3K9me) and/or lysine 27 (H3K27me), considered as constitutive or facultative heterochromatin, respectively. Methylation at these sites is catalyzed by histone lysine methyltransferases (or KMT enzymes) (6). In addition, other epigenetic modifications may contribute to heterochromatin formation, including cytosine methylation at the carbon-5 position (5mC) catalyzed by DNA methyltransferases (DNMT) (7). These “writer” proteins add the methylation mark, as opposed to “eraser” demethylases that remove it; and specific “reader” proteins, which recognize epigenetic marks for signaling, and recruit “writers” to their targets for reinforcement (8, 9).

In the fission yeast *Schizosaccharomyces pombe*, pericentric heterochromatin formation by co-transcriptional gene silencing (CTGS) serves as a paradigmatic example. RNA polymerase II-based transcription from repetitive sequences triggers RNA interference (RNAi), generating small noncoding RNAs (sRNAs) that guide Argonaute and associated RNAi induced transcriptional silencing (RITS) complex components to complementary nascent transcripts (10–12). Next, the RITS complex recruits the “writer” methyltransferase Clr4/KMT1 to catalyze H3K9me (13), and methylation is reinforced, maintained, and extended by several chromodomain-containing “readers”: Chp1 from the RITS complex (14), the heterochromatin protein 1 (HP1) proteins Chp2 and Swi6 (15, 16), and Clr4 (17). Beyond this paradigm, heterochromatin formation has been characterized in ascomycetous and basidiomycetous fungi revealing new insights into 5mC DNA methylation and Polycomb silencing through KMT6 mediated H3K27me (18–24). Interestingly, *S. pombe* lacks both DNA methylation and H3K27me and the necessary “writer” proteins to produce these, highlighting the importance of alternative fungal models and developing techniques to study them. The fact that the Dikarya –Ascomycota and Basidiomycota– comprises only two out of the numerous fungal phyla suggests other regulatory, epigenetic mechanisms remain to be discovered in the early-diverging fungi, which the latest classifications estimate may comprise as many as seventeen phyla (25, 26).

The Mucorales are a group of early-diverging fungal pathogens known to cause devastating infections called mucormycoses. These infections cause significant morbidity and mortality (27), and mechanisms leading to antifungal resistance and pathogenesis are poorly understood. The fungus *Mucor lusitanicus* serves a model for this group of pathogens. Molecular techniques have been developed and applied to characterize RNA interference (RNAi) as a post transcriptional gene silencing (PTGS) mechanism (28). Doublestranded RNA (dsRNA) produced by RNA-dependent RNA polymerases (RdRP) (29, 30) is cleaved into sRNAs by Dicer ribonucleases (30–32). sRNAs are loaded into the RNAi-induced silencing complex (RISC) where Argonaute (Ago) directs degradation of the complementary messenger RNA by base-pairing homology and ribonuclease activity (33, 34). The canonical RNAi machinery competes with an alternative RNAi degradative pathway relying on RdRPs and the novel ribonuclease R3B2 to silence its own set of target transcripts (35, 36). These RNAi pathways modulate distinct biological processes including pathogenesis, regulation of centromeric transposons, and epigenetic antimicrobial drug resistance (36–41).

RNAi-based epimutations were discovered for the first time in *Mucor lusitanicus* and *Mucor circinelloides*, causing drug resistance by sRNA silencing of the *fkbA* gene encoding the FK506 target, FKBP12 (37). Later, epimutations were found to silence other drug targets, e.g., uracil biosynthetic genes targeted by pyrimidine analogs (5-FOA) (39). These epimutations result in unstable, transient drug resistances that revert to sensitivity after several mitotic growth cycles in the absence of drug. During host infection, epimutations persist and are induced in specific organs, suggesting a role in pathogenesis (42). Epimutations that promote drug resistance and host colonization pose a concern both for public health and agriculture (24, 37, 43–45), as they could elude detection due to their transient, unstable nature. Therefore, it is critical to elucidate the repertoire of epigenetic modifications these pathogens deploy, and how these pathways operate and interact to modify gene expression in response to environmental cues.

The diversity of RNAi mechanisms in *Mucor* spp. suggests distinct roles and functional specialization that may involve recruiting other epigenetic modifications as seen in several other eukaryotes (46–48). Here, we define the main forces driving heterochromatin formation in the Mucorales, characterize the role of RNAi and other epigenetic modifications in establishing and maintaining altered chromatin states, and determining the impact of epigenetic regulation on centromere identity and genome stability.

## Results

### Clr4-mediated histone-lysine 9 methylation is an essential regulatory mechanism in Mucorales

The Mucorales express a diverse repertoire of RNAi components that are involved in distinct repressive, regulatory pathways relying on sRNA-directed degradation of complementary mRNAs that results in PTGS. These sRNAs also serve as initiators of chromatin modifications leading to transcriptional repression in other eukaryotic organisms (12, 46, 48). While many of these modifications have been characterized in the Dikarya, their conservation and role in early diverging fungi was unexplored. Therefore, the presence of chromatin modifications was first surveyed in early-diverging fungal genomes, focusing on the Mucorales. As in most eukaryotes, Mucoralean histone proteins H3 and H4 are highly conserved (40). The *M. lusitanicus* genome encodes four histone H3 and three histone H4 copies (*SI Appendix*, Fig. S1), and the KMT-lysine substrates and neighboring regions are also conserved. Thus, these proteins could harbor chromatin modifications catalyzed by KMT enzymes or other histone-lysine modifiers. To identify RNAi components, and KMT and DNMT homologs in the Mucorales, 86 fungal proteomes were analyzed including at least one representative of most fungal phyla, focusing on the Mucoromycota phylum and major fungal pathogens (*SI Appendix*, Dataset S1). Out of these proteomes, 43 were selected as representatives to capture the presence or absence of different epigenetic modifying enzymes (Fig. 1). We identified homologs of the three major RNAi enzymes –RdRP, Dicer, and Ago– across all fungal phyla and particularly, in every Mucoromycota species, though not in some isolated Dikarya species or clades, e.g., *Saccharomyces cerevisiae, Candida auris, Ustilago maydis*, and *Malassezia pachydermatis*. These losses are well documented (49–51) and validate the approach. In *S. pombe*, CTGS leading to constitutive heterochromatin initiates in regions producing complementary sRNA that are loaded into the RITS complex by the Argonaute siRNA chaperones (ARC), Arb1 and Arb2. These two proteins are mostly conserved in the Mucorales, ascomycetes that have retained RNAi, and some early-diverging fungi. However, RITS-exclusive components –Chp1 and Tas3– are only found in *S. pombe*, indicating that any possible interplay among RNAi and heterochromatin formation might require novel components.

**Fig. 1.**
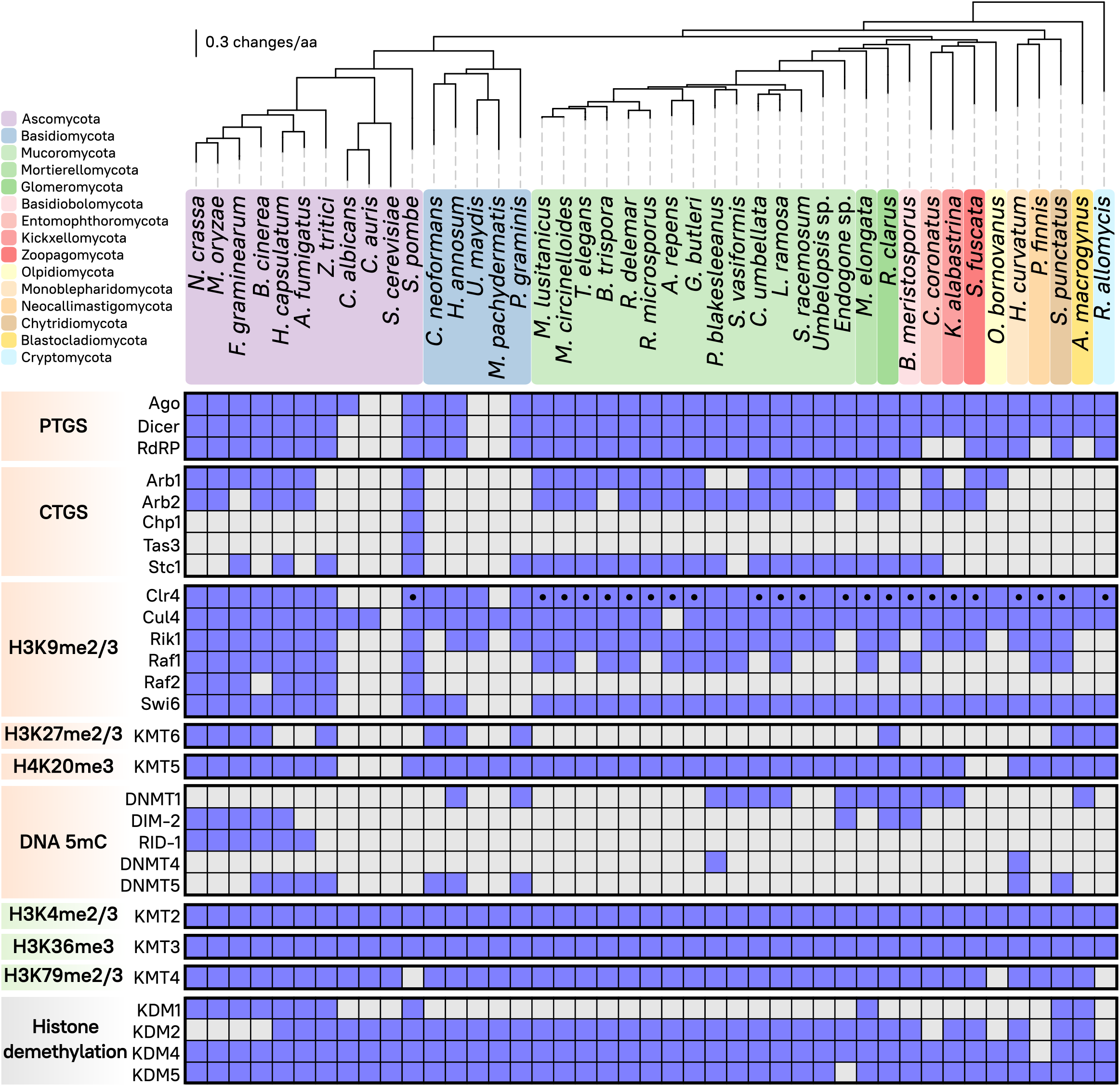
Distribution of RNA interference components and epigenetic regulatory methyltransferases in Fungi. The matrix displays presence (blue) or absence (gray) of different epigenetic regulatory complexes and their corresponding proteins (left) across the fungal kingdom (top). Predicted N-terminal chromodomain in Clr4 methyltransferase is depicted (black dot). RNAi components involved in post- or co-transcriptional silencing (PTGS or CTGS respectively, orange), methyltransferases involved in either transcriptional activation (green) or repression (orange), as well as histone demethylases (gray) are shown. The fungal phylogenomic tree contains 43 species selected as representatives from each major color-coded phylum and a robust sampling of Mucoromycota and other fungal pathogens, showing evolutionary distance as substitutions per amino acid (solid black lines).

Next, KMT homologs and accessory proteins in the methyltransferase complexes were identified. KMT enzymes are often composed of a SET domain which is involved in methyltransferase activity, and a variety of other motifs that mediate binding to specific lysine targets and accessory proteins. Clr4/KMT1 (also known as DIM-5, Su(var)3-9, and SUV39H), KMT6 (SET-7, E(z), and EZH2), and KMT5 (Set9, SET-10, Hmt4-20, and SUV420H) catalyze methylation of H3K9, H3K27, and H4K20, respectively (*SI Appendix*, Fig. S1), resulting in transcriptional repression. Clr4 and associated proteins, Rik1 and the ubiquitin ligase Cul4, are highly conserved in fungi, particularly in the Mucorales (Fig. 1). In *S. pombe*, the Clr4 complex (CLRC) has been shown to interact with Ago1 through Stc1, which connects both pathways to establish H3K9me via RNAi (50). Interestingly, Stc1 homologs are present in the Mucoromycota despite the lack of other RITS components. The heterochromatin protein 1 (HP1) complex (Swi6/Chp2 homologs) is also conserved among the Mucoromycota, and all early-diverging fungi that have retained RNAi. Surprisingly, KMT6 was lost in every Mucoromycota species, yet is conserved in some other early-diverging fungal phyla. It was also lost in some Ascomycota and many Basidiomycota, suggesting that H3K27me and Polycomb silencing was lost in many independent events during fungal evolution.

We also searched for DNMT enzymes, specifically those capable of producing transcription-repressive 5mC DNA methylation (Fig. 1). Fungal specific *de novo* methyltransferase RID-1 is exclusively conserved in the Pezizomycotina, while DIM-2-like homologs were identified in early-diverging Endogonales and Glomeromycota. Traditionally, DIM-2 and RID-1 were considered the main *de novo* DNA methyltransferases operating in fungi, and recent phylogenetic studies aligns them with a monophyletic clade that includes DNMT1 from animals and plants (52). While DNMT1 homologs were identified in some Mucorales and other early diverging fungi, it was notably absent in the pathogenic *Mucor* and *Rhizopus* species.

Other KMT enzymes are involved in transcription activation: KMT2 (Set1, SET-1, and SETD1), KMT3 (Set2, SET-2, and SET2D), and KMT4 (Dot1, DOT-1/HLM-2, Gpp, and DOT1L) methylate H3K4, H3K36, and H3K79, respectively. These KMT enzymes are conserved across fungal evolution (Fig. 1), except for the H3K79 methyltransferase KMT4 in *S. pombe*. The dynamics of histone-lysine methylation are also controlled by “eraser” enzymes known as histone demethylases (KDMs) that catalyze the opposite reaction, removing methylation marks. KDM1, KDM2, KDM4, and KDM5 are mostly conserved in fungi, except for KDM1 (Lsd1 and AOF2) that is absent in the Basidiomycota, Mucoromycota, and other lineages of early-diverging fungi; and KDM2 (JHD1, Epe1, and JHDM1) loss in several filamentous ascomycetes.

Overall, most Mucoromycota species have lost KMT enzymes and associated proteins involved in H3K27me-mediated Polycomb silencing through facultative heterochromatin, as well as DNMT proteins catalyzing DNA 5mC. Because Clr4/KMT1 is the only transcription-repressive chromatin-modifying enzyme present in every Mucoralean species analyzed, we focused on the *M. lusitanicus clr4* gene. First, the previous *clr4* gene model prediction was curated and reannotated using our RNA sequencing transcriptomic data. This revealed three additional exons at the 5’-end of the reannotated gene, encoding a predicted chromodomain essential for H3K9me binding activity (*SI Appendix*, Fig. S2A). The presence of a chromodomain in the *M. lusitanicus* Clr4 protein prompted us to inspect every Clr4 homolog in species with available transcriptomic data and alternative gene models. We found many N-terminally truncated, misannotated proteins and uncovered as many chromodomains in the early-diverging fungi (*SI Appendix*, Fig. S2B, Dataset S2 Clr4/KMT1). A phylogenetic analysis across the fungal kingdom revealed that most fungal, specifically early-diverging, Clr4 enzymes harbor an N-terminal chromodomain, suggesting six major and independent losses of this domain in the last common ancestors of the Basidiomycota, Saccharomycotina and Pezizomycotina, Phycomycetaceae, Umbelopsi-dales, Olpidiomycota, and Blastocladiomycota (Fig. 1, *SI Appendix*, Fig. S2B).

To gain a better understanding of Clr4 function in Mucorales, we attempted to delete *clr4* through CRISPR-Cas9 and homologous recombination in *M. lusitanicus*. Mucorales have non-septate hyphae and multinucleated asexual spores. As such, genetic transformation often results in initially heterokaryotic mutants, producing spores that harbor both wildtype and mutant nuclei. Mutant allele segregation by sexual reproduction is not possible because *M. lusitanicus* is not able to undergo a complete sexual cycle under laboratory conditions. To render these mutants homokaryotic, i.e., harboring mutant nuclei exclusively, transformants must be passaged through several vegetative growth cycles in a medium that positively selects the mutant allele. Surprisingly, every transformation experiment aiming at *clr4* deletion resulted in heterokaryotic mutants that were unable to achieve complete mutant homokaryosis, and after twenty vegetative passages they still harbored wild-type nuclei (*SI Appendix*, Fig. S3). We tested if Clr4 activity could be essential for viability by genotyping the asexual progeny of a heterokaryotic mutant, reasoning an equal rate of wild-type or mutant homokaryons in the spore population is expected for a non-deleterious mutation. To do so, 40 spores from a heterokaryotic mutant were dissected and plated on non-selective medium, and after incubation, 30 spores were able to germinate. Next, we analyzed the germinated spores and found either wild-type homokaryotic or heterokaryotic colonies (with wild-type and *clr4*Δ nuclei), yet no *clr4*Δ homokaryons were observed (*SI Appendix*, Fig. S3). These results indicate that homokaryotic *clr4*Δ spores are not able to survive, implying Clr4 is essential and suggesting that a functional H3K9me-mediated heterochromatin formation mechanism is required for cell viability in *M. lusitanicus*.

### Regions targeted by H3K9 methylation are transcriptional deserts enriched with repeats

Chromatin immunoprecipitation followed by Illumina sequencing (ChIP-seq) using α-H3K9 di- (H3K9me2) or tri-methylated (H3K9me3) antibodies was performed to characterize H3K9me in *M. lusitanicus*. Reads were mapped to the publicly available *M. lusitanicus* MU402 v1.0 genome (https://mycocosm.jgi.doe.gov/Muccir1_3/Muccir1_3.home.html), and H3K9me2 and -me3 enriched peaks were identified across the genome. Genome-wide observations (Fig. 2A and *SI Appendix*, Fig. S4) revealed a striking overlapping pattern between H3K9me2 and -me3, which was confirmed by linear correlation measurements of every enriched peak detected (Fig. 2B).

**Fig. 2.**
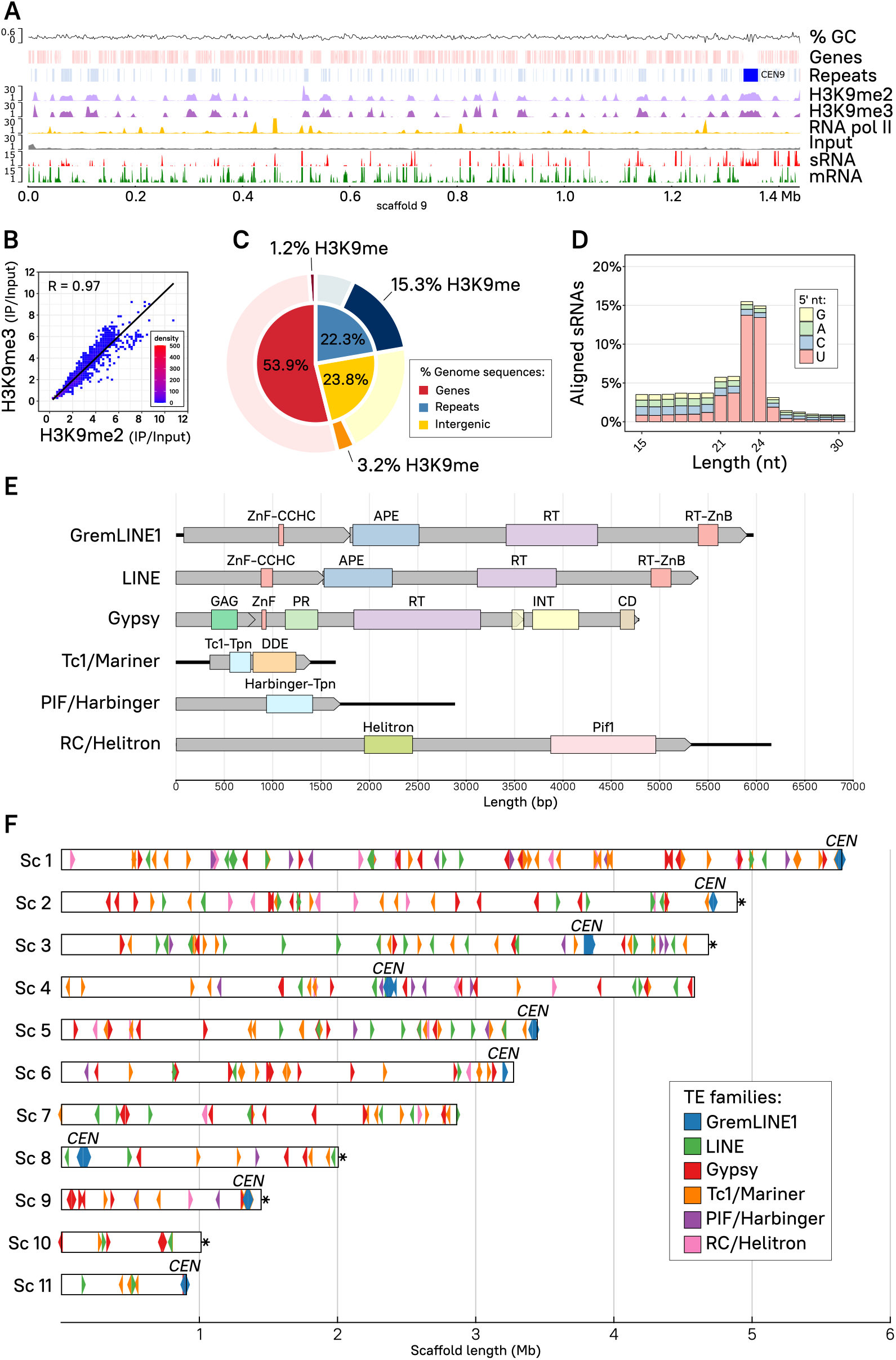
H3K9me heterochromatin and RNAi are coincident at transcriptionally silent, repeated sequences. **(A)** H3K9me2 (light purple) and -me3 (dark purple) and RNA polymerase II (RNA pol II, gold) enriched distribution is shown across scaffold 9. Enrichment coverage is computed as the ratio of immunoprecipitated (IP) DNA/Input DNA reads, previously normalized as log_2_ counts per million (CPM). Input control coverage (gray) is shown as normalized log_2_ CPM for reference. Transcripts (mRNA, green) and sRNA (red) coverages correspond to log_2_ CPM. Genes (light red) and repeated sequences (light blue) including pericentric regions (bright blue) are displayed, as well as GC content coverage as a percentage. **(B)** Dot plot and Pearson’s correlation coefficient (R, dark line) of H3K9me2 (x-axis) and -me3 (y axis) average log_2_ CPM enrichment values are plotted as 10-kb regions representing the whole genome. Overlapping dots are color-coded as a density scale. **(C)** Proportion of gene, repeated, and intergenic sequences are shown color-coded as percentage of total base pairs in the genome (inner pie chart). The proportion of H3K9-methylated base pairs in the same categories is highlighted (darker shade, outer pie chart) from the remaining non-methylated base pairs (lighter shade, outer pie chart). **(D)** The percentage of small RNA aligned reads according to length and 5’-end nucleotide distribution (color-coded) is depicted. **(E)** Schematic view of the six most abundant TE families identified in the genome, showing total length (bp) of the elements (solid black lines), open reading frames (gray rectangular arrows), and predicted protein domains (color coded rectangles) abbreviated as follows: Zinc finger, CCHC-type (ZnF-CCHC); Endonuclease/exonuclease/phosphatase superfamily (APE); Reverse transcriptase domain (RT); Reverse transcriptase zinc-binding domain (RT-ZnB); Retrotransposon GAG domain (GAG); Aspartic peptidase domain superfamily (PR); Integrase zinc-binding domain and catalytic core (INT); Chromodomain (CD); Homeobox-like domain superfamily (HOX); Transposase, Tc1 like (Tc1-Tpn) and DDE domain (DDE); Harbinger transposase-derived nuclease domain (Harbinger-Tpn); Helitron helicase-like domain (Helitron); DNA helicase Pif1-like (Pif1). **(F)** Genomic distribution of the six most abundant TE families shown as color-coded rectangular arrows to indicate transcription direction (element length is scaled to the rectangle width, not including the arrow point). Centromeric (*CEN)* and telomeric (*) regions are designated in the plot. The first eleven out of twenty-four scaffolds (> 94% of the genome) containing nine centromeres and their length (Mb) are shown for reference.

To determine if H3K9me-mediated heterochromatin results in transcriptional silencing, ChIP-seq against RNA polymerase II (RNA pol II) and ribosomal RNA-depleted strand-specific RNA sequencing (RNA-seq) were performed. H3K9me peaks emerge as heterochromatin islands in transcriptional deserts, which are devoid of mRNA and RNA pol II occupancy (Fig. 2A and *SI Appendix*, Fig. S4). Interestingly, genes are absent from most of these heterochromatic regions, suggesting H3K9me is targeting other genomic elements. A *de novo* repeat identification analysis was conducted across the genome of *M. lusitanicus*, resulting in an estimate of 22.3% genome-wide repeated DNA content, 53.9% gene sequences, and the remaining percent intergenic sequences. An in-depth, base pair-resolution analysis confirmed the genome-wide observations, showing that ~19.7% of the genome is bound by H3K9me chromatin which is mainly directed at repeats (15.3%, Fig. 2C). In contrast, only 1.2% of the genome is targeted by H3K9me across coding sequences, suggesting that H3K9me is mostly controlling repeats rather than regulating gene expression.

Because previous studies have shown that *M. lusitanicus* Dicer-dependent, canonical sRNAs are able to target repeated and transposon sequences (30, 36) and most of the H3K9me islands are localized to repeats, the sRNA distribution across the whole genome was analyzed by conducting sRNA-enriched RNA purification and sequencing (sRNA-seq). The sRNA content analyzed exhibits key features of canonical sRNAs: 1) a discrete length ranging from 21 to 24 nt and 2) a strong preference for uracil at the 5’-end (Fig. 2D). These canonical sRNAs coincide extensively with H3K9me at silenced repeated regions (Fig. 2A and *SI Appendix*, Fig. S4), indicating a dual regulation by post-transcriptional and transcriptional silencing mechanisms. In many eukaryotic genomes, repeated regions represent copies of transposable elements (TE) whose transposition gave rise to fine-tuned regulatory mechanisms that shaped their genome evolution (53).

To determine the extent of transposable elements regulated by either or both H3K9me and RNAi, we conducted the first comprehensive TE identification analysis in the *Mucor* species complex. Briefly, open reading frames (ORFs) were predicted in every repeat type identified in our previous analysis, and searched for protein domains frequently associated with TE activity. The presence and relative position of these domains within each sequence was used to manually curate TE predictions, identifying six different TE families: DNA or Class II elements Tc1/Mariner, PIF/Harbinger, and RC/Helitron; and RNA or Class I elements non-LTR GremLINE1 and other LINE, and LTR Ty3/Gypsy (Fig. 2E, *SI Appendix*, Table S1, Dataset S3). All of these families comprised several distinct elements that were classified into subfamilies according to sequence similarity, particularly the previously identified centromeric GremLINE1 retrotransposons (40). After generating this curated TE library, the whole genome and annotated TE sequences were screened using restrictive parameters, ensuring only highly similar and full-length elements were included. Then, TE distribution across the genome was examined to detect conspicuous clustering patterns (Fig. 2F). Only the GremLINE1 elements occupy a specific genomic region, defining the pericentric regions of *M. lusitanicus*, which are transcriptionally silent regions devoid of genes (40). Conversely, the remaining TE families have a cosmopolitan distribution across the genome. Telomeric regions containing the repeated sequence 5-TTAGGG-3 were identified at the 3’-end of scaffolds 2, 3, 8, 9, and 10 (indicated as asterisks in Fig. 2F). These telomeric and subtelomeric (~50 kb) regions were not enriched in TE of any family, although several structural repeats and H3K9me marks were found at these regions (*SI Appendix*, Fig. S5A). Sex determination in Mucorales is controlled by the sex locus, comprising an allelic High-mobility group (HMG) transcription factor *–sexP* and *sexM* alleles– flanked by triose phosphate transporter (*tptA*) and RNA helicase (*rnhA*) encoding genes (54). Despite its epigenetic regulation in other fungal models (55), *M. lusitanicus* mating-type locus did not show any specific silencing mark (*SI Appendix*, Fig. S5B).

Taken together, these findings reveal an intimate overlap between H3K9me2 and -me3 islands primarily forming in repeated regions subject to transcriptional silencing. Interestingly, most of these genomic regions are also targeted by sRNAs, indicating a possible interconnection between RNAi and heterochromatin formation pathways to control transposition that was explored further.

### RNAi is dispensable for heterochromatin maintenance at transposable elements

RNAi relies on sRNAs to induce post-transcriptional silencing of complementary transcripts but in addition, these sRNAs may participate in CTGS by interacting with other repressive epigenetic mechanisms to initiate and maintain heterochromatin formation (12, 46–48). To explore the interaction between the principal epigenetic repressive mechanisms in *M. lusitanicus*, RNAi and H3K9me levels were compared in the newly identified TEs. Class I and II transposons are traditionally classified according to their transposition intermediates: RNA or DNA, respectively. An RNA intermediate could be more readily targeted by RNAi and consequently, the analysis was focused on RNA transposons. H3K9me binds to most copies of GremLINE1 and Gypsy, and several copies of LINE in the wild-type *M. lusitanicus* strain (Fig. 3). In these elements, H3K9me is deposited across the entire sequence, with the highest enrichment at the start and end positions. As expected, sRNAs are abundantly produced from these regions but interestingly, they target more TE copies than H3K9me. For instance, some LINE subfamilies generate sRNAs, but H3K9me is completely absent (Fig. 3, e.g., LINE subfamily highlighted in green). Nevertheless, none of these elements are being actively transcribed, not even those lacking H3K9me, as evidenced by the transcriptomic and RNA pol II occupancy data (Fig. 3). Because several TE copies are differentially targeted by H3K9me or sRNAs, these mechanisms may be acting independently to regulate these elements.

**Fig. 3.**
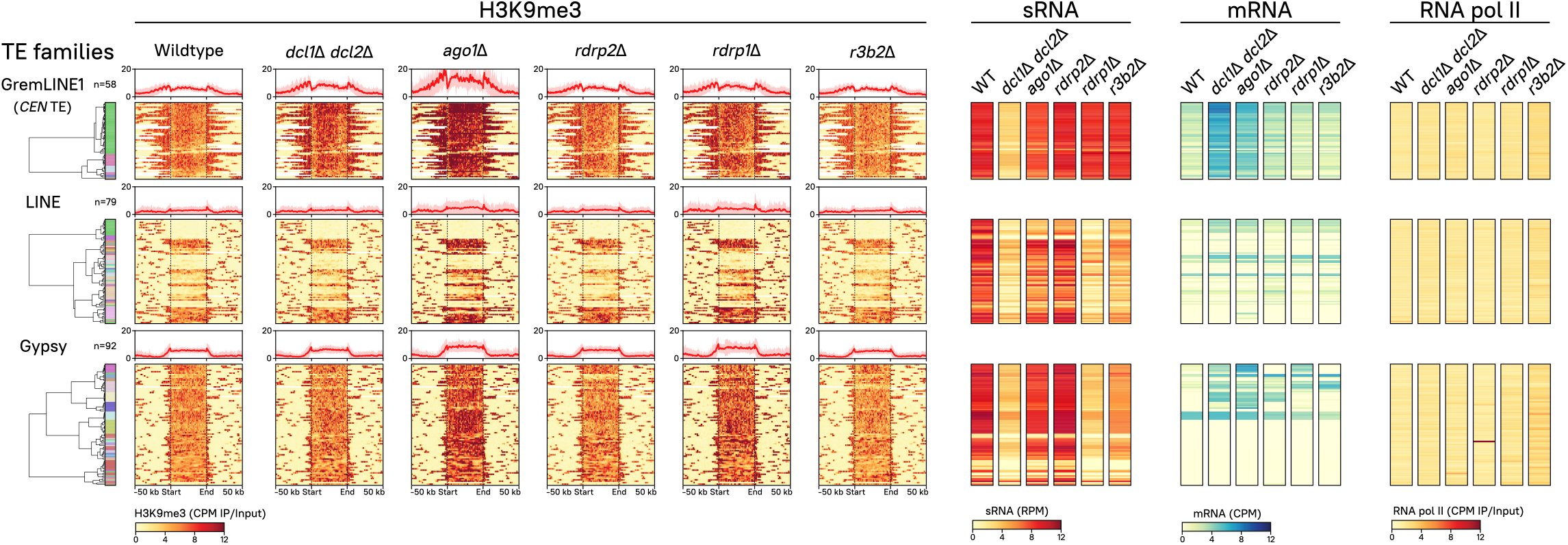
Dual epigenetic regulation –H3K9me-based heterochromatin and RNAi– targets RNA transposons. All copies of the GremLINE1 (n=58), other LINE (n=79), and Gypsy (n=92) retrotransposons identified in the genome are shown. TE copies from the same family are clustered according to their subfamily, H3K9me3, sRNA, and mRNA abundance in the wild-type strain. Heatmaps depicting H3K9me3, sRNA, mRNA, and RNA polymerase II abundance in wild-type and RNAi deficient strains are plotted (see plot name above). Each row represents a single TE copy, which are aligned across all heatmaps to compare their values that read from left to right as follows. First, a clustered dendrogram indicates each element subfamily (color-coded). Second, H3K9me3 enrichment values are shown as the immunoprecipitation (IP)/input ratio of normalized log_2_ CPM (CPM, yellow-to-red legend) from start to end of each copy (divided in 250 bins) and 50 kb upstream and downstream; as well as the mean value of all of the copies in each family shown in a profile plot (on top, bold red line shows the average and faded red background shows the standard deviation). Third, sRNA values normalized to log_2_ reads per million (RPM, yellow-to-red legend) are presented. Fourth, sense RNA values normalized to log_2_ CPM (yellow to-blue legend) are shown. Lastly, RNA pol II occupancy values are depicted as IP/Input ratio of normalized log_2_ CPM (yellow-to-red legend).

To investigate this hypothesis, an extensive mutant collection comprised of every key RNAi enzyme identified in *M. lusitanicus* was studied. To confirm their involvement in RNAi, sRNA content was sequenced and compared to previous studies (30, 36) (*SI Appendix*, Fig. S6A). These results recapitulate that sRNAs are almost depleted in *dcl1*Δ *dcl2*Δ, a double mutant lacking any Dicer activity; and in *ago1*Δ, lacking the main Argonaute enzyme, confirming that both activities are essential for sRNA biogenesis and canonical RNAi (*SI Appendix*, Fig. S6B). These canonical sRNAs are mostly produced from exons and repeated sequences (*SI Appendix*, Fig. S6C), suggesting gene and transposon regulation are their primary targets. Therefore, we focused our analysis on canonical (*dcl1*Δ *dcl2*Δ, *ago1*Δ, and *rdrp2*Δ) and alternative (*rdrp1*Δ and *r3b2*Δ) RNAi pathway mutants (Fig. 3). Other RNAi paralog mutants (*dcl1*Δ, *dcl2*Δ, *ago2*Δ, *ago3*Δ and, *rdrp3*Δ) are covered in supplementary material because their function is not essential for *M. lusitanicus* RNAi (*SI Appendix*, Fig. S7). ChIP and RNA-seq analyses were conducted to identify H3K9me2 and -me3 enrichment, as well as transcription across the genome.

Strikingly, not a single RNAi-defective mutant showed a significant decrease in H3K9me3 at any TE analyzed (Fig. 3, *SI Appendix*, Fig. S7), and no differences in H3K9me2 and me3 content were observed in any genomic region (*SI Appendix*, Fig. S8). These findings indicate that RNAi is neither sufficient nor necessary to maintain histone methylation across transposable elements. Despite not showing alterations in H3K9me, both mRNA expression and sRNA accumulation were substantially affected in the mutants. The production of sRNAs targeting most of these regions decreased dramatically in *dcl1*Δ *dcl2*Δ, and to a lesser extent in *ago1*Δ, as expected of Dicer- and Ago-dependent, canonical sRNAs. More importantly, the absence of Dicer or Ago activity increased transcript levels in many TE despite being embedded in H3K9me chromatin, especially but not restricted to Grem-LINE1 elements. Conversely, some TE subfamilies showed a marked decrease in canonical sRNA production in RNAi-defective mutants but remained transcriptionally silent or inactive, suggesting H3K9me and RNAi may operate independently to repress TE expression. Interestingly, RNA pol II occupancy remained stable in all of the RNAi mutants, indicating that increased transcript accumulation was a result of lacking PTGS instead of increased transcription. These results confirm that *M. lusitanicus* RNAi pathways act exclusively via PTGS.

*S. pombe* Rdp1 and the RNA-directed RNA polymerase complex (RDRC) are essential to initiate co-transcriptional gene silencing at pericentric repeats (56). In contrast, *M. lusitanicus* encodes three subspecialized RdRP enzymes (29, 57), suggesting more complex regulatory dynamics. To address this, sRNA production and transcript expression across TE sequences were studied in the RdRP and alternative RNAi pathway mutants. Surprisingly, obvious differences were observed in centromeric GremLINE1 retrotransposons compared to other elements (Fig. 3). Rdrp1 and R3B2 –the main components of the alternative pathway– are important to produce sRNAs from LINE and Gypsy retrotransposons and to regulate their expression, suggesting an interaction between canonical and alternative RNAi pathways to control these elements. In contrast, post-transcriptional silencing of GremLINE1 elements is completely dependent on the canonical pathway because sRNA and transcript levels remain unchanged in mutants of the alternative RNAi pathway. These differences in TE regulation suggest that centromeric Grem-LINE1s are controlled by a unique RNAi-based mechanism, possibly as a consequence of their role in centromere identity.

To confirm the extent of GremLINE1 unique regulation, the remaining DNA transposable elements (Tc1/Mariner, PIF/Harbinger, and RC/Helitron) were examined (*SI Appendix*, Fig. S9A). Most copies of DNA transposons show H3K9me across the coding sequence. Despite relying on a DNA intermediate for transposition, sRNAs are produced from Mariner and Harbinger copies as well as several Helitron subfamilies through canonical and alternative RNAi pathways. Only Mariner and Helitron subfamilies 1 (*SI Appendix*, Fig. S9A, green in both Mariner and Helitron) depend on RNAi to repress their expression as *dcl1*Δ *dcl2*Δ and *ago1*Δ exhibit increased transcript levels, similar to Gypsy and LINE RNA transposons. On the other hand, the remaining DNA element copies are transcriptionally silenced, possibly as a result of H3K9me. Due to the presence of full-length elements and the necessary protein products for autonomous transposition, we reasoned that transcripts encoding transposase protein domains might be targeted by RNAi. Indeed, sRNAs are specifically directed to the transposase-encoding transcript, suggesting RNAi is silencing transposase transcripts to prevent DNA transposition events as a genome defense mechanism (*SI Appendix*, Fig. S9B). Intriguingly, H3K9me3 is more broadly distributed, extending to neighboring structural repeats, reinforcing that histone methylation is maintained in regions not targeted by RNAi and that both epigenetic mechanisms may recognize and target repeats differently.

### RNAi prevents genome instability by repressing GremLINE1 transposition to non centromeric regions

GremLINE1 elements represent a novel LINE L1-like subfamily of RNA TEs exclusively located in pericentric regions and they are conserved in Mucoromycota species that lost CENP-A, suggesting an important role in centromere-kinetochore dynamics (40). We investigated the regulatory mechanisms controlling GremLINE1 expression, analyzing H3K9me and sRNA accumulation at the centromeres. H3K9me delimits the pericentric regions, extending further from the GremLINE1s and their complementary sRNAs into other repeated sequences, but sparing neighboring genes (Fig. 4A). Notably, H3K9me is absent at the kinetochore binding region, possibly to maintain an open chromatin state that allows kinetochore binding. Additionally, RNAi exerts fine-tuned, post-transcriptional silencing of GremLINE1 elements (Fig. 3) independently of Rdrp1 –the main RdRP associated with silencing initiation in *M. lusitanicus* (29).

**Fig. 4.**
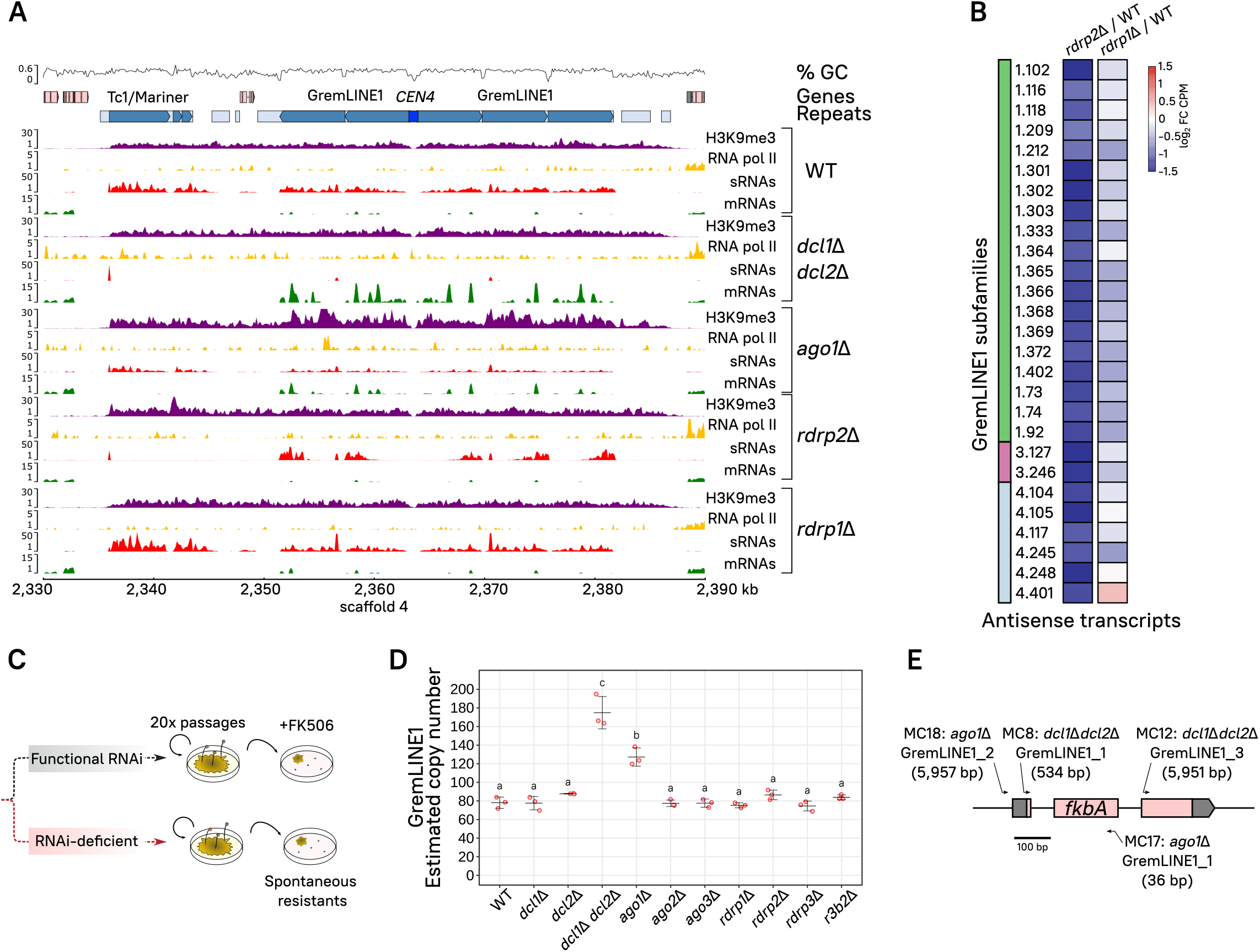
H3K9me defines pericentric boundaries while canonical RNAi inhibits GremLINE1 transposition to non-centromeric regions. **(A)** Genomic plot of *CEN4* and its pericentric regions displaying H3K9me3 enrichment (purple), and sRNA (red) and transcript levels (mRNA, green) in a wild-type (WT) and canonical RNAi-deficient strains (double *dcl1*Δ *dcl2*Δ and *ago1*Δ). Read coverage is normalized to log_2_ CPM, and ChIP enrichment is shown as IP/Input DNA ratio. GC proportion (black line) is computed across the whole sequence. Genes (light red), repeated sequences (structural repeats in light blue and TE in turquoise), as well as the centromeric sequence (bright blue) are shown for reference. **(B)** Heatmap of significant antisense transcription differences across GremLINE1 copies (FDR ≤ 0.05) in either *rdrp2*Δ or *rdrp1*Δ compared to the wild-type strain (color-coded values shown as log_2_ fold change of CPM). GremLINE1 subfamilies (color-coded) and identification are shown. **(C)** Artificial evolution experiment in RNAi mutants. Briefly, the strains were cultured through 20 vegetative growth passages, and analyzed for GremLINE1 copy numbers **(D)** or FK506 spontaneous resistance **(E)**. **(D)** GremLINE1 copies in different RNAi mutants after 20 vegetative passages are shown. Copy numbers are estimated as the relative fold (2^-ΔCT^) between GremLINE1 (multiple copies) and *vma1* (single copy) sequence amplification in triplicate quantitative PCR reactions. Significantly distinct values are indicated with different letters and were calculated by one-way ANOVA and Tukey’s HSD tests. **(E)** GremLINE1 transposition into the *fkbA* locus was positively selected on FK506-containing medium. Each insertion was named after their genetic background, the TE inserted, and a systematic number; insertion length (in bp) is also shown after the name. Insertion sites are indicated by arrows, whose direction follows the element orientation with respect to *fkbA*, and whose width reflects the scaled length of the target site duplication marking the insertion site.

Because all other TEs rely on Rdrp1 to generate sRNAs, we determined which RdRPs were triggering RNAi to target the GremLINE1s by searching for differences in natural antisense transcription and thus, dsRNA, across the GremLINE1 retrotransposons. Many GremLINE1 copies exhibited a significant decrease of antisense transcripts in *rdrp2*Δ compared to the wildtype and *rdrp1*Δ (Fig. 4B, FDR ≤ 0.05), indicating that Rdrp2 may generate dsRNA from these elements and induce RNAi. Rdrp2 is the only RdRP that harbors an RNA-recognition motif in its N-terminus (*Appendix SI*, Dataset S2 RdRP), and its activity is essential to maintain silencing through an sRNA-amplifying positive feedback loop (29). We propose that both activities, generating antisense transcription resulting in dsRNA and amplifying silencing, are important to initiate and maintain PTGS directed at the GremLINE1s, and could explain the slight increase of Grem-LINE1 transcript levels in *rdrp2*Δ (Fig. 3).

Our data confirms that GremLINE1 regulation is unique compared to other repeated elements outside of the centromeres, therefore we hypothesized that loss of RNAi could incite their transposition and lead to genome instability. To test this hypothesis, an artificial evolution experiment was conducted by culturing RNAi-defective and wild-type strains for twenty asexual passages in minimal medium (Fig. 4C). We estimated GremLINE1 copy number variation across these strains as the fold change compared to a single-copy gene by qPCR, resulting in a significant increase in *dcl1*Δ *dcl2*Δ and *ago1*Δ –key components of PTGS (Fig. 4D). Interestingly, transcript levels seem to correlate with the increase in copy number in each given mutant, particularly in *dcl1*Δ *dcl2*Δ and *ago1*Δ, but also in *dcl2*Δ and *rdrp2*Δ though their increase in copy numbers followed a trend that was not statistically significant (Fig. 3 and Fig. 4D).

To further characterize these transposition events, the strains were cultured under positive selection exerted by FK506, an antifungal drug that binds FKBP12, generating a protein-drug complex that inhibits calcineurin and enforces yeast-like growth. Mycelium-growing colonies were selected to capture GremLINE1s in the FKBP12-encoding locus *fkbA* and thus cause FKBP12 loss-of-function and FK506 resistance (Fig. 4C). After passage on the selective medium, 36 spontaneous FK506-resistant colonies were isolated: 10 in *dcl1*Δ *dcl2*Δ, 5 in *ago1*Δ, 1 in *rdrp2*Δ, 10 in *rdrp1*Δ, and 10 in the wild-type strains. Mutation types in each of these isolates were characterized by analyzing *fkbA* fragment size and whole-locus Sanger-sequencing, classifying these as insertions, point mutations or deletions, or epimutations when no genetic alteration was found (*SI Appendix*, Fig. S10, Table S2). Six insertions disrupting the *fkbA* locus were detected in only *dcl1*Δ *dcl2*Δ and *ago1*Δ resistant isolates. Four independent insertion events were fully characterized: one fulllength and one truncated GremLINE1 copies in each of the *dcl1*Δ *dcl2*Δ and *ago1*Δ mutants (Fig. 4E, *SI Appendix*, Fig. S10, and Table S3). In addition, two double GremLINE1 insertions were detected in *dcl1*Δ *dcl2*Δ, confirmed by PCR with a specific *fkbA* and GremLINE1 primer pair. Every insertion analyzed revealed exclusively a GremLINE1 copy, and the remaining resistant isolates in other mutant or wild-type strains did not harbor any insertions. The presence of A/T-5’-end target site duplications and truncated insertions reflect the LINE L1 canonical transposition mechanism, involving element-encoded endonuclease and reverse transcriptase activities (Fig. 2). Briefly, the endonuclease nicks genomic DNA in AT-rich regions generating an exposed 3’-OH end that primes a reverse transcription reaction of the A-rich 3’-end of the element (Fig. 5). Abrupt termination of the reverse transcription step frequently leads to incomplete insertions, explaining truncated copies (58). These findings strongly support that RNAi is preventing GremLINE1 transposition from H3K9-methylated pericentromeres to other genomic regions, limiting disruption of genes encoding antifungal drug targets.

**Fig. 5.**
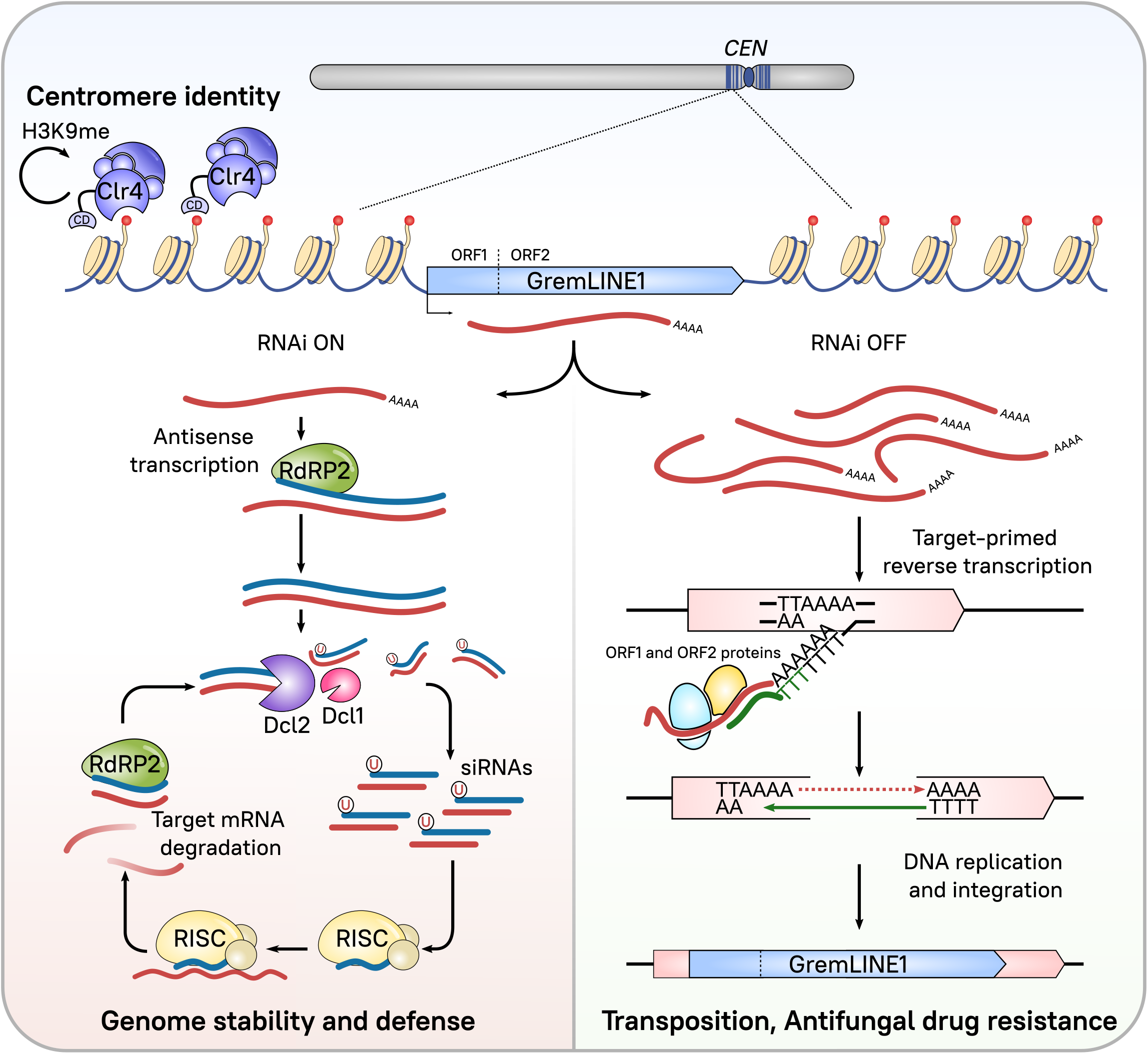
Current model for centromeric GremLINE1 dual regulation by H3K9me-based heterochromatin and RNAi. GremLINE1 copies are retrotransposons exclusively located in the pericentric regions of Mucoromycota species, forming long H3K9me heterochromatic islands that define centromere identity. Clr4 methyltransferase catalyzes H3K9me, and its chromodomain allows for H3K9me reading activity that could be involved in maintaining and spreading the mark in an RNAi-independent process. GremLINE1 elements display two ORFs encoding the enzymes necessary for their transposition, and their expression is usually post-transcriptionally repressed by RNAi in a process that involves Rdrp2 antisense transcription, Dcl2 nuclease activity that can be partially rescued by Dcl1, and RISC targeted degradation of GremLINE1 transcripts. In addition, Rdrp2 activity could reamplify the silencing signal by generating dsRNA from these degraded transcript fragments. This PTGS mechanism ensures genome stability by preventing GremLINE1 transposition and consequently, transposition is increased in RNAi-deficient strains. Lack of RNAi leads to increased GremLINE1 expression and consequently, to target-primed reverse transcription integration events in non-centromeric regions. These transposition events have been shown to affect drug targets, promoting drug resistance.

## Discussion

Here in this study, we have identified essential chromatin modifying enzymes across the fungal tree of life, focusing on the diverse yet understudied early-diverging fungi. Our findings imply that transcription-activating epigenetic mechanisms are ancestral and likely essential in fungi, present even in ascomycetous yeasts that lost many other chromatin modifying enzymes (59), which highlights the differences in transcriptional ground states between eukaryotic and prokaryotic organisms (60, 61). On the other hand, several transcriptional repressive chromatin-modifying enzymes are recursively lost in many fungi, except Clr4/KMT1 and its associated proteins from the CLRC that are conserved in most fungal lineages.

The vast majority of early-diverging fungal Clr4 proteins exhibit an N-terminal chromodomain, including those encoded in Mucoralean genomes. When present, the chromodomain H3K9me-binding activity is essential for heterochromatin maintenance and spreading to neighboring regions beyond the initiation sites in fungi (17) and animals (62). The presence of this chromodomain in fungal Clr4 enzymes is surprising because traditionally, *Schizosaccharomyces* spp. Clr4 was considered the only fungal Clr4 protein with a chromodomain, possibly due to a heavy bias towards Dikarya fungal models (9). Most Clr4 methyltransferases in animals (Su(var)3-9 or SUVH39) harbor an N-terminal chromodomain (63), whereas Clr4 enzymes in plants (SUVH) lack this domain (64). This convoluted Clr4/KMT1 evolutionary history of recurrent domain gain or loss led to the hypothesis that the fungal ancestor lacked the chromodomain (9, 65), yet our results indicate that the N-terminal chromodomain-containing variant is the ancestral Clr4 of animals and fungi.

Our research focuses on the human fungal pathogens Mucorales, and we have noted losses of certain chromatin modifying enzymes that must have important implications in their epigenetic regulatory landscape. For instance, the loss of KMT6 indicates the absence of H3K27me and Polycomb-mediated facultative heterochromatin. This is not the only relevant transcription-repressive modification lost in these fungi, because the majority of Mucorales also lack DNA 5mC methyltransferases. These mechanisms have been reported as essential for epimutations governing effector expression during host colonization in several plant pathogens (24, 44). Interestingly, *S. pombe* lost both Polycomb silencing and 5mC but still develops epimutations based on H3K9me heterochromatin –triggered by sRNAs– to acquire drug resistance (45). We reasoned that fungi may exploit any available epigenetic mechanism to modulate gene expression and better adapt to their current environment and therefore, emphasize how critical it is to characterize epigenetic modifications and their interactions that could contribute to epimutations in fungal pathogens.

Our results suggest that Clr4/KMT1, the CLRC, and HP1 complexes are the major chromatin modifying mechanisms involved in transcriptional silencing through heterochromatin in the Mucorales. Indeed, Clr4 is essential in *M. lusitanicus*, indicating that H3K9me repression controls critical biological processes. H3K9 is methylated sequentially from mono-to trimethylation, and these differentially methylated states define distinct domains of silent chromatin in many organisms, including mammals (66), nematodes (67), and plants (68). All of these organisms either lost the chromodomain necessary for Clr4 reading activity and/or possess a set of subspecialized Clr4 variants mediating the different H3K9 methylation states. In *M. lusitanicus*, H3K9me2 and -me3 enriched regions overlap and show a robust correlation across the whole genome and particularly, at repeated sequences. Similarly, wild-type *S. pombe* cells exhibit a strong correlation between H3K9me2 and -me3 in pericentric, telomeric, and the *mat* locus regions. *S. pombe* also shows distinct methylation states but these are only evident upon altering either the Clr4 chromodomain, its SET domain (69), or the bromodomain-containing protein Abo1 (70). Interestingly, the genomes of both *S. pombe* and the Mucorales encode a single Clr4 protein with an N-terminal chromodomain. We hypothesize that *M. lusitanicus* Clr4 is the main histone lysine methyltransferase driving both H3K9me2 and -me3 owing to its essentiality, and suspect that either or both marks have a critical role in cell viability.

An extensive collection of literature endorses the evolution of epigenetic modifications as a defensive mechanism against invasive nucleic acids, particularly transposable elements and repeated DNA ((53, 71) and references therein). Indeed, distinct epigenetic mechanisms for heterochromatin formation have evolved in many fungi to control transposition (71), e.g., CTGS in *S. pombe* and H3K9me and 5mC in *C. neoformans* (20, 21) and in several sordariomycetes where the process is initiated by repeat-induced point mutations (RIP) (72, 73). Interestingly in these organisms, sRNAs are also directed to RIPed sequences but the interaction between RNAi and heterochromatin to regulate repeats remains poorly understood in filamentous fungi, and has only been thoroughly established in the fission yeast.

*M. lusitanicus* deploys an extensive array of RNAi components involved in two competing RNAi pathways. The presence of both active RNAi and H3K9me prompted us to explore interactive dynamics between both mechanisms in this fungus. Our findings indicate that RNAi loss in *M. lusitanicus* does not impair heterochromatin maintenance although sRNAs and H3K9me frequently co-occur in the same regions. This is intriguing because RNAi components have been shown to participate in transcriptional or chromatin-based silencing in other eukaryotic organisms. For example, loss of Ago activity results in defective H3K9me in plant (48), animal (47), and ciliate species (46). Also, in the fission yeast, Ago, Dicer, and RdRP defective mutants are unable to establish or maintain H3K9me across pericentric regions and the mating-type locus (12, 13). Once heterochromatin is established, the RDRC and RITS complexes are directed to H3K9-methylated regions by the chromodomain-containing factor Chp1, generating more sRNAs to sustain a co dependent feedback loop between RNAi and H3K9me (74). In contrast, our findings reveal RNAi is dispensable for maintaining H3K9me across the genome of *M. lusitanicus*, and particularly at pericentric TEs, although RNAi could still be involved in establishing *de novo* H3K9me. This is reinforced by our lethality analysis (*SI Appendix*, Fig. S3), showing that Clr4 activity is essential for cell viability, in contrast to canonical or alternative RNAi pathway mutants which are viable.

Importantly, we identified regions targeted by canonical sRNAs that are devoid of H3K9me and conversely, H3K9-methylated transposable elements that do not trigger RNAi. RNAi independent epigenetic mechanisms have been extensively characterized before in the filamentous fungal model *N. crassa*. Freitag et al. uncovered that RNAi-mediated silencing and DNA methylation were independent genome defense mechanisms in *N. crassa* (75). Simultaneously, Cogoni and Macino showed that H3K9me deposited at RIP targeted transposon relics is not affected in RNAi mutants, although these relics do accumulate canonical sRNAs (76). Later, another study revealed that the RNAi machinery is also dispensable for DNA 5mC co-occurring at these transposon relics (77). Taken together, these findings suggest that RNAi and H3K9me arose as independent regulatory mechanisms that later evolved into co-dependent relationships in some organisms, probably due to their shared targets.

Overall, our work reinforces that epigenetic regulation in fungi is complex and diverse, reflecting a rich evolutionary history of loss and retention of different epigenetic mechanisms. Besides loss of both DNA 5mC and H3K27me-mediated facultative heterochromatin formation machineries, Mucoralean genomes were shaped by loss of two critical kinetochore proteins, CENP-A and CENP-C, resulting in a novel type of mosaic centromere that harbors hybrid features of point and regional centromeric sequences (40). Mosaic pericentromeres are colonized by large, inverted repeats comprising GremLINE1 retrotransposons, and our work has uncovered an exclusive, dual regulation by both H3K9me and canonical RNAi that sets them apart from the remaining transposable elements resident in the genome. H3K9-methylated pericentric regions, spanning up to 70 kilobases, are the largest heterochromatin islands in the genome (Fig. 2A, Fig. 4A, and *SI Appendix*, Fig. S4) but surprisingly, these rely on RNAi and PTGS instead of TGS to repress GremLINE1 expression (Fig. 5). This suggests an alternative regulatory role for H3K9me besides transcriptional silencing and we reasoned an involvement in centromere identity, possibly recruiting the remaining kinetochore proteins in the absence of CENP-A and explaining Clr4 essentiality. Because H3K9me-based heterochromatin by itself has been shown to recruit the kinetochore machinery in *S. pombe* (78, 79), we propose a similar scenario in which heterochromatin-bound GremLINE1 retrotransposons could act as a beacon for the inner kinetochore complex CENP-T-W-S-X (80, 81) or a novel, alternative protein to CENP A. *M. lusitanicus* centrochromatin harbors every element to support this model: large, inverted repeats that trigger H3K9me and a chromodomain-containing Clr4 that can maintain heterochromatin formation even in the absence of RNAi activity. However, there is robust RNAi activity directed against GremLINE1s and though it is dispensable for maintaining H3K9me across these regions, we hypothesized that its role is to restrain the repeats within the bounds of the pericentric regions. The absence of GremLINE1 elements beyond the pericentric regions together with their exceptional, Rdrp2-dependent post-transcriptional regulation strongly supports this hypothesis. Indeed, we have shown that lack of canonical RNAi activity results in GremLINE1 retrotransposition to non-centromeric regions, with multiple implications for *M. lusitanicus* biology. First and foremost, GremLINE1s are strictly controlled by canonical Dicer and Ago RNAi activities. Second, this control is essential to preserve genome stability by preventing transposition to other genomic regions. Third, our results indicate that GremLINE1 retrotransposition may disrupt drug targets to confer antifungal drug resistance, confirming TE insertion as a source of genomic variability that favors adaptation (Fig. 5). The antagonistic interaction between canonical and alternative RNAi pathways in *M. lusitanicus* (36, 57) may fine-tune transposition and act as a fail-safe mechanism, typically ensuring proper Grem-LINE1 repression and genome stability but enabling transposition when beneficial to overcome adverse stressful conditions.

## Materials and methods

### Fungal strains, culturing, and transformation

Every strain used in this work derives from the natural isolate *Mucor lusitanicus* CBS277.49. The parental strain of all the mutant strains is the leucine and uridine double auxotroph MU402 (Ura-, Leu-). This strain was used for *clr4* deletion mutant transformation following a CRISPR-Cas9 procedure as previously described (28, 82). Briefly, we designed two guide RNAs (gRNA) to generate breaks up- and downstream of the reannotated *clr4* locus (gRNA_clr4_5’: CTCCTGGTGACTGGTGAAAGTGG, and gRNA_clr4_3’: GGGTTTTCATTGGCCGTGTCTCC), and a linear DNA construct containing the selectable marker *pyrG* flanked by 1-kb upstream and downstream regions of the gRNA cleavage sites (*SI Appendix*, Table S4). Ribonucleoprotein complexes with Cas9 and gRNAs were assembled in vitro following the supplier instructions (Alt-R™ CRISPR Custom Guide RNAs and Cas9 enzyme, Integrated DNA Technologies). The DNA cassette and the ribonucleoprotein complexes were electroporated into *Mucor* protoplasts. After transformation, colonies were plated on selective minimal medium with casamino acids (MMC) at pH of 3.2 to positively select prototrophic colonies. DNA deletion allele integration and heterokaryosis was assessed by PCR using primers that generate discriminatory amplicons (*SI Appendix*, Table S4), as described previously (28). At least 10 vegetative passages were conducted in an attempt to achieve homokaryotic mutants, consisting of collecting spores from a single asexual sporangium and subsequent plating in MMC medium. Lethality of the *clr4*Δ allele was assessed by dissecting spores from a heterokaryotic mutant onto rich, non-selective YPD medium, and analyzing the *clr4* locus as described above. RNAi mutants analyzed in this study include single Dicer mutant *dcl1*Δ (strain MU406), *dcl2*Δ (MU410), and double mutant *dcl1*Δ *dcl2*Δ (MU411) (31, 32), Argonaute mutants *ago1*Δ, *ago2*Δ, and *ago3*Δ (MU413, MU416 and MU414, respectively) (33), RdRP mutants *rdrp1*Δ, *rdrp2*Δ, and *rdrp3*Δ (MU419, MU420 and MU439, respectively) (29, 57), and the alternative ribonuclease R3B2 mutant *r3b2*Δ (MU412) (35). Media were supplemented with uridine (200 mg/L) or leucine (20 mg/L) when needed to supplement auxotrophic requirements and cultures were grown at room temperature unless otherwise stated.

### Ortholog search and phylogenomic tree inference

Experimentally curated protein sequences were used as queries in a PSI-BLAST v2.12.0 against the publicly available proteomes of 85 fungal species (*SI Appendix*, Dataset S1). Then, BLASTp searches were conducted to retrieve those matches that displayed a positive reciprocal BLAST hit. Protein domains were predicted on those hits using InterProScan v5.59-91.0 and visually examined for manual curation (SI *Appendix*, Dataset S2). BUSCO v5.4.3 identified 389 singlecopy fungal orthologs from fungi_odb10 database that were present in ≥ 90 % of the species. These single-copy gene protein sequences were aligned using MAFFT v7.475, alignments trimmed by TrimAl v1.4.rev15, and used to infer a phylogenomic species tree using IQ-TREE v2.2.0.3.

### Repeats and transposable element prediction

Repeated DNA sequences were predicted using RepeatModeler2 v2.0.3. Then, EMBOSS v6.6.0.0 predicted ORFs across these repeats, and InterProScan identified protein domains. Repeat sequences were manually curated to retain those encoding protein domains frequently associated with TE activity and were annotated according to presence and relative position of those domains. Both raw structural repeat and curated TE libraries were used by RepeatMasker v4.1.3 to produce annotation files.

### Chromatin immunoprecipitation and RNA isolation and sequencing

2.5×10^5^ spore/mL YPD cultures were grown for 16 hours at 26 °C and 250 rpm in duplicates. The same culture was utilized to obtain ChIP and RNA samples to ensure minimal experimental differences. ChIP-grade antibodies α-H3K9me2 (ab1220, Abcam), α-H3K9me3 (ab8898, Abcam), α-RNA pol II (39497, Active Motif) were utilized to immunoprecipitate chromatin-bound DNA, following a previously-established procedure (40). Libraries were prepared using Roche KAPA HyperPrep Kits and sequenced with Illumina NovaSeq 6000 sequencing system for 100-bp paired-end reads. Total RNA was purified with a QIAGEN miRNeasy Mini Kit and RNA samples were divided into small and long RNA preparations. rRNA depleted RNA libraries (long RNA) were prepared using Illumina Stranded Total RNA Prep with Ribo-Zero Gold rRNA Removal Kit and *M. lusitanicus* rDNA specific probes, and cDNA sequenced in a NovaSeq 6000 sequencing system for 150-bp paired-end reads. Lastly, sRNA libraries were amplified using QIAseq miRNA library kit, and sequenced in an Illumina NextSeq 500 High-Output sequencing system to obtain 75-bp single-reads.

### Sequencing data analyses

FASTQ dataset quality was assessed by FASTQC v0.11.9 and reads processed by TrimGalore! V0.6.7 to remove adapters, low quality reads and adjust minimal length for subsequent analyses. Reads were aligned to the *M. lusitanicus* MU402 genome (https://mycocosm.jgi.doe.gov/Muccir1_3/Muccir1_3.info.html) employing BWA-MEM v.0.7.17 for ChIP DNA, STAR v.2.7.10a for long RNA, and ShortStack v3.8.5 for sRNA reads. ChIP enrichment was determined as IP/Input DNA ratio of counts per million (CPM) reads. Broad ChIP-enriched regions were identified by MACS2 v2.2.7.1. Long RNA data was quantified by featureCounts v2.0.1 from the Subread package, modifying strandness to the forward or reverse strand as needed to generate sense or antisense counts, respectively. Log_2_ CPM values were determined using the limma package v3.50.3, normalizing by the trimmed mean of M-values (TMM) method. Pairwise comparisons and log_2_ fold change values were determined and tested for significance (FDR test corrected by the Benjamini-Hochberg). sRNA aligned reads were deduplicated by UMI-tools v1.1.2. Reads aligning to tRNA or rRNA sequences were removed, and the remaining genome-aligning reads were quantified by ShortStack and normalized to reads per million (RPM). Coverage files for ChIP DNA and RNA alignments were generated by Deeptools2 v3.5.1.

### Artificial evolution experiment and isolation of spontaneous FK506 resistant isolates

RNAi mutant and parental MU402 strains were cultured during 20 vegetative passages by collecting spores from single colonies and plating them in MMC at pH of 4.5. After 20 passages, spores were collected, counted, and adjusted to the same concentration to plate them on YPD supplemented with FK506 (1 mg/L). After 3-7 days incubation, colonies growing mycelial sectors were selected and plated in YPD supplemented with FK506 and rapamycin (0.1 mg/L) to confirm resistance was caused by FKBP12 loss-of-function. Genomic DNA was purified using MasterPure™ Complete DNA and RNA Purification Kit (Lucigen). The whole *fkbA* locus from the FK506 resistant isolates was amplified using primers JOHE52223 and JOHE52224 and LA Taq DNA polymerase for long-range PCR with GC Buffer I (Takara). PCR conditions were optimized for long fragments as follows: 94°C 1min, 30 cycles: 94°C 5s, 55°C 30s, 72°C 12min; and 72°C 10min. PCR fragments were Sanger-sequenced using primers listed in *SI Appendix*, Table S4. A primer walking strategy was designed to sequence the full-length insertions. Their full-sequence was used as query for BLASTp searches against a nucleotide database containing every TE element previously identified, ensuring alignment length covered ≥ 97 % of the query length (qcov_hsp_perc 97), i.e., the full insertion sequence except for possible mismatches at either 5’- or 3’-end.

### Quantitative PCR

Genomic DNA was purified using MasterPure™ Complete DNA and RNA Purification Kit (Lucigen). Quantitative PCR (qPCR) was performed using diluted genomic DNA from each sample, SYBR Green PCR Master Mix (Applied Biosystems™) and primers that specifically amplified a TE subfamily or the single-copy gene *vma1* (*SI Appendix*, Table S4). qPCR reactions were performed in triplicate using a QuantStudio™ 3 Real-Time PCR System. TE Ct values were normalized to *vma1* using the 2^-ΔCT^ method to estimate copy numbers.

## Data availability and supporting information

Raw sequencing datasets will be available upon publication under the following NCBI’s Sequence Read Archive project accession number: PRJNA903107. A Supporting Information (SI) Appendix file contains a detailed method explanation and supplementary figures and tables. Supplementary datasets, figures, and tables can also be accessed through the following Mendeley Data repository: Supplementary material.

## ACKNOWLEDGEMENTS

We thank Drs. Minou Nowrousian, Patricia Peterson, and Sheng Sun for critical reading and insightful suggestions on the manuscript. Also, thanks to our laboratory manager Anna Floyd Averette for her constant support. We extend our gratitude to Drs. Rosa M. Ruiz-Vázquez, Santiago Torres-Martínez, and Francisco E. Nicolás for providing fungal strains. We appreciate Drs. Kaustuv Sanyal, Victoriano Garre, and their teams for their input and discussion sessions on fungal epigenetics and centromere biology. Particularly, we thank Dr. Shweta Panchal for her counsel on H3K9me2 antibodies for chromatin immunoprecipitation. We are also grateful to Drs. Zanetta Chang and Silvia Calo for their recommendations on isolating FKBP12 loss-of-function insertional mutants in *M. lusitanicus.* We commend the Duke Computer Cluster for their computing equipment and Duke’s Sequencing and Genomic Technologies Core Facility for their assistance, especially its Associate Director Dr. Devjanee Swain Lenz for her sage advice and expertise. Also, special thanks to Dr. Fred Dietrich for his invaluable computational resources and dedication. This study was supported by NIH/National Institute of Allergy and Infectious Diseases R01 Grant R01 AI170543 awarded to J.H. J.H. is Co-Director and Fellow of the CIFAR program Fungal Kingdom: Threats & Opportunities.

