## Supporting Information Appendix for "Heterochromatin and RNAi act independently to ensure genome stability in Mucorales human fungal pathogens"

[2]Equally-contributing authors

*To whom correspondence may be addressed

Corresponding authors: Joseph Heitman, María Isabel Navarro-Mendoza, and Carlos Pérez-Arques. 322 Research Drive, CARL Building, Box 3546. Durham, NC 27710. USA. Telephone: 919-684-2824

**This PDF file includes:**

Supporting information text

Extended Materials and Methods

Figures S1 to S10

Tables S1 to S4

Legends for datasets S1 to S4

SI References

**Other supporting materials for this manuscript include the following:**

Datasets S1 to S4

**SI Appendix**

**Supporting information text**

**Extended Materials and Methods**

**Ortholog search and phylogenomic tree inference**

Experimentally curated KMT, DNMT, KDM, and accessory proteins multi-FASTA datasets were used as queries for a PSI-BLAST v2.12.0 search (83) against a proteomic database comprised of 86 fungal species (84-117), knowingly intending to oversample Mucoromycota and fungal pathogen species but also selecting at least one representative of each fungal phylum if genomes were publicly available (see *SI Appendix*, Dataset S1 for a comprehensive list of query protein sequences and species analyzed). PSI-BLAST searches were iterated for three consecutive runs, retrieving matches with E-values ≤ 0.001. Then, every match was subjected to a reciprocal BLASTp search against a database comprising the whole proteomes of the same species that were utilized as queries. From all of the initial matches, only those that satisfied a positive reciprocal BLASTp match were regarded as putative homologs (protein FASTA sequences are contained in Dataset S4). Non-reciprocal matches were examined manually to assess the efficacy of our strategy and correct any possible error in our workflows or protein databases. InterPro (IPR) protein domains from CDD, Gene3D, HAMAP, PANTHER, Pfam, PIRSF, PRINTS, PROSITE, SFLD, SMART, SUPERFAMILY, and TIGFRAM databases were predicted on those protein matches sequences deemed putative homologs using InterProScan v5.59-91.0 (118). Protein predicted domains and full-length sequences were plotted as rectangle diagrams for visual evaluation. Briefly, every database protein domain that matched the same IPR entry was collapsed to show only unique IPR domains, and these were color-coded, scaled, and plotted for every chromatin remodeler or accessory protein analyzed (see *SI Appendix*, Dataset S2). Hierarchically redundant IPR entries, e.g., super-, sub- and families, were also collapsed, showing those that bore the maximum number of hits in the datasets. Species were sorted according to phylogenies shown on Fig. 1 and *SI Appendix* Fig. S2 to facilitate visual evaluation, and those phylogenies were generated as follows. BUSCO v5.4.3 fungal OrthoDB v10 dataset "fungi_odb10" (119) was employed to conduct orthology searches in a subset of 65 species' proteomes to avoid redundancy, selecting a total of 389 single-copy orthologs that were present in ≥ 90 % of the species and in *Rozella allomycis,* that was used as the outgroup for further phylogenetic analyses. Protein sequences of those single-copy orthologs in every species were aligned by MAFFT v7.475 (120), automatically selecting an appropriate alignment strategy, using the Smith-Waterman algorithm (--localpair) and a maximum of 1,000 iterations. Protein alignments were trimmed and gaps removed by TrimAl v1.4.rev15 (121) and the gappyout method, and trimmed alignments were utilized as partitions by IQ-TREE v2.2.0.3 (122) to infer a phylogenetic tree with 1,000 ultrafast bootstraps and SH-aLRT replicates, automatically detected best fitted protein substitution model, and setting *R. allomycis* as outgroup. In addition, 42108 was used as seed for reproducibility.

**Repeats and transposable element prediction**

Structural repeats and putative TEs were predicted by RepeatModeler2 v2.0.3 (123) allowing for LTR structural identification (-LTRstruct). This raw repeat library was merged with the RepBase database "RepBaseRepeatMaskerEdition-20181026" (124) to annotate every possible repeat in the genome using RepeatMasker v4.1.3 (http://www.repeatmasker.org/) slow search (-s), skipping bacterial insertion element check (-no_is). The subsequent GFF file was regarded as a raw structural repeat annotation file, but additional effort was directed to generating a curated TE library and annotation file. First, ORFs longer than 300 bp were predicted across the sequences contained in the RepeatModeler2 library using EMBOSS v6.6.0.0 getorf (125). Then, InterProScan was utilized to predict protein domains encoded by this ORFs. Each full-length element together with their ORFs and protein domain were scaled, color-coded, and plotted to allow visual inspection as described above. Briefly, repeats encoding protein domains frequently associated with TE activity (e.g., reverse transcriptase, transposase, RNase, nucleic acid binding) were retained, and the domain type and relative position within each sequence was used to manually curate bona fide TE sequences, particularly the position of an Integrase-related protein domain before or after the reverse transcriptase and RNase tandem to discriminate between Ty1/Copia or Ty3/Gypsy LTR retrotransposons, respectively (126). Lastly, this curated TE library was used with RepeatMasker to annotate every TE copy with the same parameters as above, but also skipping simple repeats (-nolow), small RNA genes (-norna), and ensuring a stringent cut-off value of 5,000 (-cutoff 5000) to obtain mostly full-length elements to generate a curated TE annotation file.

**ChIP and RNA isolation and sequencing**

$2.5x10^7$ spores from each RNAi mutant and MU402 wild-type strains were cultured in 100 mL of YPD medium for 16 hours with constant shaking (250 rpm) at 26 ºC. Two biological replicates were harvested and processed for each sample. Samples were sonicated using a Bioruptor (Diogenode) with pulses of 30 s ON/OFF for 35 cycles to obtain a sheared chromatin of 100-300 bp, then they were divided for antibody IP and input DNA control. For ChIP samples, anti-Histone H3 (di methyl K9) ChIP Grade ab1220 (Abcam), anti-Histone H3 (tri methyl K9) ChIP Grade ab8898 (Abcam), and RNA pol II antibody 39497 (Active Motif) antibodies were used to immunoprecipitated the DNA. According to their isotypes, primary ChIP antibodies were conjugated to protein A or G magnetic beads before DNA purification. Samples were washed and de-cross-linked following a previously established procedure (40), and DNA was purified and eluted using a phenol:chloroform extraction method. IP and Input DNA samples were used to prepare libraries with Roche KAPA HyperPrep Kit, loaded into an Illumina NovaSeq 6000 S Prime flow cell and sequenced to obtain 100-bp paired-end reads. For total RNA purification, mycelial samples were purified with a miRNeasy Mini Kit (QIAGEN) following the procedure for total RNA purification supplied by the manufacturer. The same total RNA sample was used to generate both small and long RNA libraries and subsequent sequencing, aiming to minimize biological differences between small and long RNA datasets from the same strain. Long RNA libraries were prepared by an rRNA removal method, using Illumina Stranded Total RNA Prep with Ribo-Zero Gold rRNA Removal Kit. This method allowed using customized *M. lusitanicus* 28S, 18S, and 5.8S rDNA probes, predicted by Barrnap v0.9. Libraries were loaded into an Illumina NovaSeq 6000 S Prime flow cell and sequenced for 150-bp paired-end reads. sRNA libraries were amplified using QIAseq miRNA library kit, loaded into a High-Output flow cell, and sequenced in an Illumina NextSeq 500 sequencing system to obtain 75-bp single-reads.

**ChIP-seq data analysis**

NovaSeq Control Software v1.7.5 generated one FASTQ dataset per flow-cell lane and sample that were concatenated if needed into a single dataset per sample. Dataset quality was assessed by FastQC v0.11.9 (https://www.bioinformatics.babraham.ac.uk/projects/fastqc/) and results were jointly evaluated after being processed by MultiQC v1.12 (127). Reads were processed by Trim Galore! v0.6.7 powered by Cutadapt v1.18 (128) to remove adapters, low quality reads (--quality 20), and reads shorter than 50 nt (--length 50). Processed reads were aligned using the BWA-MEM v.0.7.17 (arXiv:1303.3997) algorithm for medium and long reads to the *M. lusitanicus* MU402 genome (https://mycocosm.jgi.doe.gov/Muccir1_3/Muccir1_3.info.html), set at default parameters, and alignment files sorted with Samtools v1.3.1 (129). Coverage bigWig files for H3K9me2, -me3, and RNA pol II were generated from alignments using Deeptools2 v3.5.1 (130) bamCoverage for IP and Input DNA separately and bamCompare to assess the ratio IP/Input, normalizing using CPM and a binsize of 25 nucleotides. H3K9me2 and -me3 enrichment values in 10-kb windows across the whole genome were determined using Deeptools2 multiBigwigSummary. To identify broad enriched regions between each IP/Input pair of H3K9me2 or -me3, MACS2 v2.2.7.1 predictd (131) was used to predict fragment length, and this value fed into callpeak, keeping duplicates and setting an effective genome size as precalculated by Kent's utils v438 faCount (132). Customized BED files containing genes (coding sequence including introns), repeated, and intergenic sequences were extracted from a publicly available GFF3 annotation file and the previously generated repeat file using BEDtools v2.27.1 utilities sort and complement (133). H3K9me2 and -me3 broad peaks in the wild-type strain were intersected using BEDtools intersect and the resulting overlapping H3K9me2/me3 peaks were mapped to gene, repeat, and intergenic features to calculate the number of H3K9me base pairs in each feature using BEDtools intersect and Bioawk v20110810 (https://github.com/lh3/bioawk/).

**Long RNA-seq data analysis**

Initial raw read processing was conducted as above, and processed reads were mapped to the genome using RNA-seq aligner STAR v.2.7.10a (134), ensuring all best scoring alignments were reported as primary alignments up to 500 alignments, and a predicted intron size ranging from 10 to 1000 bases. Alignments were sorted and classified into forward and reverse stranded by Samtools sort, view, and merge according to their flag values as follows: forward alignments included first-pair alignments from the reverse strand (-f 64 and 16) and second-pair alignments excluding those from the reverse strand (-f 128 and -F 16), and reverse alignments included second-pair alignments from the reverse strand (-f 128 and 16) and first-pair alignments excluding those from the reverse strand (-f 64 and -F 16). Then, bamCoverage bigWig files were generated as described above. Counts per transcript were quantified using featureCounts v2.0.1 (135) from the Subread package and a customized annotation file including gene annotation from JGI and the newly identified TEs. Briefly, primary alignments from read pairs were mapped to exon features in the annotation file, ignoring chimeric alignments, and assigning a fraction of counts in case of multimapping alignments or alignments overlapping two or more features to ensure transposable elements were properly quantified. In addition, strandedness was modified to forward or reverse strand as needed to generate sense or antisense counts, respectively. A countmatrix containing all samples was utilized by the limma package v3.50.3 (136) to determine counts per million (CPM) values across genes and TEs, normalizing values by the trimmed mean of M-values (TMM) method. $Log_2$ fold change values were determined and a False Discovery Rate (FDR) test corrected by the Benjamini-Hochberg method was performed in pairwise sample comparisons across all genes and TEs.

**sRNA-seq data analysis**

Initial raw read processing was conducted as above, with some exceptions. NextSeq Control Software v4.0.1.41 was used for base-calling. In addition, 12-base pair Unique Molecular Identifiers (UMIs) were detected and printed into each FASTQ sequence header by UMI-tools v1.1.2 (137), discarding QIAGEN 3'-adapters (5'-AACTGTAGGCACCATCAAT-3'). After that, reads were further processed by Trim Galore! to remove Illumina PCR 3'-adapters, short and low-quality reads (--length 10 --quality 20), with unusually low stringency parameters (--stringency 4) to avoid trimming real bases from the 3'-end which would bias the sRNA length analysis, i.e., adapter sequences at 3-end were removed only if 4 or more bases matched an adapter sequence. Default stringency parameters (--stringency 1) would trim the 3'-end if only 1 base matched an adapter sequence, resulting in a random 1-bp 3-trimming of ~25% reads. After that, processed reads were aligned to tRNA, rRNA, and whole genome sequences to generate quantitative statistics, concatenating replicates from the same strain and also, separately. In more detail, reads were aligned to the genome by Bowtie v1.3.1 (138), ensuring the best possible alignment and no mismatches (-v 0 --best). Then, aligned reads were deduplicated by UMI-tools to remove possible contaminants, tRNA and rRNA sequences were predicted using tRNAscan-SE v2.0.9 (139) and Barrnap v0.9 (https://github.com/tseemann/barrnap/), respectively. Reads were aligned separately to tRNA and rRNA sequences using identical parameters as above. All of these alignments were counted using Samtools v1.3.1 and Bioawk. Aligned reads mapping to the genome but not to tRNA or rRNA sequences were used for further analyses. Read length and the presence of uracil/thymine as their first, 5’-nucleotide was assessed using Samtools and Bioawk. Aligned reads mapping to intronic, exonic, repeated, and intergenic features (see above) were counted using BEDtools coverage. Then, sRNA content across genes and transposable elements was quantified using ShortStack v3.8.5 (140) and a manually generated ShortStack annotation file comprising JGI gene annotation and our newly identified transposable elements. ShortStack realigned the reads allowing for multi-mapping alignment discovery, powered by Bowtie and reporting every possible alignment lacking mismatches (in short, ShortStack parameters --mismatches 0 --bowtie_m 'all'; that result in Bowtie parameters -v 0 -a). Multi-mapping alignments were placed as a fraction of all mapped reads (--mmap f), and siRNA predicted sizes were set to 21-25 nucleotides (--dicermin 21 --dicermax 25). bamCoverage files were generated as described above.

**Data statistical analyses and visualization**

Cartesian, two-dimensional graphs were plotted using R ggplot2 v3.3.6 (141) and ggpubr v0.4.0 (https://github.com/kassambara/ggpubr/) packages. One-way ANOVA and Tukey’s Honestly Significant Difference (HSD) testing was performed using the aov and TukeyHSD functions, respectively, and pairwise comparisons summarized in a compacted letter display employing the multcomp v1.4-20 package (142). Pearson’s correlation coefficients and their assigned *p*-values, regression linear models, were computed and plotted using the ggpubr stat_cor function, and regression linear models with ggplot2 geom_smooth function. Genomic plots including ChIP enrichment, sRNA, and mRNA coverage and genomic feature data were rendered by Deeptools2 pyGenomeTracks v3.7 (143) utilizing customized configuration files, coverage files, and a BED12 annotation file comprising gene, repeated, and transposable element features. ChIP enrichment heatmaps across full-length sequences were plotted by Deeptools2 plotHeatmap, as well as enrichment profiles using plotProfile to display average values as the mean and standard deviation. Small and long RNA normalized or $\log_2$ fold change heatmaps were plotted using the pheatmap package v1.0-12 (https://github.com/raivokolde/pheatmap/). Single-copy ortholog inferred phylogenomic trees were plotted by the ape v5.6-2 package (144), rotating branches and pruning leaves for visual effects, and using true branch lengths.

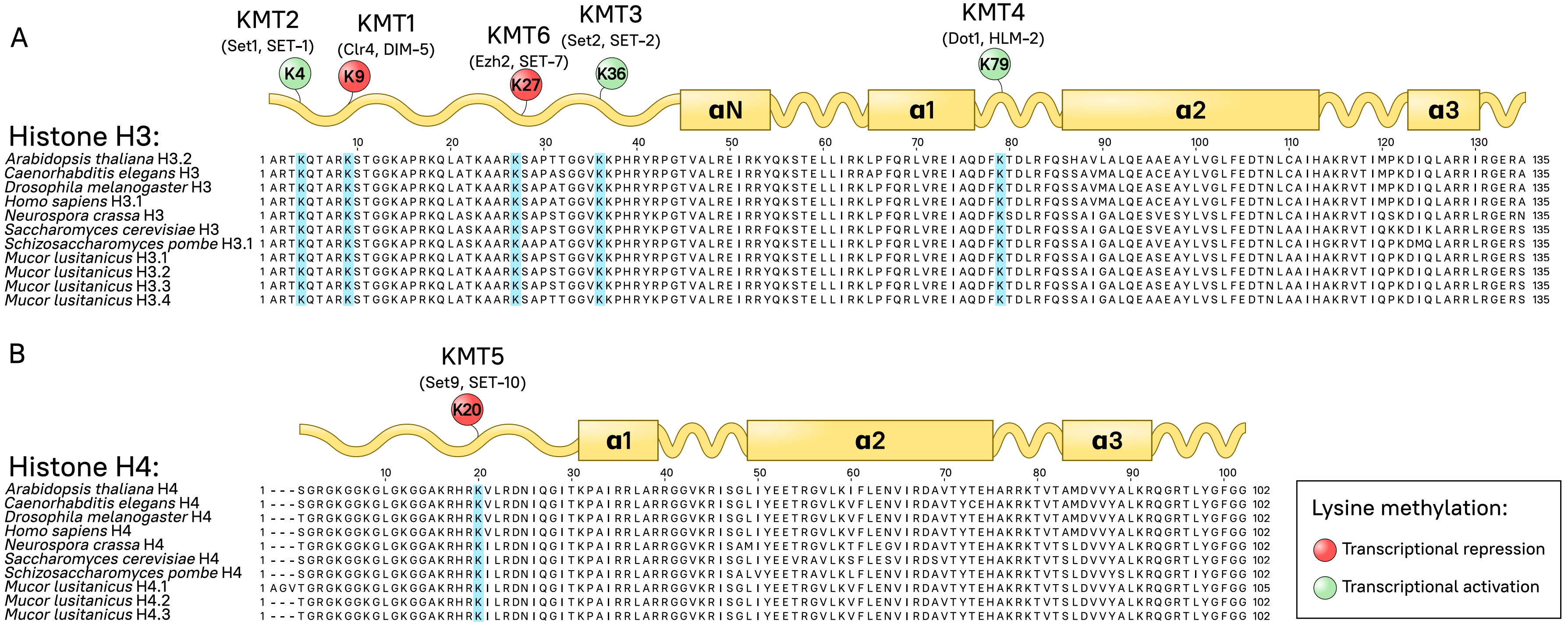


**Fig. S1.** *Mucor* histones display conserved residues for histone-lysine methyltransferase (KMT) activity. Histone H3 **(A)** and H4 **(B)** protein sequence alignment of several model organisms and *M. lusitanicus* is shown. Key lysine residues are highlighted in the alignment (cyan). Methyltransferases involved in histone-lysine methylation and the residue they target are depicted in a schematic representation of each histone protein, showing the N-terminal tail and the histone fold domain comprised of several α-helices (rectangles). Transcriptional repressive (red) and activating (green) modifications are displayed.

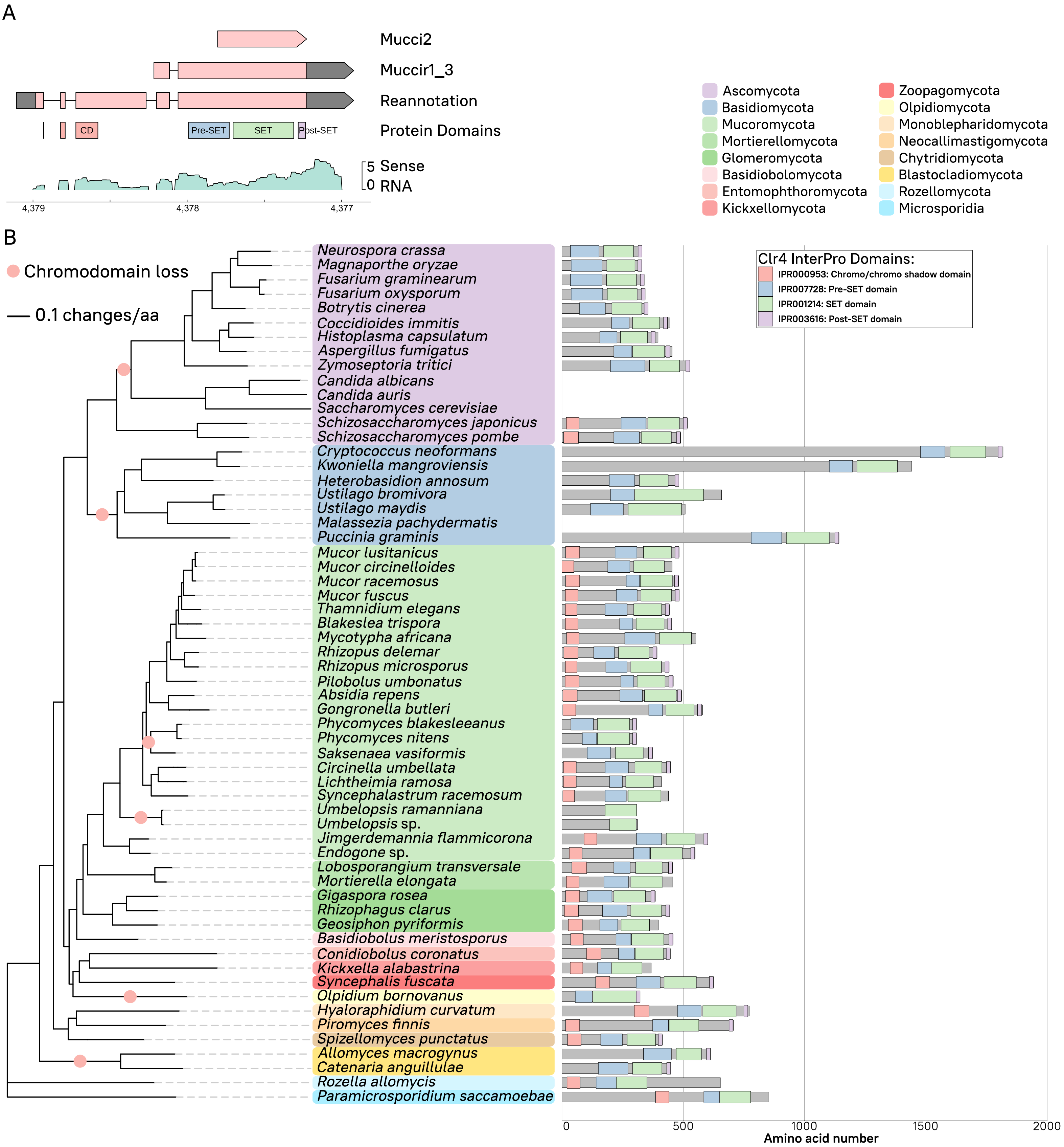


**Fig. S2**. Mucoromycota and other early-diverging fungi harbor an N-terminal chromodomain in Clr4. **(A)** Genomic plot of the *clr4* locus in *M. lusitanicus* MU402. Three different *clr4* gene models are shown (red arrowed blocks, dark gray for UTR regions), including the last reannotation according to our transcriptomic data. Position of Clr4 protein domains encoded in the DNA sequence is shown below gene models for reference. Sense transcripts are plotted as normalized $\log_2$ counts per million (CPM) values (green). **(B)** Clr4 methyltransferases of 60 fungal species clustered by phylogenomic relationship (left, tree), color-coded by different fungal phyla. The full-length, scaled protein sequence is shown along its predicted protein domains (legend). Chromodomain loss is pointed as a red circle in the phylogenomic tree. Phylogenomic distance (black solid line) is shown as changes per amino acid.

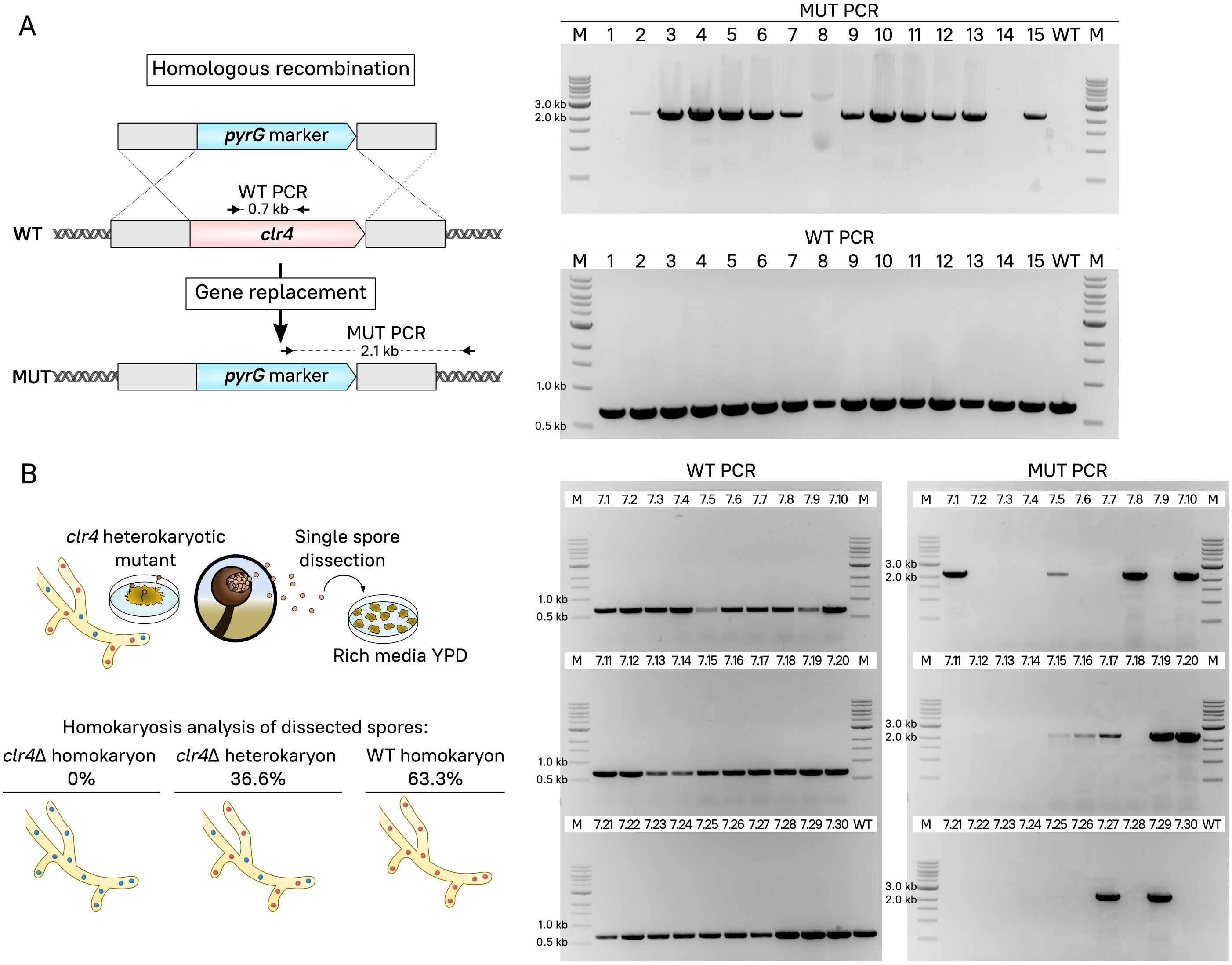


**Fig. S3.** *clr4* homokaryotic deletion results in lethality. **(A)** Schematic strategy followed for *clr4* gene deletion, replacing it with the *pyrG* marker. Primer binding sites for wild-type allele (WT PCR) and 3' mutant allele junction (MUT PCR) are shown. Gel electrophoresis showing PCR products from WT and MUT PCR in 15 transformants (1-15) and wild-type strain (WT). **(B)** Procedure to obtain homokaryotic mutants by single spore dissection from the heterokaryotic mutant #7. WT and MUT PCRs of 30 dissected spores (from 7.1 to 7.30). Spore genotype (mutant homokaryon, heterokaryon, and wild-type homokaryon) is summarized as percentage of total spores. M indicates DNA ladder in electrophoresis gels.

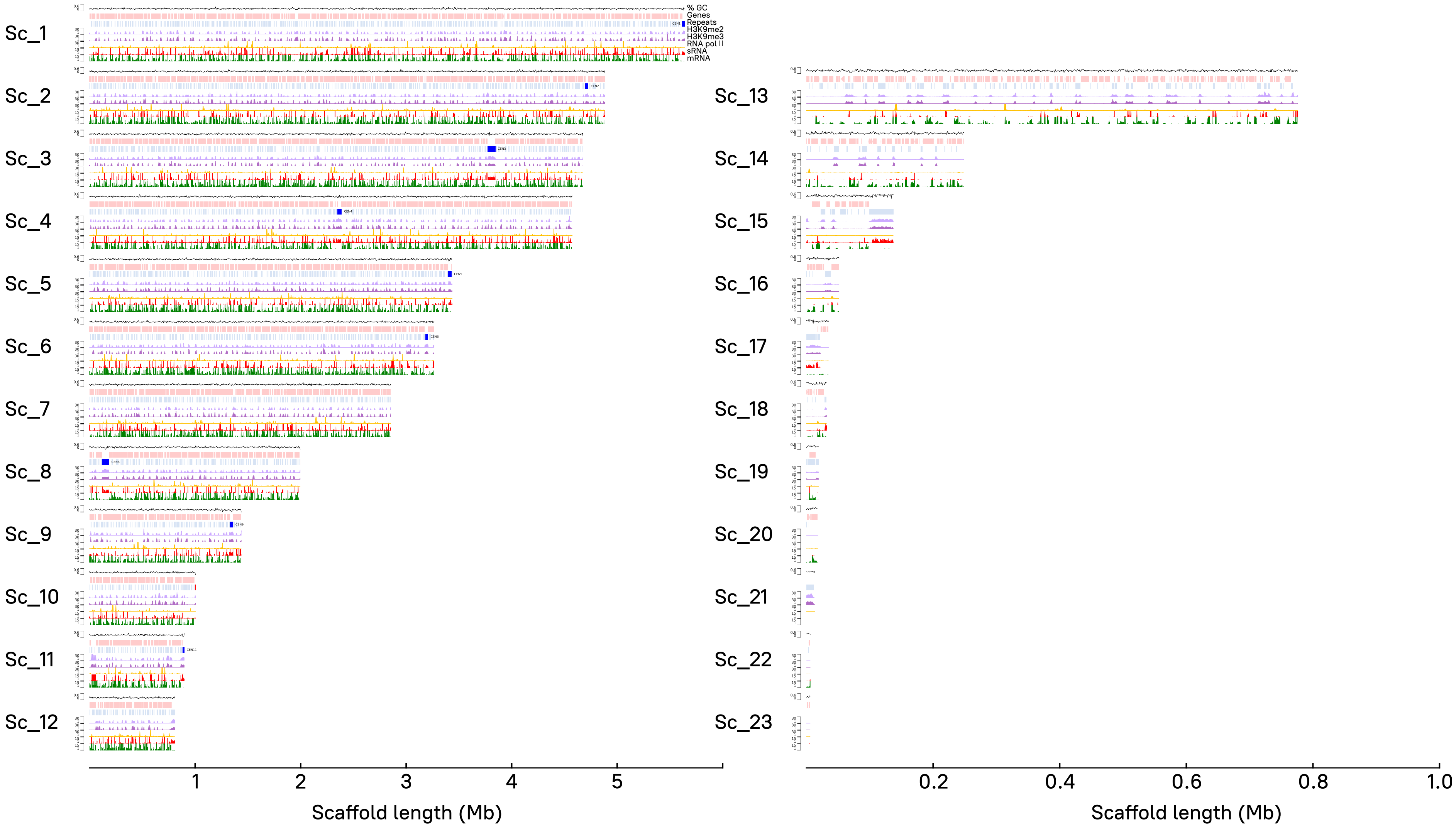


**Fig. S4.** H3K9me2, -me3, RNA pol **II**, sRNA, and mRNA genome-wide distribution in *M. lusitanicus*. Read coverage normalized as $log_2$ CPM is shown as immunoprecipitated (IP) DNA/Input DNA ratio for H3K9me2 (light purple) and -me3 (purple) and RNA polymerase **II** (RNA pol **II**, gold); messenger RNA (mRNA, green) and sRNA (red) coverage as $log_2$ CPM. Genes (light red) and repeated sequences (light blue) including pericentric regions (bright blue) are displayed, as well as GC content coverage as a percentage (black line). Every scaffold (1-23) and its length are plotted, but note that larger scaffolds comprising ≥ 95 % of the genome (1-12, left) are scaled differently than shorter scaffolds (13-23, right).

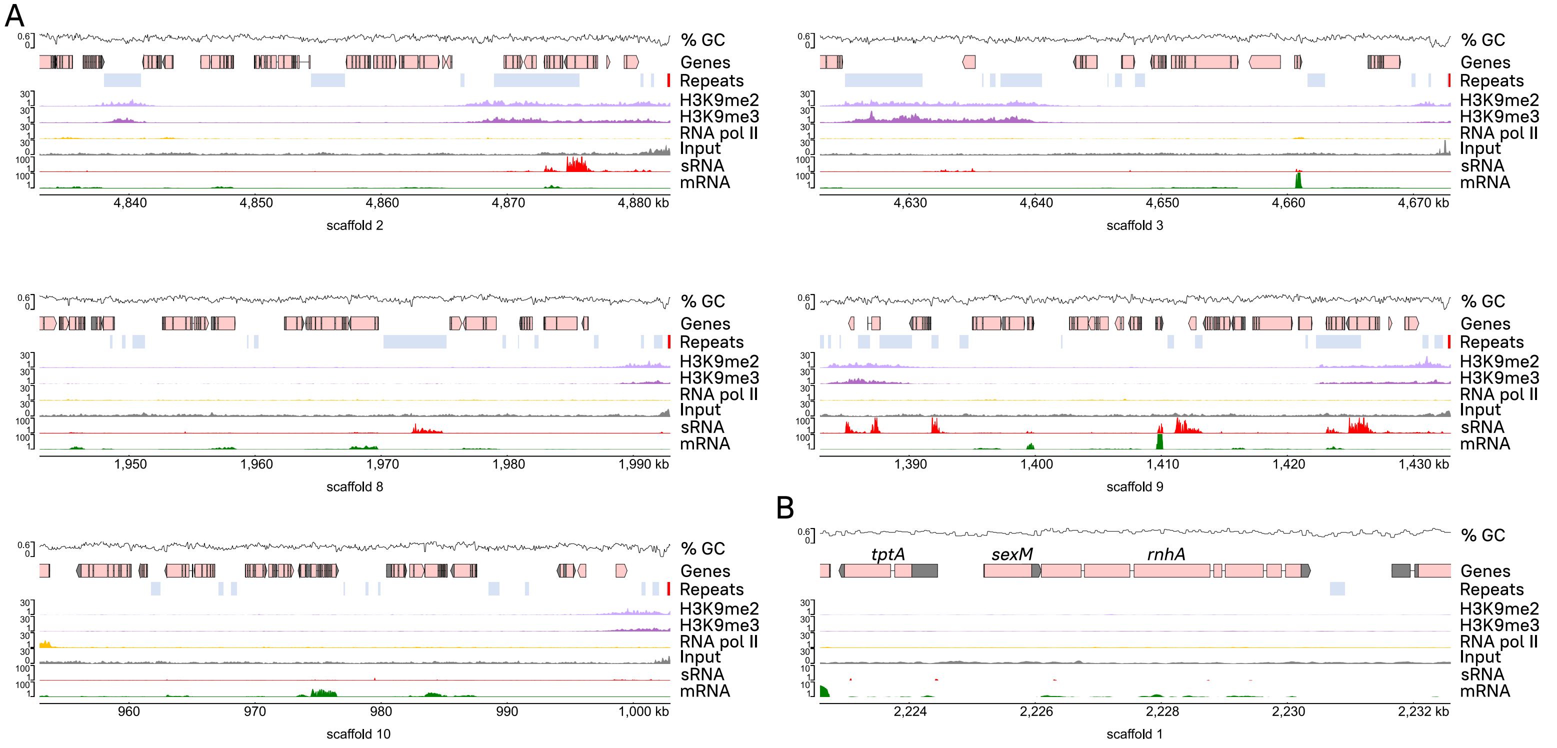


**Fig. S5**. H3K9me and sRNA enrichment in telomeric and sex-determining regions. H3K9me2 (light purple), -me3 (dark purple), and RNA polymerase II (RNA pol II, gold) IP/Input ratio, as well as Input control coverage for reference, are shown across 50-kb telomeric regions identified **(A)** and the *sex* locus **(B)**, and normalized as previously described. Transcripts (mRNA, green) and sRNA (red) coverages are also depicted. Genes (light red) and repeated sequences (light blue) including telomeric repeats (bright red) are displayed, as well as GC content coverage as a percentage.

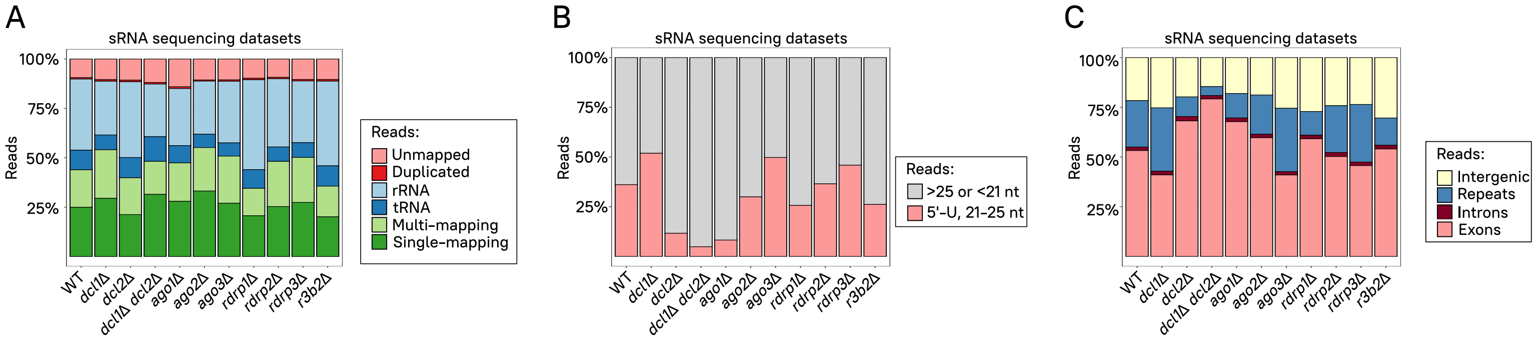


**Fig. S6.** Canonical sRNAs targeting primarily exonic sequences are generated by Dicer and Argonaute activity. **(A)** The percentage of unmapped, duplicated, rRNA or tRNA mapping, multi- or single-mapping reads to the genome in RNAi mutant and wild-type strain sRNA samples is depicted. **(B)** The percentage of aligned reads with a size ranging from 21 to 25 and bearing a 5'-uracil in the same samples is shown. **(C)** The percentage of aligned reads mapping to intergenic, repeated, intronic, and exonic features in the same samples is plotted.

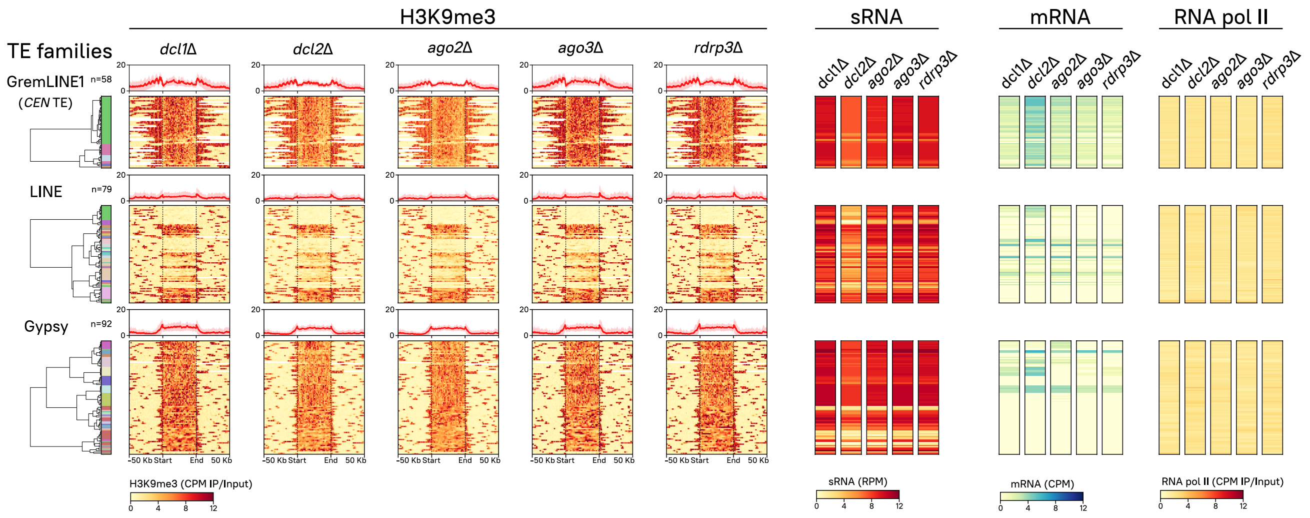


**Fig. S7**. H3K9me3, sRNA, mRNA, and RNA pol II enrichment across RNA transposons in non-essential RNAi paralog mutants. GremLINE1, other LINE, and Gypsy RNA transposon copies are clustered as in Fig. 3. Heatmaps depicting H3K9me3, sRNA, mRNA, and RNA polymerase II values in the wildtype and mutants in non-essential RNAi enzymes are plotted. H3K9me3 enrichment values are shown as IP/input DNA ratio of normalized $\log_2$ CPM from start to end of each copy (divided in 250 bins) and 50 kb upstream and downstream; as well as the mean value of all the copies in each family shown in a profile plot on top of each heatmap. sRNA and mRNA values are normalized to $\log_2$ reads per million (RPM) and CPM, respectively. RNA pol II enrichment values are also displayed as IP/Input ratio of normalized $\log_2$ CPM.

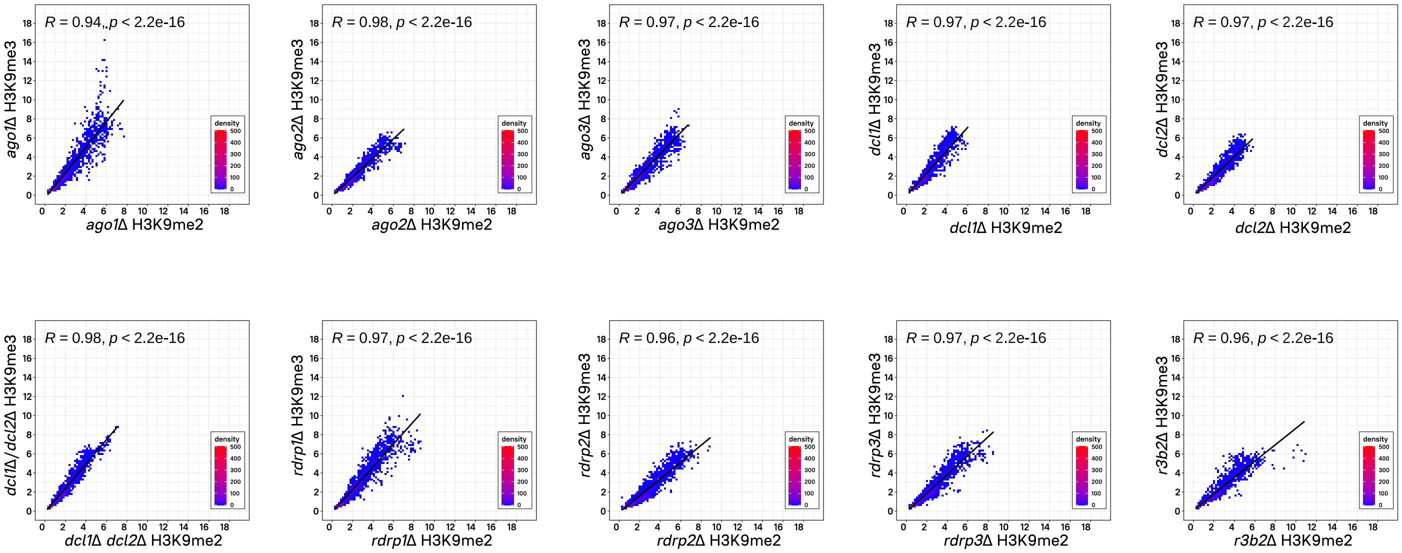


**Fig. S8.** H3K9me2 and -me3 distribution is significantly correlated in RNAi-deficient mutants. Pearson's correlation coefficient (R, dark line) and its *p*-value shown in a dot plot of H3K9me3 (y-axis) and -me2 (x-axis) average $\log_2$ CPM enrichment values in 10-kb regions representing the whole genome of *M. lusitanicus*. Overlapping dots are shown as blue-to-red colored corresponding to plotting density.

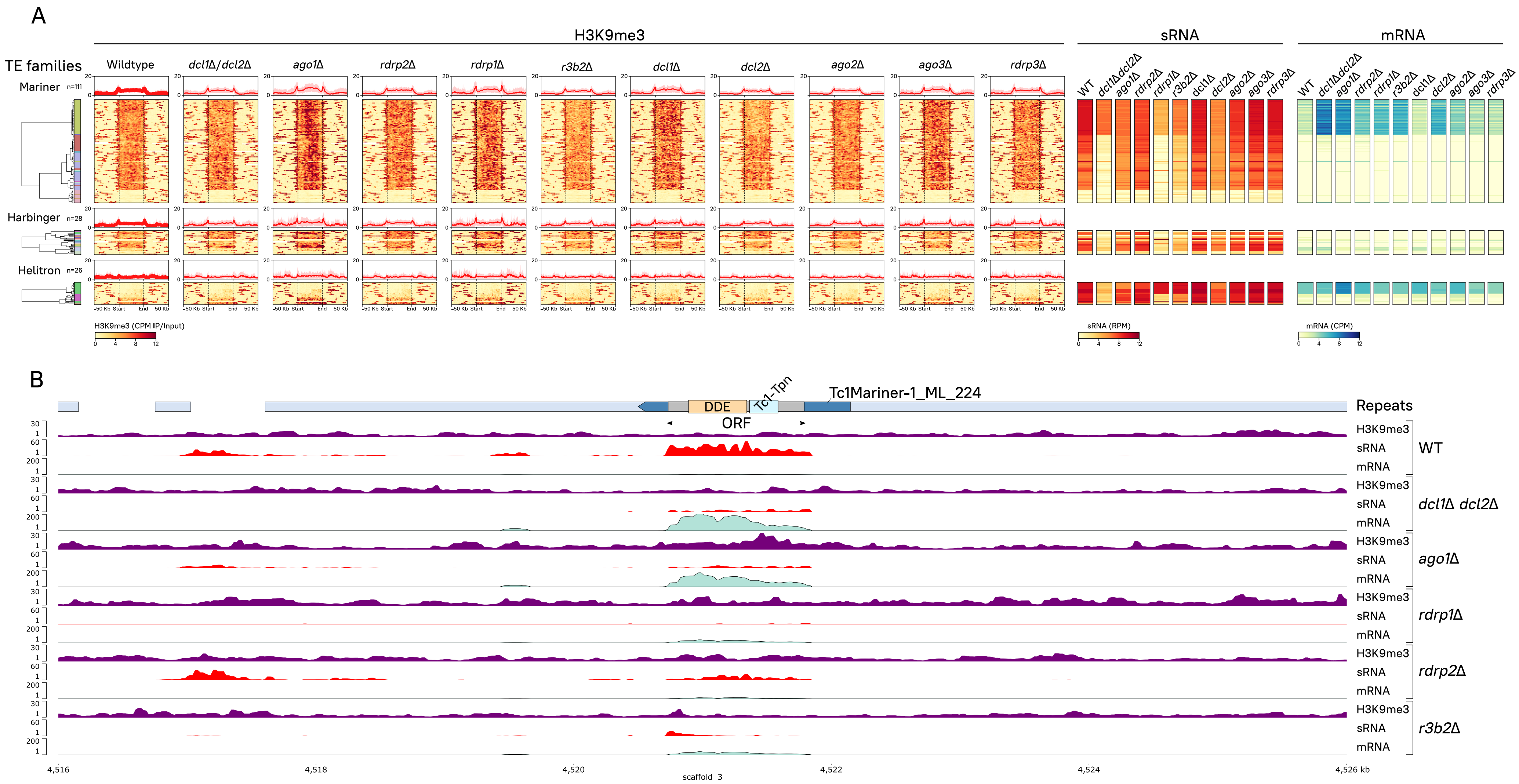


**Fig. S9.** DNA transposons are also targeted by both RNAi and H3K9me-based heterochromatin. **(A)** Every copy of Tc1/Mariner (n=111), PIF/Harbinger LINE (n=28), and RC/Helitron (n=26) DNA or Class II transposons identified in the genome are clustered according to their subfamily, H3K9me3, sRNA, and mRNA values in the wild-type *M. lusitanicus* strain. Heatmaps depicting H3K9me3, sRNA, mRNA, and RNA polymerase II values in wild-type and all RNAi mutant strains –essential and non-essential for RNAi– are plotted (see plot name above). As in Fig. 3, each row represents a single DNA transposon copy, and data values are aligned across all heatmaps to facilitate comparison. From left to right, a color-coded dendrogram indicating each element subfamily. Next, H3K9me3 enrichment values are shown as IP/input ratio of $\log_2$ CPM, from the start to the end of each copy and 50-kb upstream and downstream regions. The average of all the copies in each family is plotted in an H3K9me3 profile on top of each heatmap (bold red line shows the average and faded red background shows the standard deviation). Next heatmap to the right shows sRNA values normalized to $\log_2$ RPM, followed by a sense transcript heatmap (mRNA) in $\log_2$ CPM. The last heatmap on the right depicts RNA pol II enrichment values as IP/Input ratio of $\log_2$ CPM. **(B)** Tc1/Mariner-1 (blue arrowed block) representative genomic plot showing H3K9me3, sRNA, and mRNA data normalized as in **(A)** for canonical (*dcl1*Δ *dcl2*Δ, *ago1*Δ, *rdrp2*Δ, and *rdrp1*Δ) and alternative (*rdrp1*Δ and *r3b2*Δ) RNAi-deficient mutants. Transposition-related protein domains (colored rectangles) encoded by Tc1/Mariner-1_ML_224 ORF sequence (gray rectangle) and their position is shown. Other structural repeats (light blue rectangles) are plotted for reference.

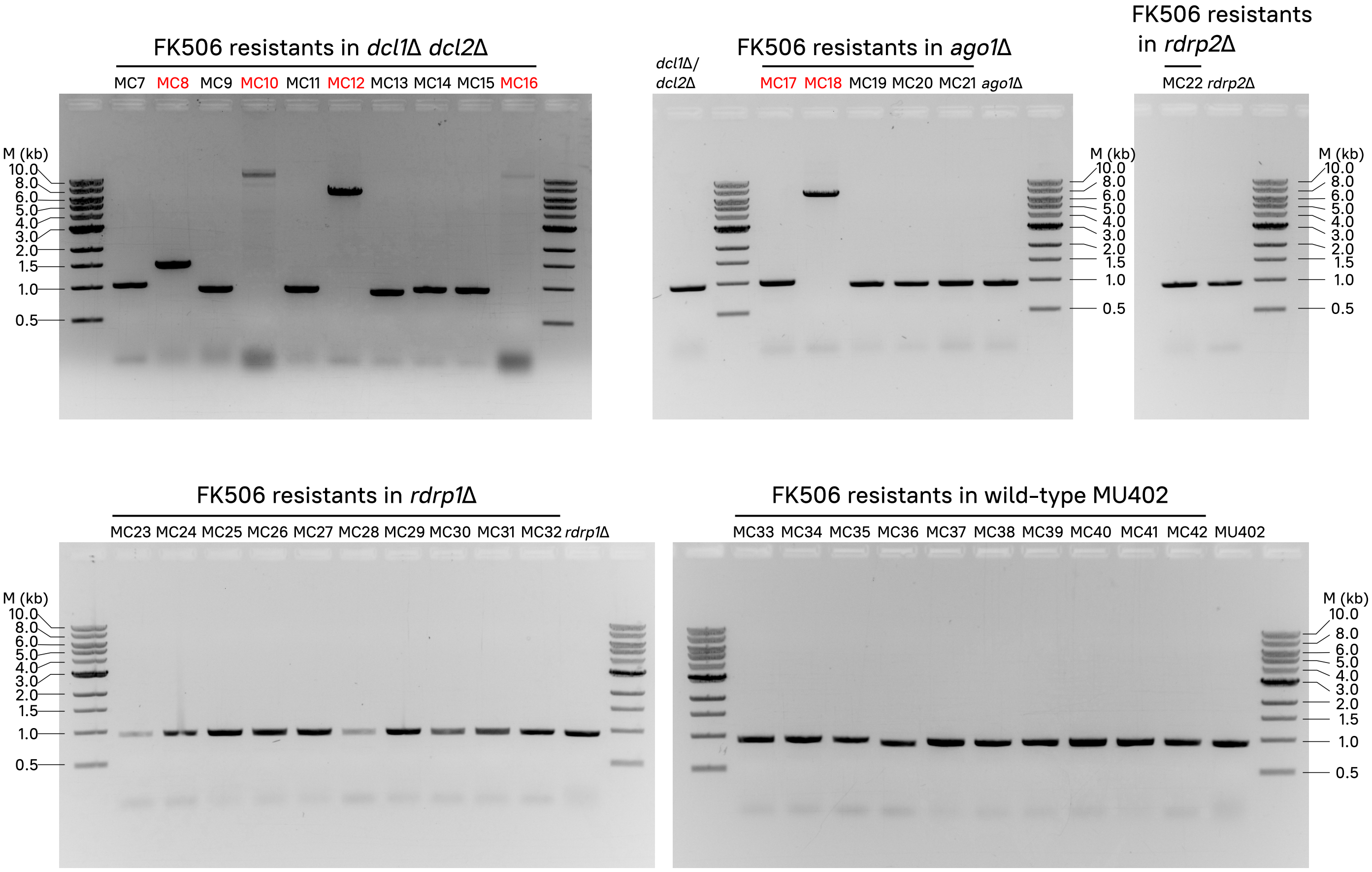


**Fig. S10.** Loss of canonical Dicer or Ago activities leads to large insertions and FK506 target disruption. The FKBP12-encoding *fkbA* locus was PCR-amplified in FK506-resistant isolates in different RNAi-deficient backgrounds (shown on top of each electrophoresis gel picture). *fkbA* sequences containing insertions (also confirmed by Sanger-sequencing) are highlighted in red, and were only found in *dcl1*Δ *dcl2*Δ and *ago1*Δ mutant strains. RNAi-deficient and wild-type strains not exposed to FK506 were also analyzed and included as FK506-sensitive controls. Lanes marked as M indicate DNA ladder and expected fragment sizes.

**Table S1. Transposable elements identified in *M. lusitanicus*[1]**

| Name | Scaffold | Start | End | Strand |
|---|---|---|---|---|
| RChelitron-2_ML_1 | scaffold_1 | 59075 | 64314 | + |
| Tc1Mariner-2_ML_2 | scaffold_1 | 506088 | 507741 | + |
| Tc1Mariner-4_ML_3 | scaffold_1 | 533090 | 534276 | - |
| Gypsy-11_ML_4 | scaffold_1 | 548786 | 550886 | + |
| LINE-19_ML_5 | scaffold_1 | 636962 | 638300 | - |
| DIRS1-1_ML_6 | scaffold_1 | 669754 | 671914 | - |
| Tc1Mariner-2_ML_7 | scaffold_1 | 680928 | 682581 | - |
| Tc1Mariner-1_ML_8 | scaffold_1 | 767427 | 769074 | - |
| Tc1Mariner-2_ML_9 | scaffold_1 | 900299 | 901952 | + |
| PIFharbinger-2_ML_10 | scaffold_1 | 1075426 | 1076217 | + |
| PIFharbinger-10_ML_11 | scaffold_1 | 1083344 | 1084188 | + |
| RChelitron-1_ML_12 | scaffold_1 | 1095022 | 1105954 | + |
| LINE-11_ML_13 | scaffold_1 | 1206459 | 1207551 | - |
| LINE-6_ML_14 | scaffold_1 | 1232677 | 1237995 | + |
| LINE-7_ML_15 | scaffold_1 | 1239265 | 1240306 | - |
| LINE-11_ML_16 | scaffold_1 | 1246524 | 1247309 | - |
| Gypsy-1_ML_17 | scaffold_1 | 1337965 | 1343514 | - |
| LINE-8_ML_18 | scaffold_1 | 1470621 | 1472646 | + |
| Tc1Mariner-1_ML_19 | scaffold_1 | 1476466 | 1478114 | + |
| PIFharbinger-2_ML_20 | scaffold_1 | 1642829 | 1643621 | + |
| Gypsy-13_ML_21 | scaffold_1 | 1712849 | 1715105 | + |
| PIFharbinger-1_ML_22 | scaffold_1 | 1813721 | 1816602 | - |
| RChelitron-2_ML_23 | scaffold_1 | 2145195 | 2150369 | + |
| Tc1Mariner-2_ML_24 | scaffold_1 | 2239161 | 2240812 | + |
| LINE-3_ML_25 | scaffold_1 | 2240968 | 2242011 | - |
| PIFharbinger-3_ML_26 | scaffold_1 | 2243268 | 2244195 | + |
| LINE-3_ML_27 | scaffold_1 | 2249852 | 2250558 | - |
| Tc1Mariner-2_ML_28 | scaffold_1 | 2250841 | 2252493 | - |
| LINE-3_ML_29 | scaffold_1 | 2252491 | 2257999 | - |
| RChelitron-1_ML_30 | scaffold_1 | 2409515 | 2415660 | + |
| Gypsy-1_ML_31 | scaffold_1 | 2443021 | 2448569 | - |
| Gypsy-10_ML_32 | scaffold_1 | 2450069 | 2450900 | - |
| Tc1Mariner-1_ML_33 | scaffold_1 | 2505397 | 2507045 | + |
| Gypsy-14_ML_34 | scaffold_1 | 2602990 | 2607776 | - |
| PIFharbinger-1_ML_35 | scaffold_1 | 2724893 | 2727772 | - |
| LINE-9_ML_36 | scaffold_1 | 2750807 | 2751366 | + |
| LINE-11_ML_37 | scaffold_1 | 2874519 | 2875619 | + |
| Gypsy-9_ML_38 | scaffold_1 | 3232247 | 3237336 | - |
| PIFharbinger-1_ML_39 | scaffold_1 | 3238966 | 3241846 | + |
| Tc1Mariner-5_ML_40 | scaffold_1 | 3327534 | 3329207 | - |
| Gypsy-6_ML_41 | scaffold_1 | 3333312 | 3334483 | - |
| Tc1Mariner-1_ML_42 | scaffold_1 | 3334961 | 3336609 | + |

| Name | Scaffold | Start | End | Strand |
|---|---|---|---|---|
| Tc1Mariner-2_ML_43 | scaffold_1 | 3338046 | 3339699 | + |
| Tc1Mariner-2_ML_44 | scaffold_1 | 3367481 | 3369134 | + |
| Tc1Mariner-1_ML_45 | scaffold_1 | 3450298 | 3451947 | - |
| RChelitron-1_ML_46 | scaffold_1 | 3550469 | 3556685 | - |
| Tc1Mariner-1_ML_47 | scaffold_1 | 3607135 | 3609115 | - |
| Tc1Mariner-11_ML_48 | scaffold_1 | 3847065 | 3847738 | + |
| RChelitron-2_ML_49 | scaffold_1 | 3855592 | 3860835 | + |
| Tc1Mariner-3_ML_50 | scaffold_1 | 3901016 | 3902677 | - |
| Tc1Mariner-3_ML_51 | scaffold_1 | 3944010 | 3945671 | + |
| PIFharbinger-2_ML_52 | scaffold_1 | 3953490 | 3954284 | - |
| Tc1Mariner-3_ML_53 | scaffold_1 | 3983408 | 3985070 | - |
| Gypsy-15_ML_54 | scaffold_1 | 4362330 | 4368330 | + |
| Gypsy-15_ML_55 | scaffold_1 | 4413846 | 4420198 | - |
| Gypsy-2_ML_56 | scaffold_1 | 4471178 | 4477260 | - |
| Tc1Mariner-5_ML_57 | scaffold_1 | 4477992 | 4479884 | + |
| Tc1Mariner-2_ML_58 | scaffold_1 | 4666907 | 4668560 | + |
| Gypsy-2_ML_59 | scaffold_1 | 4889828 | 4895911 | + |
| RChelitron-1_ML_60 | scaffold_1 | 4904164 | 4915041 | - |
| LINE-L1-1_ML_61 | scaffold_1 | 4974903 | 4977632 | + |
| Tc1Mariner-1_ML_62 | scaffold_1 | 4998772 | 5000420 | - |
| LINE-6_ML_63 | scaffold_1 | 5096344 | 5098227 | - |
| DIRS1-2_ML_64 | scaffold_1 | 5154541 | 5155087 | - |
| PIFharbinger-2_ML_65 | scaffold_1 | 5235549 | 5236343 | + |
| Tc1Mariner-3_ML_66 | scaffold_1 | 5317802 | 5319463 | + |
| Tc1Mariner-3_ML_67 | scaffold_1 | 5322967 | 5324628 | - |
| Tc1Mariner-1_ML_68 | scaffold_1 | 5475065 | 5476714 | + |
| Gypsy-16_ML_69 | scaffold_1 | 5504344 | 5505622 | + |
| Tc1Mariner-3_ML_70 | scaffold_1 | 5616002 | 5617663 | - |
| PIFharbinger-4_ML_71 | scaffold_1 | 5617617 | 5618148 | - |
| GremLINE1-4_ML_72 | scaffold_1 | 5621128 | 5622681 | - |
| GremLINE1-1_ML_73 | scaffold_1 | 5623342 | 5629299 | + |
| GremLINE1-1_ML_74 | scaffold_1 | 5629440 | 5635388 | + |
| GremLINE1-1_ML_75 | scaffold_1 | 5635389 | 5640055 | + |
| Gypsy-4_ML_76 | scaffold_10 | 0 | 5223 | - |
| Tc1Mariner-1_ML_77 | scaffold_10 | 260462 | 262111 | + |
| LINE-14_ML_78 | scaffold_10 | 288135 | 290462 | + |
| Gypsy-1_ML_79 | scaffold_10 | 320761 | 323377 | + |
| Gypsy-10_ML_80 | scaffold_10 | 715965 | 716818 | + |
| Gypsy-17_ML_81 | scaffold_10 | 722374 | 724472 | - |
| Gypsy-4_ML_82 | scaffold_10 | 726057 | 732172 | + |
| LINE-19_ML_83 | scaffold_10 | 794118 | 799540 | - |
| Tc1Mariner-2_ML_84 | scaffold_10 | 800066 | 801718 | - |
| LINE-6_ML_85 | scaffold_11 | 142409 | 143871 | + |

| Name | Scaffold | Start | End | Strand |
|---|---|---|---|---|
| Tc1Mariner-4_ML_86 | scaffold_11 | 439043 | 440333 | - |
| Tc1Mariner-3_ML_87 | scaffold_11 | 499857 | 501519 | - |
| LINE-3_ML_88 | scaffold_11 | 501932 | 507440 | + |
| Tc1Mariner-6_ML_89 | scaffold_11 | 552119 | 553213 | - |
| Gypsy-2_ML_90 | scaffold_11 | 872668 | 878748 | + |
| GremLINE1-1_ML_91 | scaffold_11 | 884717 | 885276 | - |
| GremLINE1-1_ML_92 | scaffold_11 | 885902 | 891861 | + |
| GremLINE1-1_ML_93 | scaffold_11 | 891941 | 897425 | + |
| Gypsy-10_ML_94 | scaffold_12 | 70333 | 71183 | + |
| LINE-11_ML_95 | scaffold_12 | 130570 | 131190 | + |
| Gypsy-10_ML_96 | scaffold_12 | 223130 | 223982 | + |
| DIRS1-3_ML_97 | scaffold_12 | 406677 | 408738 | - |
| Tc1Mariner-6_ML_98 | scaffold_12 | 494042 | 495098 | + |
| Gypsy-18_ML_99 | scaffold_12 | 561593 | 563744 | - |
| Gypsy-19_ML_100 | scaffold_12 | 565907 | 567095 | + |
| LINE-9_ML_101 | scaffold_12 | 759313 | 764708 | + |
| GremLINE1-1_ML_102 | scaffold_12 | 780213 | 786148 | - |
| GremLINE1-1_ML_103 | scaffold_12 | 786153 | 792098 | - |
| GremLINE1-4_ML_104 | scaffold_12 | 792097 | 798024 | - |
| GremLINE1-4_ML_105 | scaffold_12 | 798150 | 804078 | - |
| GremLINE1-1_ML_106 | scaffold_12 | 804204 | 809647 | - |
| PIFharbinger-5_ML_107 | scaffold_13 | 69983 | 71315 | - |
| Gypsy-1_ML_108 | scaffold_13 | 115801 | 121345 | - |
| RChelitron-2_ML_109 | scaffold_13 | 426726 | 431888 | + |
| Tc1Mariner-1_ML_110 | scaffold_13 | 506811 | 508460 | + |
| RChelitron-4_ML_111 | scaffold_13 | 576863 | 577709 | + |
| PIFharbinger-6_ML_112 | scaffold_13 | 577760 | 579284 | + |
| LINE-L1-1_ML_113 | scaffold_13 | 642983 | 644422 | + |
| Tc1Mariner-2_ML_114 | scaffold_15 | 25664 | 27317 | - |
| GremLINE1-4_ML_115 | scaffold_15 | 104067 | 109983 | - |
| GremLINE1-1_ML_116 | scaffold_15 | 109988 | 115956 | - |
| GremLINE1-4_ML_117 | scaffold_15 | 115977 | 121903 | - |
| GremLINE1-1_ML_118 | scaffold_15 | 122023 | 127963 | - |
| GremLINE1-1_ML_119 | scaffold_15 | 127962 | 133913 | - |
| GremLINE1-2_ML_120 | scaffold_15 | 133946 | 137282 | - |
| Piggybac-1_Mcir_121 | scaffold_16 | 33804 | 35019 | + |
| LINE-16_ML_122 | scaffold_17 | 0 | 4474 | - |
| GremLINE1-1_ML_123 | scaffold_17 | 4582 | 10517 | - |
| GremLINE1-4_ML_124 | scaffold_17 | 11176 | 12729 | + |
| PIFharbinger-4_ML_125 | scaffold_17 | 15709 | 16240 | + |
| Tc1Mariner-3_ML_126 | scaffold_17 | 16194 | 17855 | + |
| GremLINE1-3_ML_127 | scaffold_18 | 29067 | 31827 | - |
| Gypsy-4_ML_128 | scaffold_2 | 344999 | 351242 | - |

| Name | Scaffold | Start | End | Strand |
|---|---|---|---|---|
| Gypsy-5_ML_129 | scaffold_2 | 406710 | 412700 | - |
| Gypsy-5_ML_130 | scaffold_2 | 517655 | 523645 | + |
| Tc1Mariner-3_ML_131 | scaffold_2 | 639810 | 641480 | + |
| LINE-11_ML_132 | scaffold_2 | 725212 | 725916 | + |
| Tc1Mariner-2_ML_133 | scaffold_2 | 935659 | 937311 | - |
| LINE-L1-1_ML_134 | scaffold_2 | 1032990 | 1035596 | - |
| LINE-10_ML_135 | scaffold_2 | 1033400 | 1038935 | - |
| RChelitron-1_ML_136 | scaffold_2 | 1202553 | 1208001 | + |
| RChelitron-2_ML_137 | scaffold_2 | 1391435 | 1396610 | - |
| Gypsy-4_ML_138 | scaffold_2 | 1492129 | 1498180 | + |
| Gypsy-3_ML_139 | scaffold_2 | 1498772 | 1504699 | + |
| Gypsy-6_ML_140 | scaffold_2 | 1526998 | 1531557 | - |
| LINE-11_ML_141 | scaffold_2 | 1557397 | 1560142 | + |
| Tc1Mariner-1_ML_142 | scaffold_2 | 1561960 | 1563940 | - |
| Gypsy-9_ML_143 | scaffold_2 | 1674234 | 1675012 | - |
| LINE-10_ML_144 | scaffold_2 | 1701120 | 1702220 | + |
| LINE-L1-1_ML_145 | scaffold_2 | 1701125 | 1702832 | + |
| Gypsy-20_ML_146 | scaffold_2 | 1722870 | 1723542 | - |
| Tc1Mariner-2_ML_147 | scaffold_2 | 1790854 | 1792507 | + |
| RChelitron-1_ML_148 | scaffold_2 | 1859355 | 1870298 | - |
| RChelitron-1_ML_149 | scaffold_2 | 2110168 | 2116283 | + |
| Tc1Mariner-3_ML_150 | scaffold_2 | 2325951 | 2327611 | - |
| Tc1Mariner-4_ML_151 | scaffold_2 | 2430681 | 2432095 | - |
| RChelitron-1_ML_152 | scaffold_2 | 2489109 | 2490823 | + |
| Tc1Mariner-1_ML_153 | scaffold_2 | 2697135 | 2698782 | - |
| Gypsy-10_ML_154 | scaffold_2 | 2845958 | 2852009 | + |
| Gypsy-6_ML_155 | scaffold_2 | 3029085 | 3033644 | - |
| LTR/Unknown-1_ML_156 | scaffold_2 | 3168609 | 3170817 | + |
| Gypsy-10_ML_157 | scaffold_2 | 3443399 | 3449982 | - |
| LINE-11_ML_158 | scaffold_2 | 3575861 | 3577891 | + |
| Gypsy-4_ML_159 | scaffold_2 | 3758052 | 3764103 | - |
| LINE-L1-1_ML_160 | scaffold_2 | 3792266 | 3794144 | + |
| LINE-10_ML_161 | scaffold_2 | 4258762 | 4264208 | + |
| LINE-L1-3_ML_162 | scaffold_2 | 4264208 | 4264708 | + |
| RChelitron-1_ML_163 | scaffold_2 | 4289389 | 4295519 | - |
| Piggybac-1_Mcir_164 | scaffold_2 | 4314014 | 4315852 | - |
| LINE-L1-1_ML_165 | scaffold_2 | 4352316 | 4354834 | - |
| Gypsy-3_ML_166 | scaffold_2 | 4362188 | 4368115 | + |
| Tc1Mariner-1_ML_167 | scaffold_2 | 4381538 | 4383184 | - |
| Tc1Mariner-3_ML_168 | scaffold_2 | 4679027 | 4680689 | - |
| GremLINE1-2_ML_169 | scaffold_2 | 4707808 | 4713773 | - |
| GremLINE1-2_ML_170 | scaffold_2 | 4714562 | 4716735 | + |
| Tc1Mariner-3_ML_171 | scaffold_3 | 417685 | 419345 | + |

| Name | Scaffold | Start | End | Strand |
|---|---|---|---|---|
| Gypsy-2_ML_172 | scaffold_3 | 420421 | 426501 | + |
| Tc1Mariner-2_ML_173 | scaffold_3 | 502955 | 504608 | - |
| LINE-19_ML_174 | scaffold_3 | 679573 | 681107 | + |
| LINE-L1-3_ML_175 | scaffold_3 | 762959 | 768378 | - |
| Tc1Mariner-1_ML_176 | scaffold_3 | 768470 | 770117 | - |
| DIRS1-4_ML_177 | scaffold_3 | 773443 | 775369 | + |
| PIFharbinger-3_ML_178 | scaffold_3 | 776054 | 776981 | + |
| LINE-6_ML_179 | scaffold_3 | 939860 | 945178 | - |
| Tc1Mariner-3_ML_180 | scaffold_3 | 954500 | 956161 | - |
| Tc1Mariner-3_ML_181 | scaffold_3 | 959093 | 960754 | + |
| Gypsy-5_ML_182 | scaffold_3 | 989556 | 995546 | - |
| Tc1Mariner-1_ML_183 | scaffold_3 | 1019051 | 1021173 | + |
| Copia_1_ML_184 | scaffold_3 | 1106691 | 1108038 | - |
| Tc1Mariner-1_ML_185 | scaffold_3 | 1107973 | 1109621 | + |
| LINE-13_ML_186 | scaffold_3 | 1164914 | 1166593 | + |
| LINE-9_ML_187 | scaffold_3 | 1590619 | 1593886 | + |
| LINE-10_ML_188 | scaffold_3 | 1985187 | 1990634 | + |
| LINE-L1-3_ML_189 | scaffold_3 | 1990634 | 1991221 | + |
| LINE-14_ML_190 | scaffold_3 | 2301870 | 2305301 | - |
| Tc1Mariner-2_ML_191 | scaffold_3 | 2305763 | 2307418 | - |
| Tc1Mariner-4_ML_192 | scaffold_3 | 2381467 | 2382888 | - |
| Gypsy-22_ML_193 | scaffold_3 | 2391890 | 2397659 | + |
| LINE-15_ML_194 | scaffold_3 | 2470995 | 2476429 | + |
| Tc1Mariner-1_ML_195 | scaffold_3 | 2573625 | 2575271 | + |
| LINE-11_ML_196 | scaffold_3 | 2689286 | 2691082 | - |
| Tc1Mariner-1_ML_197 | scaffold_3 | 2836997 | 2838646 | - |
| LINE-L1-1_ML_198 | scaffold_3 | 2941233 | 2942449 | - |
| Tc1Mariner-2_ML_199 | scaffold_3 | 3155493 | 3157146 | - |
| Gypsy-3_ML_200 | scaffold_3 | 3277815 | 3283742 | - |
| LINE-15_ML_201 | scaffold_3 | 3302882 | 3307116 | - |
| PIFharbinger-5_ML_202 | scaffold_3 | 3641565 | 3642897 | - |
| Tc1Mariner-3_ML_203 | scaffold_3 | 3704235 | 3705896 | + |
| GremLINE1-4_ML_204 | scaffold_3 | 3776246 | 3782169 | + |
| GremLINE1-1_ML_205 | scaffold_3 | 3785060 | 3791025 | + |
| GremLINE1-1_ML_206 | scaffold_3 | 3791202 | 3797165 | + |
| GremLINE1-1_ML_207 | scaffold_3 | 3797164 | 3802935 | + |
| GremLINE1-1_ML_208 | scaffold_3 | 3802934 | 3808709 | + |
| GremLINE1-1_ML_209 | scaffold_3 | 3808709 | 3814476 | + |
| GremLINE1-4_ML_210 | scaffold_3 | 3814534 | 3820460 | + |
| GremLINE1-1_ML_211 | scaffold_3 | 3820464 | 3826333 | + |
| GremLINE1-1_ML_212 | scaffold_3 | 3826333 | 3832204 | + |
| GremLINE1-1_ML_213 | scaffold_3 | 3841429 | 3843599 | - |
| Gypsy-23_ML_214 | scaffold_3 | 4074669 | 4075740 | - |

| Name | Scaffold | Start | End | Strand |
|---|---|---|---|---|
| Tc1Mariner-1_ML_215 | scaffold_3 | 4121604 | 4123253 | - |
| Tc1Mariner-3_ML_216 | scaffold_3 | 4123962 | 4125621 | - |
| LINE-10_ML_217 | scaffold_3 | 4135453 | 4140881 | + |
| LINE-L1-1_ML_218 | scaffold_3 | 4138530 | 4141466 | + |
| LINE-11_ML_219 | scaffold_3 | 4264330 | 4265982 | + |
| Tc1Mariner-4_ML_220 | scaffold_3 | 4292552 | 4293974 | - |
| PIFharbinger-7_ML_221 | scaffold_3 | 4323022 | 4325285 | + |
| PIFharbinger-5_ML_222 | scaffold_3 | 4361511 | 4362843 | + |
| LINE-19_ML_223 | scaffold_3 | 4435732 | 4436380 | - |
| Tc1Mariner-1_ML_224 | scaffold_3 | 4520503 | 4522149 | - |
| Tc1Mariner-9_ML_225 | scaffold_4 | 54645 | 56694 | - |
| Tc1Mariner-1_ML_226 | scaffold_4 | 138792 | 140441 | + |
| Tc1Mariner-4_ML_227 | scaffold_4 | 930365 | 931779 | + |
| Tc1Mariner-1_ML_228 | scaffold_4 | 1078431 | 1080079 | - |
| PIFharbinger-1_ML_229 | scaffold_4 | 1150233 | 1153114 | - |
| LTR/Unknown-2_ML_230 | scaffold_4 | 1547088 | 1548120 | - |
| Gypsy-4_ML_231 | scaffold_4 | 1592443 | 1598494 | - |
| Gypsy-10_ML_232 | scaffold_4 | 1604518 | 1605370 | - |
| DIRS1-5_ML_233 | scaffold_4 | 1731081 | 1733187 | - |
| LINE-9_ML_234 | scaffold_4 | 1735810 | 1741203 | - |
| Tc1Mariner-1_ML_235 | scaffold_4 | 1801532 | 1803178 | - |
| RChelitron-2_ML_236 | scaffold_4 | 1820957 | 1826183 | + |
| Tc1Mariner-1_ML_237 | scaffold_4 | 1840378 | 1842024 | - |
| Gypsy-24_ML_238 | scaffold_4 | 1940329 | 1940881 | + |
| LINE-3_ML_239 | scaffold_4 | 2270490 | 2271238 | - |
| LINE-3_ML_240 | scaffold_4 | 2272906 | 2278414 | - |
| PIFharbinger-8_ML_241 | scaffold_4 | 2322835 | 2323405 | - |
| LINE-3_ML_242 | scaffold_4 | 2336014 | 2341523 | + |
| LINE-3_ML_243 | scaffold_4 | 2341803 | 2342610 | + |
| LINE-3_ML_244 | scaffold_4 | 2342607 | 2343521 | + |
| GremLINE1-4_ML_245 | scaffold_4 | 2351462 | 2357388 | - |
| GremLINE1-3_ML_246 | scaffold_4 | 2357389 | 2363323 | - |
| GremLINE1-2_ML_247 | scaffold_4 | 2363869 | 2369818 | + |
| GremLINE1-4_ML_248 | scaffold_4 | 2369818 | 2375739 | + |
| GremLINE1-1_ML_249 | scaffold_4 | 2375800 | 2381730 | + |
| GremLINE1-5_ML_250 | scaffold_4 | 2421133 | 2421982 | - |
| Gypsy-5_ML_251 | scaffold_4 | 2491357 | 2497347 | - |
| PIFharbinger-9_ML_252 | scaffold_4 | 2526978 | 2527863 | + |
| PIFharbinger-10_ML_253 | scaffold_4 | 2635220 | 2636744 | - |
| RChelitron-1_ML_254 | scaffold_4 | 2863402 | 2874253 | - |
| Gypsy-25_ML_255 | scaffold_4 | 2894620 | 2896228 | - |
| PIFharbinger-1_ML_256 | scaffold_4 | 2953056 | 2954339 | + |
| Gypsy-5_ML_257 | scaffold_4 | 3002155 | 3008145 | - |

| Name | Scaffold | Start | End | Strand |
|---|---|---|---|---|
| Tc1Mariner-4_ML_258 | scaffold_4 | 3291272 | 3292781 | + |
| RChelitron-1_ML_259 | scaffold_4 | 3547034 | 3553272 | + |
| Gypsy-1_ML_260 | scaffold_4 | 3900790 | 3902041 | - |
| LINE-11_ML_261 | scaffold_4 | 4154022 | 4155120 | - |
| LINE-8_ML_262 | scaffold_4 | 4172441 | 4174172 | + |
| Tc1Mariner-2_ML_263 | scaffold_4 | 4235272 | 4236924 | + |
| Gypsy-6_ML_264 | scaffold_4 | 4237192 | 4241752 | - |
| LINE-19_ML_265 | scaffold_4 | 4451684 | 4457131 | - |
| Gypsy-4_ML_266 | scaffold_4 | 4555284 | 4561335 | - |
| Gypsy-6_ML_267 | scaffold_5 | 84402 | 88961 | + |
| RChelitron-2_ML_268 | scaffold_5 | 250271 | 255502 | - |
| Tc1Mariner-2_ML_269 | scaffold_5 | 329309 | 330962 | - |
| Gypsy-2_ML_270 | scaffold_5 | 331799 | 337905 | + |
| RChelitron-1_ML_271 | scaffold_5 | 486386 | 491849 | + |
| Tc1Mariner-4_ML_272 | scaffold_5 | 512530 | 514039 | - |
| Gypsy-5_ML_273 | scaffold_5 | 563349 | 569339 | - |
| Gypsy-7_ML_274 | scaffold_5 | 1022147 | 1027234 | + |
| Tc1Mariner-3_ML_275 | scaffold_5 | 1372092 | 1373753 | - |
| Tc1Mariner-3_ML_276 | scaffold_5 | 1401187 | 1402848 | - |
| LINE-11_ML_277 | scaffold_5 | 1572568 | 1574059 | + |
| Tc1Mariner-2_ML_278 | scaffold_5 | 1742650 | 1744303 | - |
| Tc1Mariner-1_ML_279 | scaffold_5 | 1749689 | 1751334 | - |
| Tc1Mariner-2_ML_280 | scaffold_5 | 1858153 | 1859806 | + |
| Gypsy-26_ML_281 | scaffold_5 | 1860013 | 1862481 | - |
| LINE-11_ML_282 | scaffold_5 | 1866061 | 1868081 | - |
| Gypsy-10_ML_283 | scaffold_5 | 1913934 | 1914784 | + |
| PIFharbinger-1_ML_284 | scaffold_5 | 2107720 | 2110600 | + |
| LINE-L1-1_ML_285 | scaffold_5 | 2230378 | 2232693 | + |
| LINE-11_ML_286 | scaffold_5 | 2455584 | 2457909 | + |
| Gypsy-8_ML_287 | scaffold_5 | 2595029 | 2596462 | - |
| LINE-20_ML_288 | scaffold_5 | 2596676 | 2598827 | - |
| Tc1Mariner-3_ML_289 | scaffold_5 | 2638987 | 2640648 | - |
| RChelitron-1_ML_290 | scaffold_5 | 2652066 | 2658188 | - |
| Gypsy-6_ML_291 | scaffold_5 | 2709451 | 2714010 | - |
| Tc1Mariner-1_ML_292 | scaffold_5 | 2719174 | 2720820 | + |
| LINE-10_ML_293 | scaffold_5 | 2814226 | 2815648 | + |
| LINE-21_ML_294 | scaffold_5 | 2960906 | 2964299 | + |
| PIFharbinger-11_ML_295 | scaffold_5 | 3088851 | 3089915 | - |
| Tc1Mariner-2_ML_296 | scaffold_5 | 3091517 | 3093169 | + |
| Tc1Mariner-2_ML_297 | scaffold_5 | 3208138 | 3209791 | - |
| LINE-21_ML_298 | scaffold_5 | 3212209 | 3215597 | - |
| Tc1Mariner-3_ML_299 | scaffold_5 | 3359363 | 3361024 | - |
| GremLINE1-1_ML_300 | scaffold_5 | 3401993 | 3403514 | - |

| Name | Scaffold | Start | End | Strand |
|---|---|---|---|---|
| GremLINE1-1_ML_301 | scaffold_5 | 3404199 | 3408900 | + |
| GremLINE1-1_ML_302 | scaffold_5 | 3408903 | 3414868 | + |
| GremLINE1-1_ML_303 | scaffold_5 | 3414883 | 3420759 | + |
| GremLINE1-1_ML_304 | scaffold_5 | 3420758 | 3426676 | + |
| PIFharbinger-11_ML_305 | scaffold_6 | 189507 | 190584 | - |
| Gypsy-27_ML_306 | scaffold_6 | 261456 | 262131 | - |
| Tc1Mariner-2_ML_307 | scaffold_6 | 484064 | 485718 | + |
| LINE-L1-1_ML_308 | scaffold_6 | 798979 | 801413 | + |
| Gypsy-10_ML_309 | scaffold_6 | 818480 | 819333 | + |
| Gypsy-10_ML_310 | scaffold_6 | 821986 | 822839 | - |
| Piggybac-1_Mcir_311 | scaffold_6 | 825490 | 826708 | + |
| Gypsy-5_ML_312 | scaffold_6 | 1201703 | 1207692 | + |
| LINE-L1-1_ML_313 | scaffold_6 | 1261081 | 1262188 | - |
| Tc1Mariner-2_ML_314 | scaffold_6 | 1295554 | 1297207 | + |
| Tc1Mariner-1_ML_315 | scaffold_6 | 1301159 | 1302808 | - |
| Tc1Mariner-1_ML_316 | scaffold_6 | 1398415 | 1400061 | + |
| RChelitron-1_ML_317 | scaffold_6 | 1476610 | 1478314 | + |
| Gypsy-3_ML_318 | scaffold_6 | 1480244 | 1486171 | + |
| Tc1Mariner-2_ML_319 | scaffold_6 | 1497219 | 1498723 | + |
| Gypsy-6_ML_320 | scaffold_6 | 1500247 | 1504806 | + |
| Tc1Mariner-1_ML_321 | scaffold_6 | 1623151 | 1624799 | - |
| Tc1Mariner-1_ML_322 | scaffold_6 | 1627994 | 1629640 | + |
| Gypsy-10_ML_323 | scaffold_6 | 1705645 | 1706260 | + |
| Gypsy-9_ML_324 | scaffold_6 | 2087868 | 2090076 | + |
| Tc1Mariner-2_ML_325 | scaffold_6 | 2323793 | 2325446 | + |
| Gypsy-10_ML_326 | scaffold_6 | 2834217 | 2835073 | + |
| LINE-L1-4.1_ML_327 | scaffold_6 | 2862505 | 2863720 | + |
| RChelitron-1_ML_328 | scaffold_6 | 2952733 | 2958881 | - |
| Tc1Mariner-1_ML_329 | scaffold_6 | 3012273 | 3013922 | + |
| Tc1Mariner-2_ML_330 | scaffold_6 | 3025523 | 3027175 | - |
| Tc1Mariner-1_ML_331 | scaffold_6 | 3076254 | 3077903 | + |
| Gypsy-1_ML_332 | scaffold_6 | 3112118 | 3117663 | + |
| GremLINE1-1_ML_333 | scaffold_6 | 3187674 | 3193633 | + |
| GremLINE1-1_ML_334 | scaffold_6 | 3193649 | 3199609 | + |
| Tc1Mariner-5_ML_335 | scaffold_7 | 0 | 1379 | - |
| LINE-L1-3_ML_336 | scaffold_7 | 264447 | 265025 | - |
| LINE-10_ML_337 | scaffold_7 | 265025 | 270471 | - |
| LINE-11_ML_338 | scaffold_7 | 389869 | 391725 | + |
| Gypsy-7_ML_339 | scaffold_7 | 447265 | 449599 | - |
| Gypsy-3_ML_340 | scaffold_7 | 450169 | 456096 | - |
| Gypsy-28_ML_341 | scaffold_7 | 459419 | 462658 | + |
| LTR/Unknown-3_ML_342 | scaffold_7 | 462998 | 463919 | + |
| LINE-L1-1_ML_343 | scaffold_7 | 625148 | 627509 | + |

| Name | Scaffold | Start | End | Strand |
|---|---|---|---|---|
| RChelitron-1_ML_344 | scaffold_7 | 1040083 | 1051000 | - |
| Gypsy-9_ML_345 | scaffold_7 | 1086990 | 1092080 | - |
| Tc1Mariner-1_ML_346 | scaffold_7 | 1365099 | 1366748 | + |
| LINE-22_ML_347 | scaffold_7 | 1367512 | 1370430 | - |
| Gypsy-2_ML_348 | scaffold_7 | 1376029 | 1382112 | - |
| Gypsy-29_ML_349 | scaffold_7 | 1477134 | 1478139 | - |
| Gypsy-8_ML_350 | scaffold_7 | 1834035 | 1840246 | - |
| Gypsy-5_ML_351 | scaffold_7 | 2176890 | 2182879 | + |
| Tc1Mariner-4_ML_352 | scaffold_7 | 2219779 | 2221201 | - |
| Tc1Mariner-1_ML_353 | scaffold_7 | 2286049 | 2287698 | - |
| LINE-L1-1_ML_354 | scaffold_7 | 2320671 | 2321794 | - |
| Tc1Mariner-2_ML_355 | scaffold_7 | 2445681 | 2447333 | + |
| LINE-L1-1_ML_356 | scaffold_7 | 2545528 | 2546696 | - |
| RChelitron-5_ML_357 | scaffold_7 | 2574282 | 2579010 | + |
| Tc1Mariner-4_ML_358 | scaffold_7 | 2628287 | 2629080 | + |
| Tc1Mariner-2_ML_359 | scaffold_7 | 2755486 | 2757140 | + |
| Gypsy-3_ML_360 | scaffold_7 | 2771143 | 2777070 | - |
| Tc1Mariner-4_ML_361 | scaffold_7 | 2808421 | 2809849 | - |
| LINE-11_ML_362 | scaffold_7 | 2851478 | 2852831 | + |
| LINE-3_ML_363 | scaffold_8 | 46922 | 49246 | - |
| GremLINE1-1_ML_364 | scaffold_8 | 134056 | 140018 | - |
| GremLINE1-1_ML_365 | scaffold_8 | 140018 | 145960 | - |
| GremLINE1-1_ML_366 | scaffold_8 | 145984 | 151949 | - |
| GremLINE1-2_ML_367 | scaffold_8 | 152588 | 158554 | + |
| GremLINE1-1_ML_368 | scaffold_8 | 158585 | 164538 | + |
| GremLINE1-1_ML_369 | scaffold_8 | 164538 | 170493 | + |
| GremLINE1-1_ML_370 | scaffold_8 | 170494 | 176430 | + |
| GremLINE1-1_ML_371 | scaffold_8 | 176616 | 182576 | + |
| LINE-19_ML_372 | scaffold_8 | 472858 | 478330 | + |
| Gypsy-5_ML_373 | scaffold_8 | 566194 | 572184 | - |
| Tc1Mariner-1_ML_374 | scaffold_8 | 972839 | 974488 | + |
| Tc1Mariner-10_ML_375 | scaffold_8 | 1266061 | 1266745 | + |
| Piggybac-1_Mcir_376 | scaffold_8 | 1345074 | 1346615 | + |
| PIFharbinger-12_ML_377 | scaffold_8 | 1400011 | 1401187 | + |
| Gypsy-2_ML_378 | scaffold_8 | 1633311 | 1639394 | - |
| Gypsy-5_ML_379 | scaffold_8 | 1768307 | 1774297 | - |
| Tc1Mariner-4_ML_380 | scaffold_8 | 1811043 | 1812457 | - |
| LTR/Unknown-4_ML_381 | scaffold_8 | 1830370 | 1831108 | + |
| Tc1Mariner-1_ML_382 | scaffold_8 | 1936921 | 1938569 | - |
| LINE-L1-1_ML_383 | scaffold_8 | 1974982 | 1977925 | - |
| Gypsy-4_ML_384 | scaffold_9 | 61737 | 68233 | - |
| Gypsy-4_ML_385 | scaffold_9 | 69676 | 75730 | + |
| Gypsy-1_ML_386 | scaffold_9 | 118362 | 123905 | + |

| Name | Scaffold | Start | End | Strand |
|---|---|---|---|---|
| Gypsy-10_ML_387 | scaffold_9 | 167279 | 168127 | - |
| Tc1Mariner-2_ML_388 | scaffold_9 | 301202 | 302854 | + |
| Gypsy-10_ML_389 | scaffold_9 | 351115 | 351975 | + |
| PIFharbinger-5_ML_390 | scaffold_9 | 547442 | 548774 | - |
| Tc1Mariner-2_ML_391 | scaffold_9 | 553986 | 555639 | - |
| LTR/Unknown-5_ML_392 | scaffold_9 | 710552 | 711569 | - |
| Tc1Mariner-2_ML_393 | scaffold_9 | 711521 | 713174 | - |
| RChelitron-2_ML_394 | scaffold_9 | 914572 | 919818 | + |
| PIFharbinger-3_ML_395 | scaffold_9 | 1139875 | 1140770 | - |
| Piggybac-1_Mcir_396 | scaffold_9 | 1269073 | 1271098 | + |
| Gypsy-10_ML_397 | scaffold_9 | 1292831 | 1293685 | + |
| Tc1Mariner-2_ML_398 | scaffold_9 | 1295750 | 1297402 | + |
| Tc1Mariner-1_ML_399 | scaffold_9 | 1299841 | 1301510 | + |
| Tc1Mariner-11_ML_400 | scaffold_9 | 1331458 | 1332640 | - |
| GremLINE1-4_ML_401 | scaffold_9 | 1336100 | 1342022 | - |
| GremLINE1-1_ML_402 | scaffold_9 | 1342055 | 1348020 | - |
| GremLINE1-1_ML_403 | scaffold_9 | 1348702 | 1350387 | + |
| GremLINE1-1_ML_404 | scaffold_9 | 1350401 | 1356320 | + |
| Tc1Mariner-1_ML_405 | scaffold_9 | 1357999 | 1359731 | + |

[1]Class I, RNA transposons or retrotransposons and Class II or DNA transposon predicted in the *M. lusitanicus* MU402 genome. RNA transposons are classified into three major families: GremLINE1, other non-LTR LINE elements, and LTR Ty3/Gypsy; whereas DNA transposons are classified into: Tc1/Mariner, RC/Helitron, and PIF/Harbinger. Every element copy shows a systematic name comprised of the following fields separated by an underscore: transposon subfamily (family and a number delimited by a dash), the acronym of the species in which it was identified (in most cases ML from *M. lusitanicus*), and a number. In addition, their genomic location (scaffold, start, end, and strand) is displayed.

**Table S2. FK506 spontaneous resistant isolates from RNAi-deficient and wild-type strains[1]**

| Strain | Genetic background | Mutation | Effect | GremLINE orientation | Target site duplication |
|---|---|---|---|---|---|
| MC7 | *dcl1Δ dcl2Δ* | g.48-138dup 91 bp duplication | Frameshift, premature STOP codon caused by duplication | | |
| MC8 | *dcl1Δ dcl2Δ* | g.(-8)ins, 534 bp | Frameshift, premature STOP codon caused by insertion | Forward | ATATTAAACATGGGTG |
| MC9 | *dcl1Δ dcl2Δ* | g.76G>T in splicing site | Splice site mutation caused by transversion | | |
| MC10 | *dcl1Δ dcl2Δ* | ~12 kb GremLINE insertion | Unknown | Unknown | Unknown |
| MC11 | *dcl1Δ dcl2Δ* | g.457A>C in stop codon (p.X109Y) | Nonstop mutation caused by transversion | | |
| MC12 | *dcl1Δ dcl2Δ* | g.337ins, 5951 | Frameshift, premature STOP caused by insertion | Forward | ACTCAATTGTCTGTT |
| MC13 | *dcl1Δ dcl2Δ* | g.291-341del 51nt deletion | Splice site mutation caused by transversion | | |
| MC14 | *dcl1Δ dcl2Δ* | g.455delT in stop codon (p.X109K) | Nonstop mutation caused by deletion | | |
| MC15 | *dcl1Δ dcl2Δ* | g.207C>T (p.Q48X) | Premature STOP caused by transition | | |
| MC16 | *dcl1Δ dcl2Δ* | ~12 kb GremLINE insertion | Unknown | Unknown | Unknown |
| MC17 | *ago1Δ* | g.244ins 36nt (p.E71X) | Several missense mutations, premature STOP caused by insertion | Reverse | GGTCAAGTCATCAAGGGTT |
| MC18 | *ago1Δ* | g.(-54)ins, 5955 bp | Promoter region | Forward | AACCAAACGTTTTT |
| MC19 | *ago1Δ* | g.324delCTCA (p.T65N, p.R74X) | Frameshift, premature STOP caused by deletion | | |
| MC20 | *ago1Δ* | g.374G>C (p.A82P) | Missense mutation caused by transversion | | |
| MC21 | *ago1Δ* | g.157delTC (p.L31R, p. G34X) | Frameshift, premature STOP caused by deletion | | |
| MC22 | *rdrp2Δ* | g.118delA (p.K18R, p.V23X) | Frameshift, premature STOP caused by deletion | | |
| MC23 | *rdrp1Δ* | None, wildtype | Possible epimutant | | |
| MC24 | *rdrp1Δ* | None, wildtype | Possible epimutant | | |
| MC25 | *rdrp1Δ* | None, wildtype | Possible epimutant | | |
| MC26 | *rdrp1Δ* | None, wildtype | Possible epimutant | | |
| MC27 | *rdrp1Δ* | None, wildtype | Possible epimutant | | |
| MC28 | *rdrp1Δ* | None, wildtype | Possible epimutant | | |
| MC29 | *rdrp1Δ* | None, wildtype | Possible epimutant | | |
| MC30 | *rdrp1Δ* | None, wildtype | Possible epimutant | | |
| MC31 | *rdrp1Δ* | None, wildtype | Possible epimutant | | |
| MC32 | *rdrp1Δ* | None, wildtype | Possible epimutant | | |
| MC33 | MU402 | g.241G>A (p.G59D) | Missense mutation caused by transition | | |
| MC34 | MU402 | g.341G>T (p.E71X) | Nonsense mutation, premature STOP caused by transversion | | |
| MC35 | MU402 | g.207C>T (p.Q48X) | Nonsense mutation, premature STOP caused by transition | | |
| MC36 | MU402 | g.124del 42bp deletion (p.G20_G34del) | Microdeletion | | |
| MC37 | MU402 | g.357insTGAT (p.Y81X) | Frameshift mutation, premature STOP caused by insertion | | |
| MC38 | MU402 | g.54delTGACC in intron | Possible epimutant | | |
| MC40 | MU402 | g.417delC (p.L98X) | Frameshift mutation, premature STOP caused by deletion | | |

| Strain | Genetic background | Mutation | Effect | GremLINE orientation | Target site duplication |
|---|---|---|---|---|---|
| MC41 | MU402 | None, wildtype | Possible epimutant | | |
| MC42 | MU402 | g.441T>C (p.L104P) | Missense mutation caused by transition | | |

[1]The table shows systematic strain names, genetic background or genotype before the artificial selection experiment and FK506 exposure, mutations characterized, effect of the mutation, GremLINE1 orientation with respect to *fkbA* in those strains that have insertions, and the target duplication site sequences found.

**Table S3. GremLINE1 insertions in canonical RNAi-deficient mutants[1]**

| Query | Subject | % Id | Ins length | Aln length | Mm | Gap | E-value | Score |
|---|---|---|---|---|---|---|---|---|
| ***ago1*Δ_GremLINE1_1 (MC17)** | GremLINE1-1_ML_102 | 100 | 36 | 35 | 0 | 0 | 2.77E-14 | 69.9 |
| | GremLINE1-1_ML_103 | 100 | 36 | 35 | 0 | 0 | 2.77E-14 | 69.9 |
| | GremLINE1-1_ML_106 | 100 | 36 | 35 | 0 | 0 | 2.77E-14 | 69.9 |
| | GremLINE1-1_ML_116 | 100 | 36 | 35 | 0 | 0 | 2.77E-14 | 69.9 |
| | GremLINE1-1_ML_118 | 100 | 36 | 35 | 0 | 0 | 2.77E-14 | 69.9 |
| | GremLINE1-1_ML_119 | 100 | 36 | 35 | 0 | 0 | 2.77E-14 | 69.9 |
| | GremLINE1-1_ML_123 | 100 | 36 | 35 | 0 | 0 | 2.77E-14 | 69.9 |
| | GremLINE1-1_ML_205 | 100 | 36 | 35 | 0 | 0 | 2.77E-14 | 69.9 |
| | GremLINE1-1_ML_206 | 100 | 36 | 35 | 0 | 0 | 2.77E-14 | 69.9 |
| | GremLINE1-1_ML_207 | 100 | 36 | 35 | 0 | 0 | 2.77E-14 | 69.9 |
| | GremLINE1-1_ML_208 | 100 | 36 | 35 | 0 | 0 | 2.77E-14 | 69.9 |
| | GremLINE1-1_ML_209 | 100 | 36 | 35 | 0 | 0 | 2.77E-14 | 69.9 |
| | GremLINE1-1_ML_211 | 100 | 36 | 35 | 0 | 0 | 2.77E-14 | 69.9 |
| | GremLINE1-1_ML_213 | 100 | 36 | 35 | 0 | 0 | 2.77E-14 | 69.9 |
| | GremLINE1-1_ML_301 | 100 | 36 | 35 | 0 | 0 | 2.77E-14 | 69.9 |
| | GremLINE1-1_ML_302 | 100 | 36 | 35 | 0 | 0 | 2.77E-14 | 69.9 |
| | GremLINE1-1_ML_303 | 100 | 36 | 35 | 0 | 0 | 2.77E-14 | 69.9 |
| | GremLINE1-1_ML_333 | 100 | 36 | 35 | 0 | 0 | 2.77E-14 | 69.9 |
| | GremLINE1-1_ML_334 | 100 | 36 | 35 | 0 | 0 | 2.77E-14 | 69.9 |
| | GremLINE1-1_ML_364 | 100 | 36 | 35 | 0 | 0 | 2.77E-14 | 69.9 |
| | GremLINE1-1_ML_365 | 100 | 36 | 35 | 0 | 0 | 2.77E-14 | 69.9 |
| | GremLINE1-1_ML_366 | 100 | 36 | 35 | 0 | 0 | 2.77E-14 | 69.9 |
| | GremLINE1-1_ML_369 | 100 | 36 | 35 | 0 | 0 | 2.77E-14 | 69.9 |
| | GremLINE1-1_ML_370 | 100 | 36 | 35 | 0 | 0 | 2.77E-14 | 69.9 |
| | GremLINE1-1_ML_402 | 100 | 36 | 35 | 0 | 0 | 2.77E-14 | 69.9 |
| | GremLINE1-1_ML_74 | 100 | 36 | 35 | 0 | 0 | 2.77E-14 | 69.9 |
| | GremLINE1-1_ML_91 | 100 | 36 | 35 | 0 | 0 | 2.77E-14 | 69.9 |
| | GremLINE1-1_ML_92 | 100 | 36 | 35 | 0 | 0 | 2.77E-14 | 69.9 |
| | GremLINE1-2_ML_120 | 100 | 36 | 35 | 0 | 0 | 2.77E-14 | 69.9 |
| | GremLINE1-2_ML_169 | 100 | 36 | 35 | 0 | 0 | 2.77E-14 | 69.9 |
| | GremLINE1-2_ML_170 | 100 | 36 | 35 | 0 | 0 | 2.77E-14 | 69.9 |
| | GremLINE1-2_ML_247 | 100 | 36 | 35 | 0 | 0 | 2.77E-14 | 69.9 |
| | GremLINE1-2_ML_367 | 100 | 36 | 35 | 0 | 0 | 2.77E-14 | 69.9 |
| | GremLINE1-3_ML_127 | 100 | 36 | 35 | 0 | 0 | 2.77E-14 | 69.9 |
| | GremLINE1-3_ML_246 | 100 | 36 | 35 | 0 | 0 | 2.77E-14 | 69.9 |
| | GremLINE1-4_ML_104 | 100 | 36 | 35 | 0 | 0 | 2.77E-14 | 69.9 |
| | GremLINE1-4_ML_105 | 100 | 36 | 35 | 0 | 0 | 2.77E-14 | 69.9 |
| | GremLINE1-4_ML_115 | 100 | 36 | 35 | 0 | 0 | 2.77E-14 | 69.9 |
| | GremLINE1-4_ML_117 | 100 | 36 | 35 | 0 | 0 | 2.77E-14 | 69.9 |
| | GremLINE1-4_ML_124 | 100 | 36 | 35 | 0 | 0 | 2.77E-14 | 69.9 |
| | GremLINE1-4_ML_204 | 100 | 36 | 35 | 0 | 0 | 2.77E-14 | 69.9 |
| | GremLINE1-4_ML_210 | 100 | 36 | 35 | 0 | 0 | 2.77E-14 | 69.9 |

| Query | Subject | % Id | Ins length | Aln length | Mm | Gap | E-value | Score |
|---|---|---|---|---|---|---|---|---|
| ***ago1Δ*_GremLINE1_1 (MC17)** | GremLINE1-4_ML_401 | 100 | 36 | 35 | 0 | 0 | 2.77E-14 | 69.9 |
| | GremLINE1-4_ML_72 | 100 | 36 | 35 | 0 | 0 | 2.77E-14 | 69.9 |
| | GremLINE1-4_ML_245 | 97.143 | 36 | 35 | 1 | 0 | 6.75E-12 | 61.9 |
| ***ago1Δ*_GremLINE1_2 (MC18)** | GremLINE1-1_ML_366 | 100 | 5957 | 5955 | 0 | 0 | 0 | 11805 |
| | GremLINE1-1_ML_402 | 100 | 5957 | 5955 | 0 | 0 | 0 | 11805 |
| | GremLINE1-1_ML_92 | 100 | 5957 | 5955 | 0 | 0 | 0 | 11805 |
| | GremLINE1-1_ML_74 | 99.916 | 5957 | 5934 | 5 | 0 | 0 | 11724 |
| | GremLINE1-1_ML_333 | 99.815 | 5957 | 5955 | 11 | 0 | 0 | 11718 |
| | GremLINE1-1_ML_365 | 99.865 | 5957 | 5934 | 8 | 0 | 0 | 11700 |
| | GremLINE1-1_ML_370 | 99.848 | 5957 | 5930 | 9 | 0 | 0 | 11684 |
| | GremLINE1-1_ML_73 | 99.815 | 5957 | 5931 | 11 | 0 | 0 | 11670 |
| | GremLINE1-1_ML_302 | 99.144 | 5957 | 5955 | 51 | 0 | 0 | 11401 |
| | GremLINE1-1_ML_334 | 99.144 | 5957 | 5955 | 51 | 0 | 0 | 11401 |
| | GremLINE1-1_ML_368 | 99.058 | 5957 | 5943 | 55 | 1 | 0 | 11329 |
| | GremLINE1-1_ML_103 | 99.04 | 5957 | 5937 | 52 | 3 | 0 | 11297 |
| | GremLINE1-2_ML_169 | 98.842 | 5957 | 5956 | 66 | 1 | 0 | 11256 |
| | GremLINE1-2_ML_367 | 98.842 | 5957 | 5956 | 66 | 1 | 0 | 11256 |
| | GremLINE1-1_ML_369 | 98.858 | 5957 | 5953 | 62 | 4 | 0 | 11234 |
| | GremLINE1-1_ML_205 | 98.757 | 5957 | 5955 | 74 | 0 | 0 | 11218 |
| | GremLINE1-1_ML_206 | 98.757 | 5957 | 5953 | 74 | 0 | 0 | 11214 |
| | GremLINE1-1_ML_102 | 98.804 | 5957 | 5934 | 71 | 0 | 0 | 11200 |
| | GremLINE1-1_ML_119 | 98.772 | 5957 | 5943 | 72 | 1 | 0 | 11194 |
| | GremLINE1-1_ML_116 | 98.607 | 5957 | 5958 | 80 | 1 | 0 | 11149 |
| | GremLINE1-2_ML_247 | 98.683 | 5957 | 5922 | 75 | 1 | 0 | 11117 |
| | GremLINE1-1_ML_364 | 98.556 | 5957 | 5954 | 83 | 1 | 0 | 11117 |
| | GremLINE1-1_ML_371 | 98.76 | 5957 | 5887 | 73 | 0 | 0 | 11091 |
| | GremLINE1-1_ML_118 | 98.568 | 5957 | 5937 | 82 | 1 | 0 | 11091 |
| | GremLINE1-1_ML_303 | 98.689 | 5957 | 5872 | 73 | 2 | 0 | 11018 |
| | GremLINE1-1_ML_404 | 98.33 | 5957 | 5927 | 85 | 1 | 0 | 10982 |
| | GremLINE1-1_ML_123 | 98.721 | 5957 | 5941 | 35 | 31 | 0 | 10949 |
| | GremLINE1-1_ML_211 | 98.499 | 5957 | 5864 | 87 | 1 | 0 | 10919 |
| | GremLINE1-1_ML_208 | 98.803 | 5957 | 5765 | 65 | 2 | 0 | 10869 |
| | GremLINE1-1_ML_207 | 98.768 | 5957 | 5761 | 71 | 0 | 0 | 10857 |
| | GremLINE1-1_ML_212 | 98.278 | 5957 | 5866 | 97 | 2 | 0 | 10816 |
| | GremLINE1-1_ML_209 | 98.595 | 5957 | 5764 | 78 | 1 | 0 | 10780 |
| | GremLINE1-1_ML_304 | 95.999 | 5957 | 5948 | 116 | 87 | 0 | 9283 |
| ***dcl1Δdcl2Δ*_GremLINE1_1 (MC8)** | GremLINE1-1_ML_103 | 100 | 534 | 534 | 0 | 0 | 0 | 1059 |
| | GremLINE1-1_ML_106 | 99.813 | 534 | 534 | 1 | 0 | 0 | 1051 |
| | GremLINE1-4_ML_105 | 99.813 | 534 | 534 | 1 | 0 | 0 | 1051 |
| | GremLINE1-1_ML_213 | 98.876 | 534 | 534 | 6 | 0 | 0 | 1011 |
| | GremLINE1-1_ML_102 | 98.689 | 534 | 534 | 7 | 0 | 0 | 1003 |
| | GremLINE1-1_ML_116 | 98.689 | 534 | 534 | 7 | 0 | 0 | 1003 |

| Query | Subject | % Id | Ins length | Aln length | Mm | Gap | E-value | Score |
|---|---|---|---|---|---|---|---|---|
| ***dcl1Δdcl2Δ*_GremLINE1_1 (MC8)** | GremLINE1-1_ML_209 | 98.689 | 534 | 534 | 7 | 0 | 0 | 1003 |
| | GremLINE1-1_ML_211 | 98.689 | 534 | 534 | 7 | 0 | 0 | 1003 |
| | GremLINE1-1_ML_301 | 98.689 | 534 | 534 | 7 | 0 | 0 | 1003 |
| | GremLINE1-1_ML_302 | 98.689 | 534 | 534 | 7 | 0 | 0 | 1003 |
| | GremLINE1-1_ML_333 | 98.689 | 534 | 534 | 7 | 0 | 0 | 1003 |
| | GremLINE1-1_ML_365 | 98.689 | 534 | 534 | 7 | 0 | 0 | 1003 |
| | GremLINE1-1_ML_366 | 98.689 | 534 | 534 | 7 | 0 | 0 | 1003 |
| | GremLINE1-1_ML_369 | 98.689 | 534 | 534 | 7 | 0 | 0 | 1003 |
| | GremLINE1-1_ML_370 | 98.689 | 534 | 534 | 7 | 0 | 0 | 1003 |
| | GremLINE1-1_ML_402 | 98.689 | 534 | 534 | 7 | 0 | 0 | 1003 |
| | GremLINE1-1_ML_74 | 98.689 | 534 | 534 | 7 | 0 | 0 | 1003 |
| | GremLINE1-1_ML_92 | 98.689 | 534 | 534 | 7 | 0 | 0 | 1003 |
| | GremLINE1-3_ML_246 | 98.689 | 534 | 534 | 7 | 0 | 0 | 1003 |
| | GremLINE1-4_ML_210 | 98.689 | 534 | 534 | 7 | 0 | 0 | 1003 |
| | GremLINE1-1_ML_249 | 98.684 | 534 | 532 | 7 | 0 | 0 | 999 |
| | GremLINE1-1_ML_118 | 98.502 | 534 | 534 | 8 | 0 | 0 | 995 |
| | GremLINE1-1_ML_364 | 98.502 | 534 | 534 | 8 | 0 | 0 | 995 |
| | GremLINE1-1_ML_91 | 98.502 | 534 | 534 | 8 | 0 | 0 | 995 |
| | GremLINE1-4_ML_104 | 98.502 | 534 | 534 | 8 | 0 | 0 | 995 |
| | GremLINE1-4_ML_117 | 98.502 | 534 | 534 | 8 | 0 | 0 | 995 |
| | GremLINE1-4_ML_124 | 98.502 | 534 | 534 | 8 | 0 | 0 | 995 |
| | GremLINE1-4_ML_245 | 98.502 | 534 | 534 | 8 | 0 | 0 | 995 |
| | GremLINE1-4_ML_401 | 98.502 | 534 | 534 | 8 | 0 | 0 | 995 |
| | GremLINE1-4_ML_72 | 98.502 | 534 | 534 | 8 | 0 | 0 | 995 |
| | GremLINE1-1_ML_303 | 98.496 | 534 | 532 | 8 | 0 | 0 | 991 |
| | GremLINE1-3_ML_127 | 98.496 | 534 | 532 | 8 | 0 | 0 | 991 |
| | GremLINE1-1_ML_73 | 98.491 | 534 | 530 | 8 | 0 | 0 | 987 |
| | GremLINE1-4_ML_115 | 98.315 | 534 | 534 | 9 | 0 | 0 | 987 |
| | GremLINE1-4_ML_204 | 98.315 | 534 | 534 | 9 | 0 | 0 | 987 |
| | GremLINE1-1_ML_300 | 98.308 | 534 | 532 | 9 | 0 | 0 | 983 |
| | GremLINE1-4_ML_248 | 98.124 | 534 | 533 | 10 | 0 | 0 | 977 |
| | GremLINE1-1_ML_368 | 98.124 | 534 | 533 | 9 | 1 | 0 | 969 |
| | GremLINE1-1_ML_212 | 97.932 | 534 | 532 | 11 | 0 | 0 | 967 |
| | GremLINE1-1_ML_119 | 97.753 | 534 | 534 | 12 | 0 | 0 | 963 |
| | GremLINE1-2_ML_120 | 97.753 | 534 | 534 | 12 | 0 | 0 | 963 |
| | GremLINE1-2_ML_169 | 97.753 | 534 | 534 | 12 | 0 | 0 | 963 |
| | GremLINE1-2_ML_247 | 97.753 | 534 | 534 | 12 | 0 | 0 | 963 |
| | GremLINE1-2_ML_367 | 97.753 | 534 | 534 | 12 | 0 | 0 | 963 |
| | GremLINE1-1_ML_206 | 97.378 | 534 | 534 | 14 | 0 | 0 | 948 |
| | GremLINE1-1_ML_207 | 97.378 | 534 | 534 | 14 | 0 | 0 | 948 |
| | GremLINE1-1_ML_208 | 97.378 | 534 | 534 | 14 | 0 | 0 | 948 |
| | GremLINE1-1_ML_403 | 97.368 | 534 | 532 | 14 | 0 | 0 | 944 |
| | GremLINE1-1_ML_334 | 97.191 | 534 | 534 | 15 | 0 | 0 | 940 |

| Query | Subject | % Id | Ins length | Aln length | Mm | Gap | E-value | Score |
|---|---|---|---|---|---|---|---|---|
| ***dcl1Δdcl2Δ*_GremLINE1_1 (MC8)** | GremLINE1-1_ML_123 | 97.575 | 534 | 536 | 7 | 4 | 0 | 932 |
| | GremLINE1-1_ML_205 | 96.816 | 534 | 534 | 17 | 0 | 0 | 924 |
| | GremLINE1-2_ML_170 | 96.816 | 534 | 534 | 17 | 0 | 0 | 924 |
| | GremLINE1-1_ML_304 | 96.84 | 534 | 538 | 11 | 5 | 0 | 894 |
| | GremLINE1-1_ML_404 | 95.113 | 534 | 532 | 12 | 1 | 0 | 866 |
| | GremLINE1-1_ML_371 | 93.843 | 534 | 536 | 31 | 1 | 0 | 795 |
| | GremLINE1-5_ML_250 | 93.084 | 534 | 535 | 31 | 3 | 0 | 749 |
| ***dcl1Δdcl2Δ*_GremLINE1_3 (MC12)** | GremLINE1-1_ML_366 | 99.95 | 5951 | 5951 | 3 | 0 | 0 | 11773 |
| | GremLINE1-1_ML_402 | 99.95 | 5951 | 5951 | 3 | 0 | 0 | 11773 |
| | GremLINE1-1_ML_92 | 99.95 | 5951 | 5950 | 3 | 0 | 0 | 11771 |
| | GremLINE1-1_ML_74 | 99.865 | 5951 | 5939 | 8 | 0 | 0 | 11710 |
| | GremLINE1-1_ML_365 | 99.815 | 5951 | 5938 | 11 | 0 | 0 | 11684 |
| | GremLINE1-1_ML_333 | 99.765 | 5951 | 5950 | 14 | 0 | 0 | 11684 |
| | GremLINE1-1_ML_370 | 99.798 | 5951 | 5935 | 12 | 0 | 0 | 11670 |
| | GremLINE1-1_ML_73 | 99.764 | 5951 | 5931 | 14 | 0 | 0 | 11646 |
| | GremLINE1-1_ML_302 | 99.093 | 5951 | 5951 | 54 | 0 | 0 | 11369 |
| | GremLINE1-1_ML_334 | 99.093 | 5951 | 5951 | 54 | 0 | 0 | 11369 |
| | GremLINE1-1_ML_368 | 99.007 | 5951 | 5943 | 58 | 1 | 0 | 11305 |
| | GremLINE1-1_ML_103 | 98.99 | 5951 | 5942 | 55 | 3 | 0 | 11284 |
| | GremLINE1-2_ML_169 | 98.791 | 5951 | 5954 | 69 | 1 | 0 | 11228 |
| | GremLINE1-2_ML_367 | 98.791 | 5951 | 5954 | 69 | 1 | 0 | 11228 |
| | GremLINE1-1_ML_369 | 98.808 | 5951 | 5954 | 65 | 4 | 0 | 11212 |
| | GremLINE1-1_ML_205 | 98.706 | 5951 | 5951 | 77 | 0 | 0 | 11186 |
| | GremLINE1-1_ML_206 | 98.706 | 5951 | 5951 | 77 | 0 | 0 | 11186 |
| | GremLINE1-1_ML_102 | 98.753 | 5951 | 5935 | 74 | 0 | 0 | 11178 |
| | GremLINE1-1_ML_119 | 98.721 | 5951 | 5944 | 75 | 1 | 0 | 11173 |
| | GremLINE1-1_ML_116 | 98.556 | 5951 | 5954 | 83 | 1 | 0 | 11117 |
| | GremLINE1-2_ML_247 | 98.633 | 5951 | 5927 | 78 | 1 | 0 | 11103 |
| | GremLINE1-1_ML_364 | 98.505 | 5951 | 5954 | 86 | 1 | 0 | 11093 |
| | GremLINE1-1_ML_118 | 98.519 | 5951 | 5940 | 85 | 1 | 0 | 11073 |
| | GremLINE1-1_ML_371 | 98.74 | 5951 | 5873 | 74 | 0 | 0 | 11056 |
| | GremLINE1-1_ML_303 | 98.639 | 5951 | 5877 | 76 | 2 | 0 | 11004 |
| | GremLINE1-1_ML_404 | 98.279 | 5951 | 5927 | 88 | 1 | 0 | 10958 |
| | GremLINE1-1_ML_123 | 98.671 | 5951 | 5943 | 38 | 31 | 0 | 10929 |
| | GremLINE1-1_ML_211 | 98.449 | 5951 | 5869 | 90 | 1 | 0 | 10905 |
| | GremLINE1-1_ML_208 | 98.752 | 5951 | 5770 | 68 | 2 | 0 | 10855 |
| | GremLINE1-1_ML_207 | 98.717 | 5951 | 5766 | 74 | 0 | 0 | 10843 |
| | GremLINE1-1_ML_212 | 98.227 | 5951 | 5866 | 100 | 2 | 0 | 10792 |
| | GremLINE1-1_ML_209 | 98.543 | 5951 | 5767 | 81 | 1 | 0 | 10762 |
| | GremLINE1-1_ML_304 | 95.948 | 5951 | 5948 | 119 | 87 | 0 | 9260 |

[1]The table shows whole insertion sequence (query) BLASTn hits against a nucleotide database containing every TE element sequence in the genome (subject). In addition, percentage of identity (% Id), whole length of the insertion (ins length) and the BLAST alignment (aln length), number of mismatches

(mm) or gaps (gap) in the alignment, and BLAST e-value and score are depicted. BLAST hits whose alignment length was < 97% of the query sequence were filtered out to ensure hits corresponding to most of the inserted sequence, and the remaining hits (alignment length ≥ 97% of the query sequence) are sorted by highest score, lowest e-value, and highest BLAST alignment length, in that order. Light green shading indicates the best scoring and longest alignments for each query.

**Table S4. Primers used in this study**[1]

| Name | Sequence | Description |
|---|---|---|
| JOHE51492 | aagagagtctcgtctcgtctcctggtgactggtgaTGCCTCAGCATTGGTACTTG | *clr4* deletion, overlap with *pyrG* |
| JOHE51493 | atgcattagatcctcactcggagacacggccaatgGTACACTGGCCATGCTATCG | *clr4* deletion, overlap with *pyrG* |
| JOHE51682 | CAGCGGTTTGGTTGAATGCC | *clr4* deletion |
| JOHE51604 | TCCTCCTCCGCAACAACAAC | *clr4* deletion |
| JOHE52150 | GCGAAACGAACTCAGTCATGG | *clr4* deletion, WT allele PCR |
| JOHE52151 | TGTGCCTTTGGGAATGTCTTG | *clr4* deletion, WT allele PCR |
| JOHE51684 | GGAGCTTGGTGAGGGTTAGC | *clr4* deletion, 3´spanning MUT allele PCR |
| JOHE51510 | CCTTTGTTTGTGTGCTCAAGGT | *clr4* deletion, 3´spanning MUT allele PCR |
| JOHE51718 | GTCTTGTCTCATGCAGACT | GremLINE1 qPCR |
| JOHE51719 | GCACCAAGAGAGAGACGAGC | GremLINE1 qPCR |
| JOHE51666 | CAGCACCAACAACAGTGACG | *vma1* qPCR |
| JOHE51668 | ACCTTGGTGCTCGTCTTGCC | *vma1* qPCR |
| JOHE52223 | AATCGCAACGGAATTCCTGACCGGAC | *fkbA* amplification and Sanger sequencing |
| JOHE52224 | CTCAAGACGGACGAGCCTCA | *fkbA* amplification and Sanger sequencing |
| JOHE51780 | AAAACGCGAAACAGCAAACG | *fkbA* Sanger sequencing |
| JOHE51782 | AGGCTACCTTGAACTTTGAAG | *fkbA* Sanger sequencing |
| JOHE52501 | TTGGTGATCGTGTGGATGC | TE insertion Sanger sequencing |
| JOHE52502 | GTTACACCAAGAAGCAGAAGG | TE insertion Sanger sequencing |
| JOHE52503 | TCTGGTGTTTCTGGCTCTGG | TE insertion Sanger sequencing |
| JOHE52504 | GCTAGTATTGGTCCTGGTAGT | TE insertion Sanger sequencing |
| JOHE52505 | GATGGTGATATGCTGACGGCA | TE insertion Sanger sequencing |
| JOHE52506 | GGCGTTTGAGCCTTTGCTTC | TE insertion Sanger sequencing |
| JOHE52507 | CCTTGGTTCTCTTTCCCTGAG | TE insertion Sanger sequencing |
| JOHE52508 | TGGCTATGAACATGAGTGTGG | TE insertion Sanger sequencing |
| JOHE52509 | CACTCGTTGGCTCGAACAGG | TE insertion Sanger sequencing |
| JOHE52510 | GGCTAACTCGTTACTGTTGTCC | TE insertion Sanger sequencing |
| JOHE52582 | CCACACTCATGTTCATAGCCACA | TE insertion Sanger sequencing |
| JOHE52583 | CCTGTTCGAGCCAACGAGTG | TE insertion Sanger sequencing |
| JOHE52584 | GGACAACAGTAACGAGTTAGCC | TE insertion Sanger sequencing |
| JOHE52585 | CTCATCATAGACGCTCAAAGGC | TE insertion Sanger sequencing |
| JOHE52586 | GCTACTCGTTCTTGGTGCTATG | TE insertion Sanger sequencing |

[1]Every primer shows systematic name, nucleotide sequence (in 5’-3’ direction), and a brief description explaining its use. Lowercase nucleotides overlap with the *pyrG* marker for *clr4* deletion.

**Dataset S1 (separate file).** RNA interference (RNAi) components, histone-lysine (KMT) and DNA 5mC methyltransferases (DNMT), and histone demethylases (KDM) in fungi. Homologs of RNAi components and chromatin modifying enzymes (left, Protein) identified in each given species (Species, and Source for genome references). Each identified homolog (Subject) shows PSI-BLAST percentage of identity (% Id), E-value, score, and its matching query sequence (Query). Next, percentage of identity, E-value, and score of a reciprocal BLAST against that same query (Recip). For every homolog, protein length, InterPro protein domains (IPR and Description), their start and end positions, and their prediction E-value is also shown. Subjects (identified homologs) from JGI/Broad Institute show a JGI protein ID (or FungiDB gene ID if available), whereas those from *Absidia glauca* show an Ensembl gene ID. Queries (and reciprocal hits) show their species name as a three-letter abbreviation (or four-letter for disambiguation, first letter from their genus and two/three first letters from their species epithet), the protein name, and their identifier separated by an underscore. For both subjects and queries, known and experimentally curated IDs may show their gene name/symbol at the end after a dash character.

**Dataset S2 (separate file).** Fungal RNAi, KMT, DNMT, and KDM enzyme protein domain architecture. Full-length, scaled protein sequences of every identified protein homologs and their predicted, color-coded protein domains (rectangles) are shown. Identified homologs from JGI/Broad Institute show a JGI protein ID (or FungiDB gene ID if available), whereas those from *Absidia glauca* show an Ensembl gene ID, and experimentally curated proteins may show their gene name/symbol instead of their ID.

**Dataset S3 (separate file).** Major transposable element families and subfamilies, their ORFs, and protein domain configurations. The full-length, scaled nucleotide sequences of the major transposable element families and subfamilies identified in M. lusitanicus are depicted. Each family shows ORF prediction (gray arrowed blocks), as well as encoded InterPro protein domains (color-coded rectangles).

**Dataset S4 (separate file)**. Fungal RNAi, KMT, DNMT, and KDM enzyme protein FASTA sequences.
