## Supplementary material for "Heterochromatin and RNAi act independently to ensure genome stability in Mucorales human fungal pathogens": Fig. S1

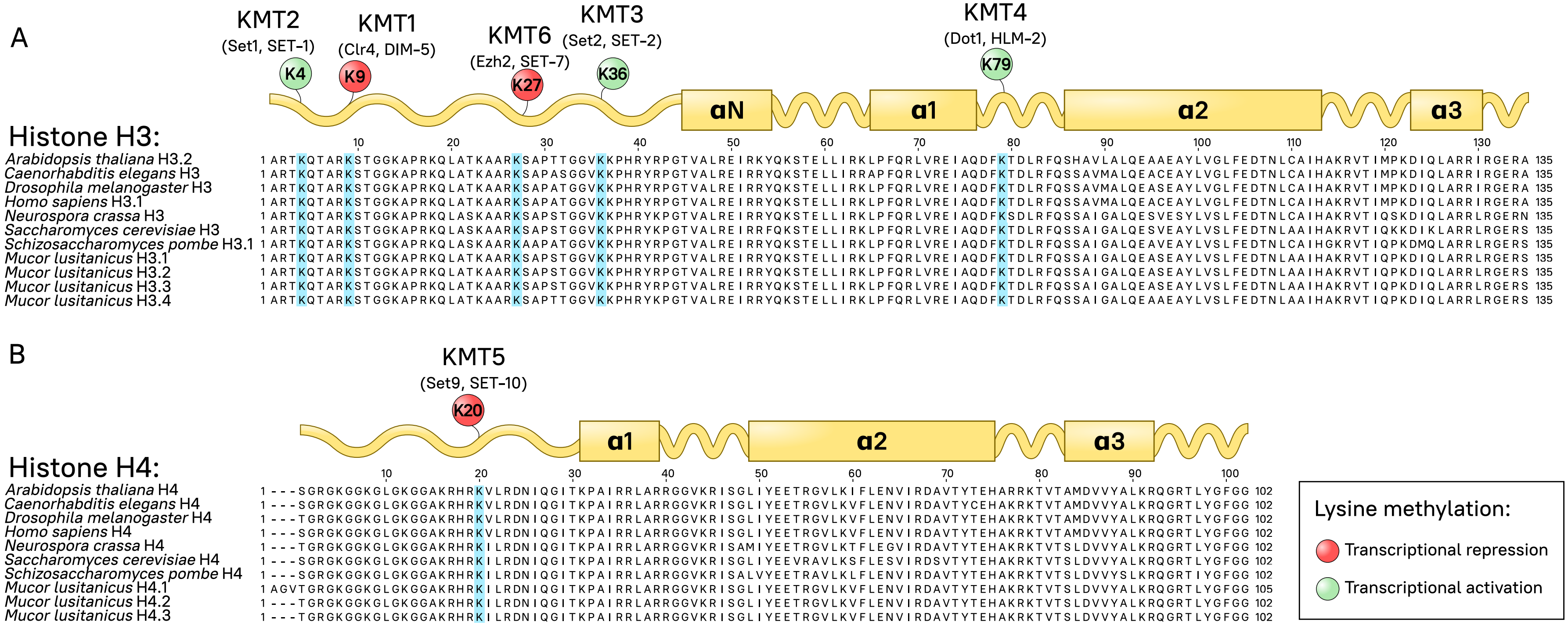


**Fig. S1.** *Mucor* histones display conserved residues for histone-lysine methyltransferase (KMT) activity. Histone H3 **(A)** and H4 **(B)** protein sequence alignment of several model organisms and *M. lusitanicus* is shown. Key lysine residues are highlighted in the alignment (cyan). Methyltransferases involved in histone-lysine methylation and the residue they target are depicted in a schematic representation of each histone protein, showing the N-terminal tail and the histone fold domain comprised of several α-helices (rectangles). Transcriptional repressive (red) and activating (green) modifications are displayed.
