## Supplementary material for "Heterochromatin and RNAi act independently to ensure genome stability in Mucorales human fungal pathogens": Fig. S2

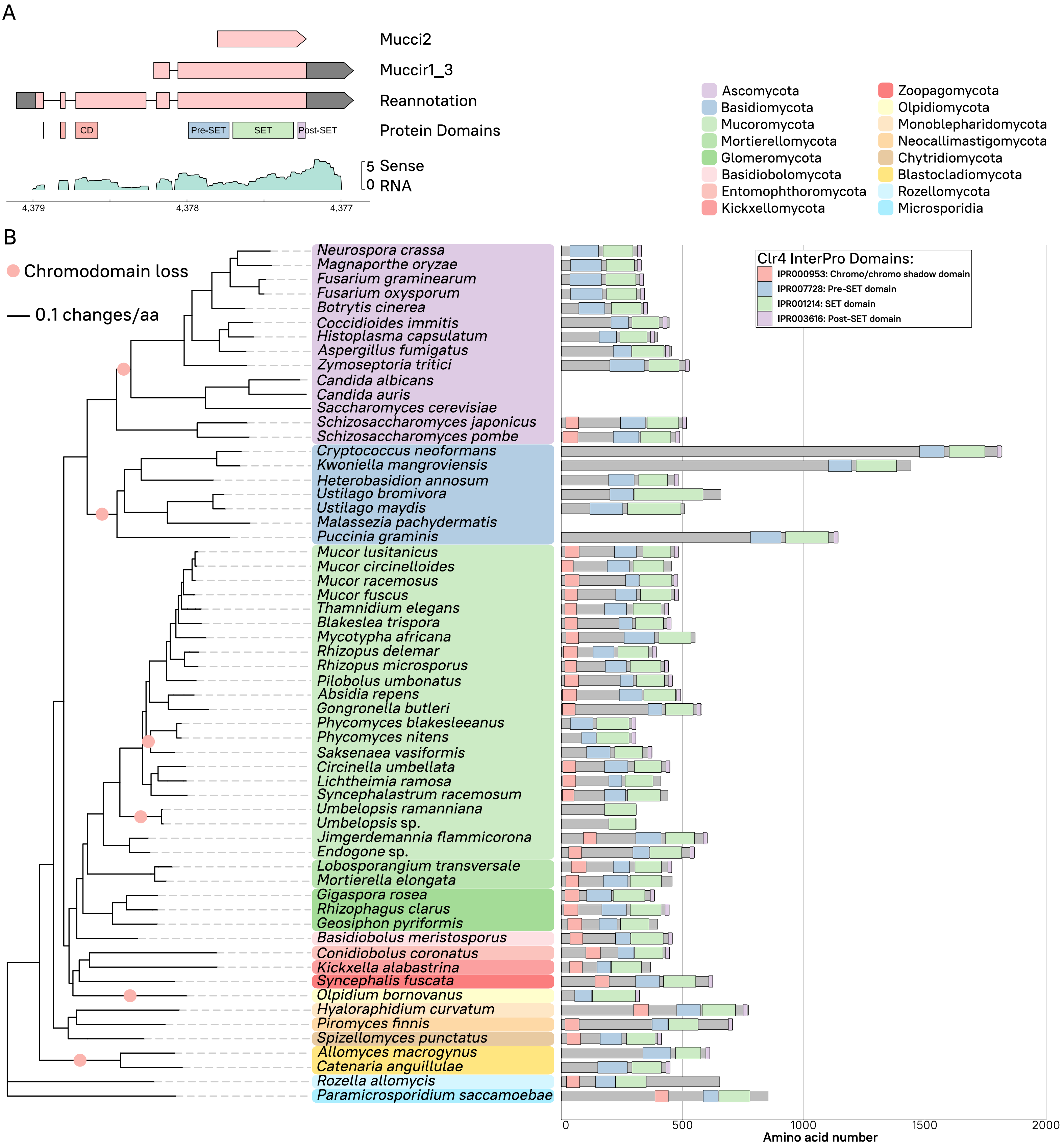


**Fig. S2**. Mucoromycota and other early-diverging fungi harbor an N-terminal chromodomain in Clr4. **(A)** Genomic plot of the *clr4* locus in *M. lusitanicus* MU402. Three different *clr4* gene models are shown (red arrowed blocks, dark gray for UTR regions), including the last reannotation according to our transcriptomic data. Position of Clr4 protein domains encoded in the DNA sequence is shown below gene models for reference. Sense transcripts are plotted as normalized $log_2$ counts per million (CPM) values (green). **(B)** Clr4 methyltransferases of 60 fungal species clustered by phylogenomic relationship (left, tree), color-coded by different fungal phyla. The full-length, scaled protein sequence is shown along its predicted protein domains (legend). Chromodomain loss is pointed as a red circle in the phylogenomic tree. Phylogenomic distance (black solid line) is shown as changes per amino acid.
