## Supplementary material for "Heterochromatin and RNAi act independently to ensure genome stability in Mucorales human fungal pathogens": Fig. S3

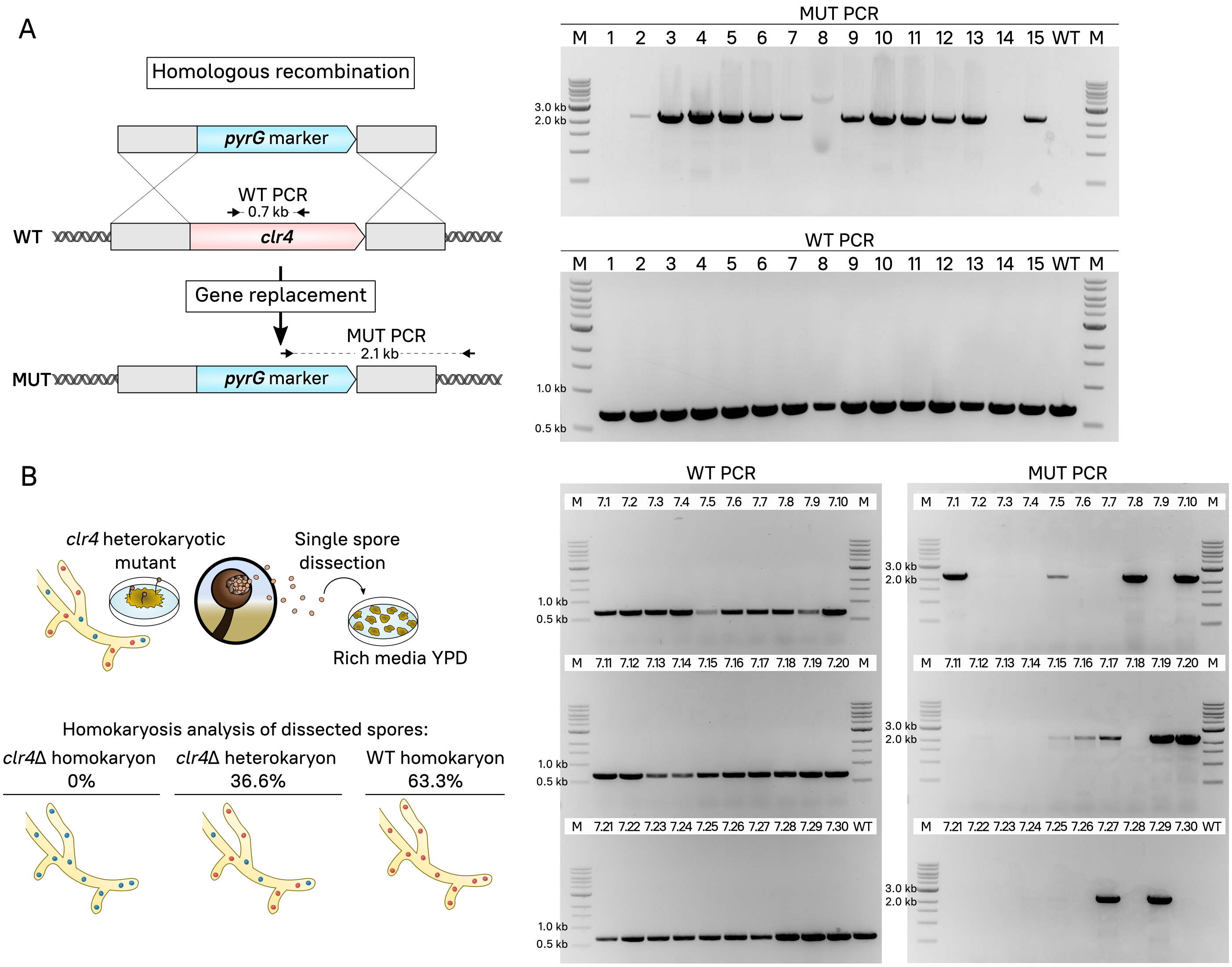


**Fig. S3.** *clr4* homokaryotic deletion results in lethality. **(A)** Schematic strategy followed for *clr4* gene deletion, replacing it with the *pyrG* marker. Primer binding sites for wild-type allele (WT PCR) and 3' mutant allele junction (MUT PCR) are shown. Gel electrophoresis showing PCR products from WT and MUT PCR in 15 transformants (1-15) and wild-type strain (WT). **(B)** Procedure to obtain homokaryotic mutants by single spore dissection from the heterokaryotic mutant #7. WT and MUT PCRs of 30 dissected spores (from 7.1 to 7.30). Spore genotype (mutant homokaryon, heterokaryon, and wild-type homokaryon) is summarized as percentage of total spores. M indicates DNA ladder in electrophoresis gels.
