## Supplementary material for "Heterochromatin and RNAi act independently to ensure genome stability in Mucorales human fungal pathogens": Fig. S4

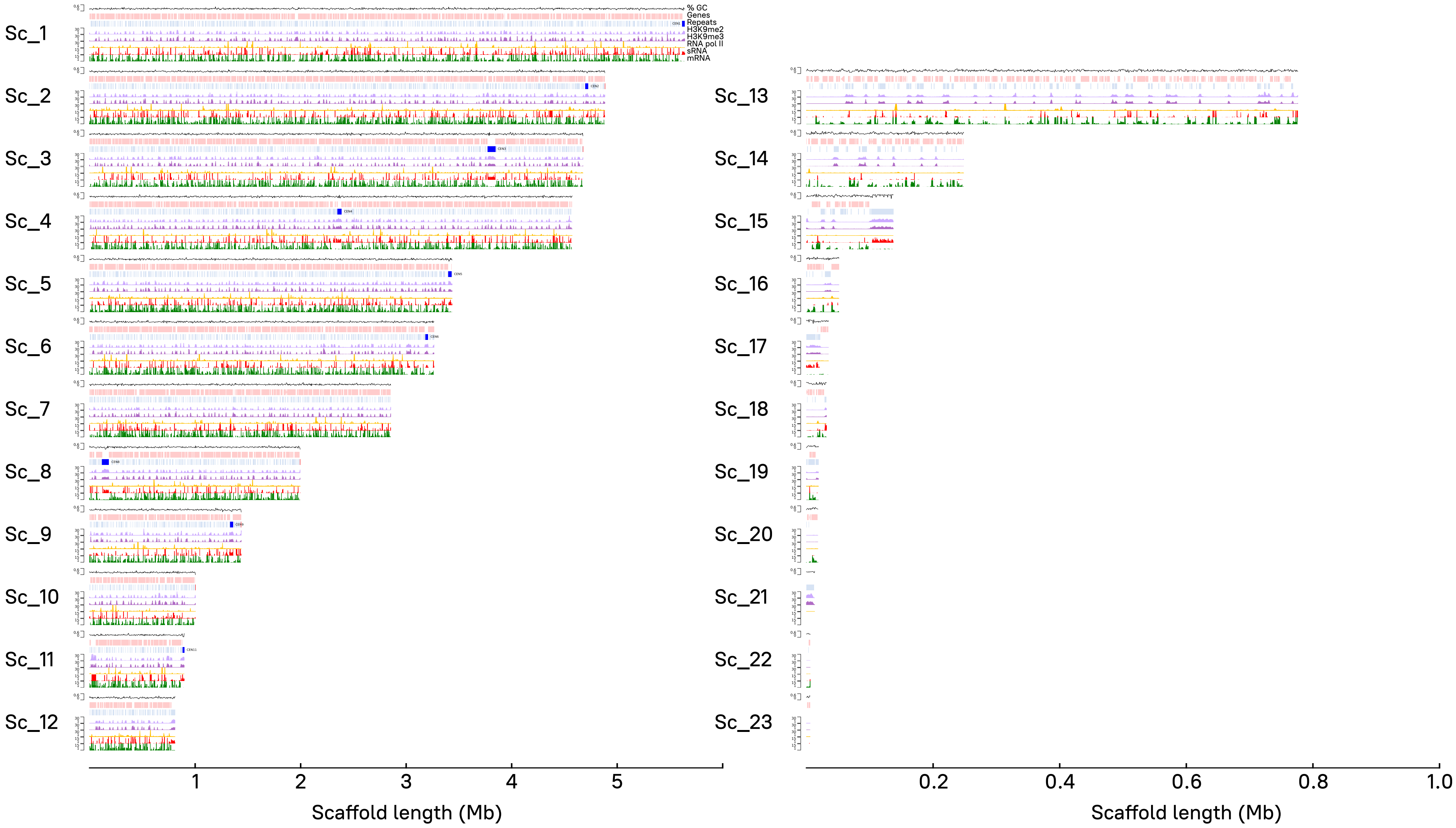


**Fig. S4.** H3K9me2, -me3, RNA pol II, sRNA, and mRNA genome-wide distribution in *M. lusitanicus*. Read coverage normalized as $log_2$ CPM is shown as immunoprecipitated (IP) DNA/Input DNA ratio for H3K9me2 (light purple) and -me3 (purple) and RNA polymerase II (RNA pol II, gold); messenger RNA (mRNA, green) and sRNA (red) coverage as $log_2$ CPM. Genes (light red) and repeated sequences (light blue) including pericentric regions (bright blue) are displayed, as well as GC content coverage as a percentage (black line). Every scaffold (1-23) and its length are plotted, but note that larger scaffolds comprising ≥ 95 % of the genome (1-12, left) are scaled differently than shorter scaffolds (13-23, right).
