## Supplementary material for "Heterochromatin and RNAi act independently to ensure genome stability in Mucorales human fungal pathogens": Fig. S5

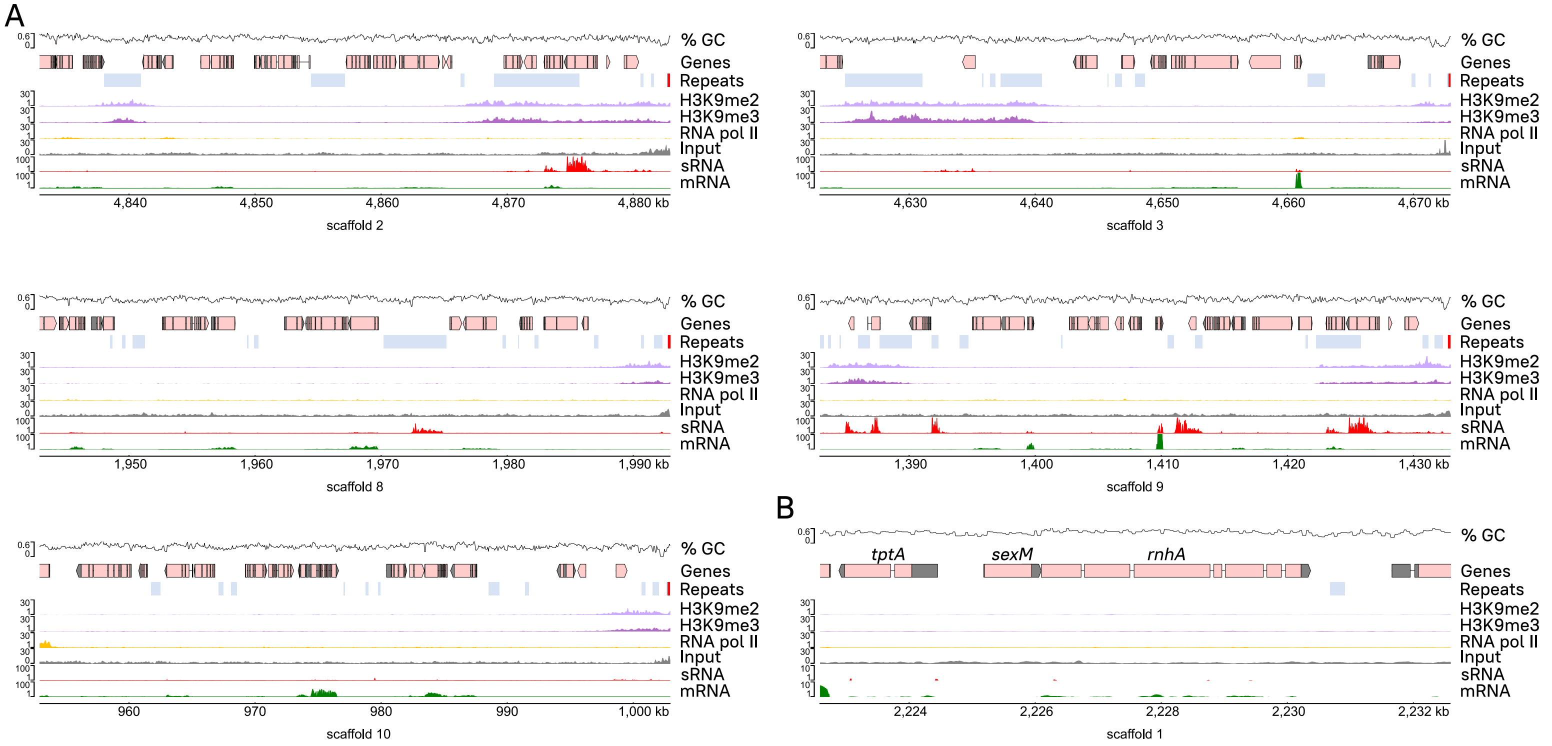


**Fig. S5**. H3K9me and sRNA enrichment in telomeric and sex-determining regions. H3K9me2 (light purple), -me3 (dark purple), and RNA polymerase II (RNA pol II, gold) IP/Input ratio, as well as Input control coverage for reference, are shown across 50-kb telomeric regions identified **(A)** and the *sex* locus **(B)**, and normalized as previously described. Transcripts (mRNA, green) and sRNA (red) coverages are also depicted. Genes (light red) and repeated sequences (light blue) including telomeric repeats (bright red) are displayed, as well as GC content coverage as a percentage.
