## Supplementary material for "Heterochromatin and RNAi act independently to ensure genome stability in Mucorales human fungal pathogens": Fig. S6

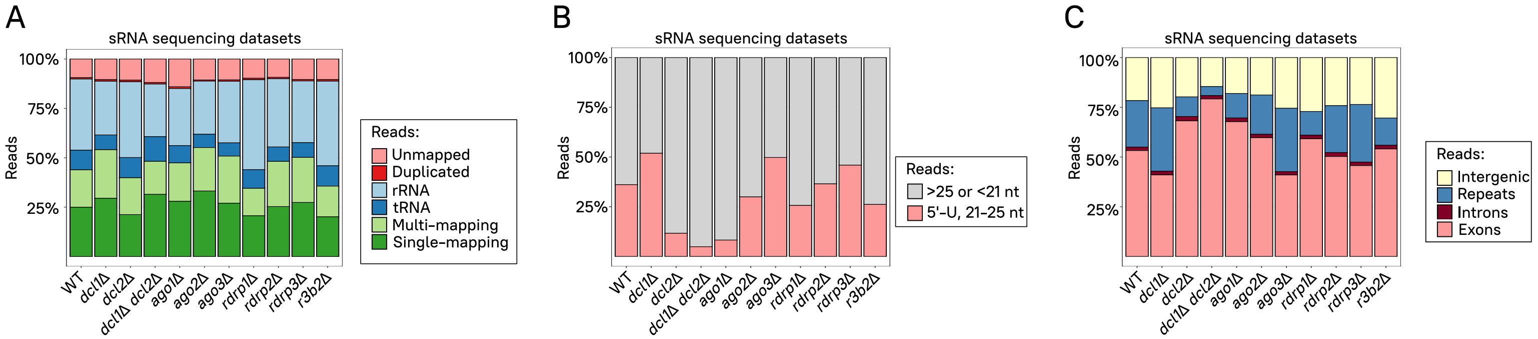


**Fig. S6.** Canonical sRNAs targeting primarily exonic sequences are generated by Dicer and Argonaute activity. **(A)** The percentage of unmapped, duplicated, rRNA or tRNA mapping, multi- or single-mapping reads to the genome in RNAi mutant and wild-type strain sRNA samples is depicted. **(B)** The percentage of aligned reads with a size ranging from 21 to 25 and bearing a 5'-uracil in the same samples is shown. **(C)** The percentage of aligned reads mapping to intergenic, repeated, intronic, and exonic features in the same samples is plotted.
