## Supplementary material for "Heterochromatin and RNAi act independently to ensure genome stability in Mucorales human fungal pathogens": Fig. S7

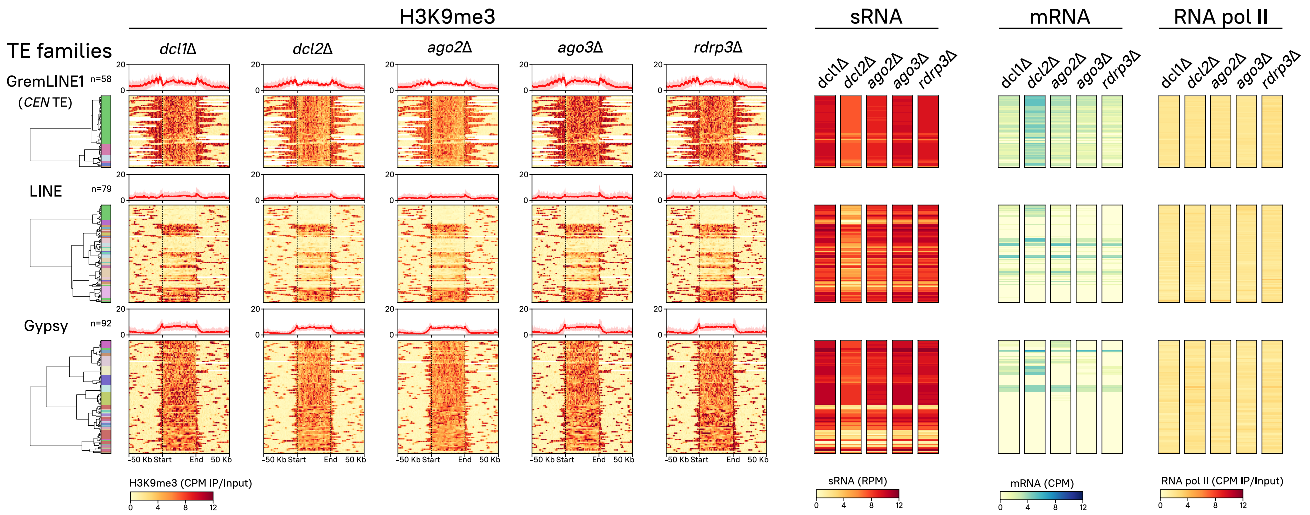


**Fig. S7**. H3K9me3, sRNA, mRNA, and RNA pol II enrichment across RNA transposons in non-essential RNAi paralog mutants. GremLINE1, other LINE, and Gypsy RNA transposon copies are clustered as in Fig. 3. Heatmaps depicting H3K9me3, sRNA, mRNA, and RNA polymerase II values in the wildtype and mutants in non-essential RNAi enzymes are plotted. H3K9me3 enrichment values are shown as IP/input DNA ratio of normalized $\log_2$ CPM from start to end of each copy (divided in 250 bins) and 50 kb upstream and downstream; as well as the mean value of all the copies in each family shown in a profile plot on top of each heatmap. sRNA and mRNA values are normalized to $\log_2$ reads per million (RPM) and CPM, respectively. RNA pol II enrichment values are also displayed as IP/Input ratio of normalized $\log_2$ CPM.
