## Supplementary material for "Heterochromatin and RNAi act independently to ensure genome stability in Mucorales human fungal pathogens": Fig. S8

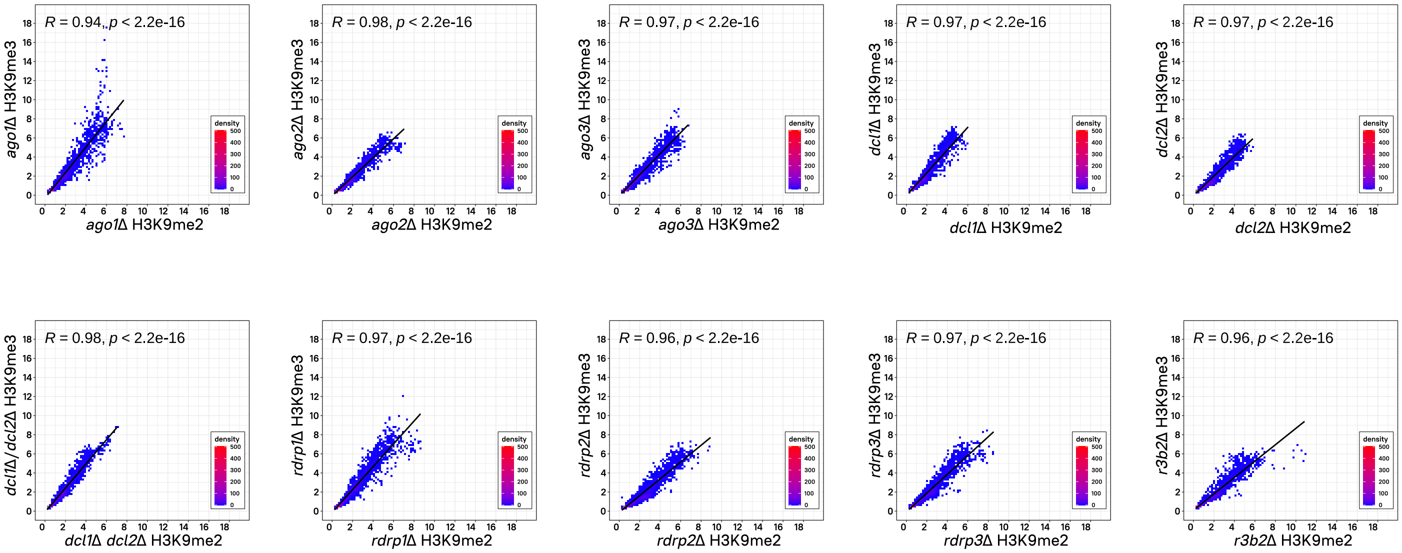


**Fig. S8.** H3K9me2 and -me3 distribution is significantly correlated in RNAi-deficient mutants. Pearson's correlation coefficient (R, dark line) and its *p*-value shown in a dot plot of H3K9me3 (y-axis) and -me2 (x-axis) average $\log_2$ CPM enrichment values in 10-kb regions representing the whole genome of *M. lusitanicus*. Overlapping dots are shown as blue-to-red colored corresponding to plotting density.
