## Supplementary material for "Heterochromatin and RNAi act independently to ensure genome stability in Mucorales human fungal pathogens": Fig. S9

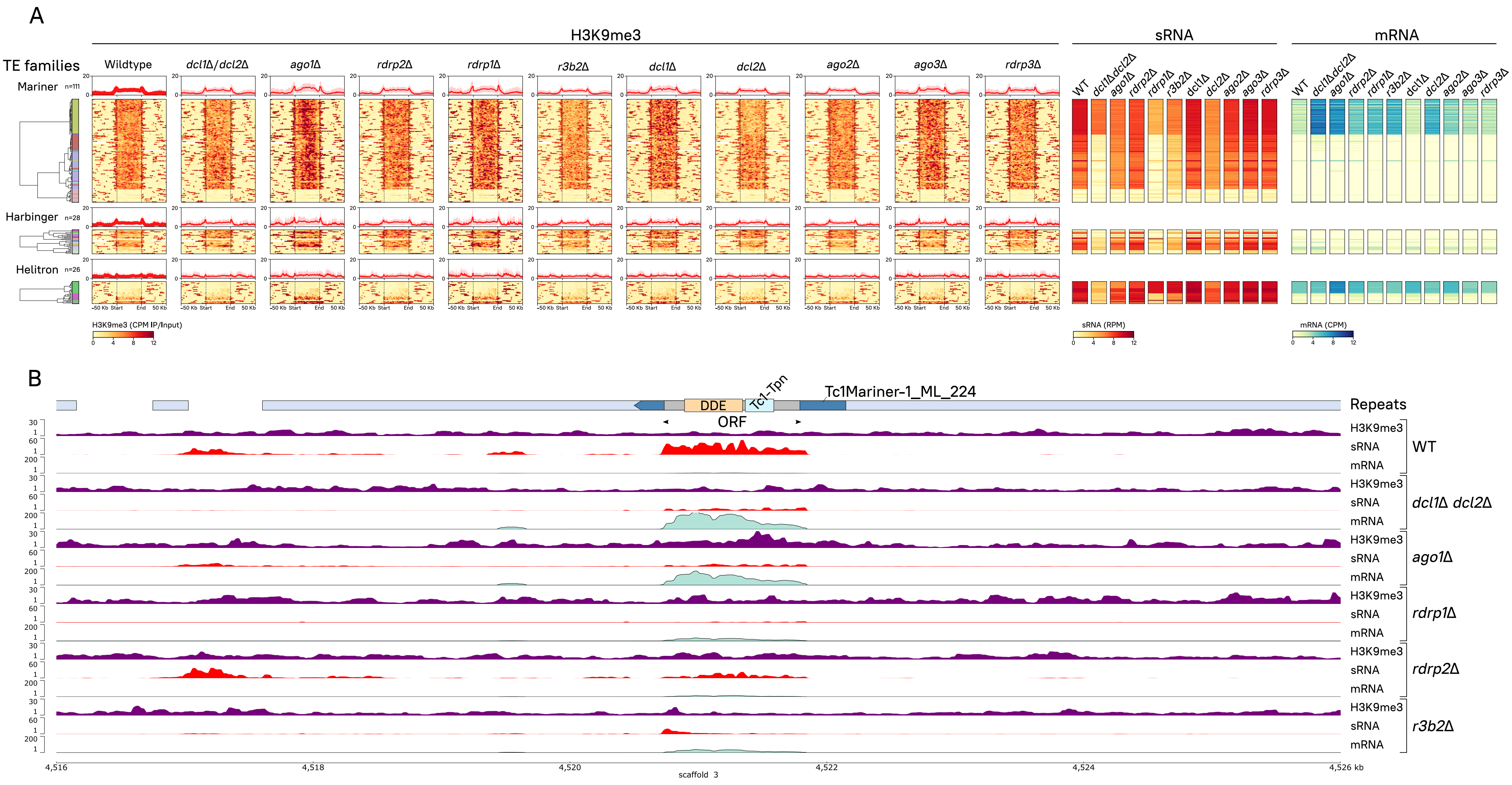


**Fig. S9.** DNA transposons are also targeted by both RNAi and H3K9me-based heterochromatin. **(A)** Every copy of Tc1/Mariner (n=111), PIF/Harbinger LINE (n=28), and RC/Helitron (n=26) DNA or Class II transposons identified in the genome are clustered according to their subfamily, H3K9me3, sRNA, and mRNA values in the wild-type *M. lusitanicus* strain. Heatmaps depicting H3K9me3, sRNA, mRNA, and RNA polymerase II values in wild-type and all RNAi mutant strains –essential and non-essential for RNAi– are plotted (see plot name above). As in Fig. 3, each row represents a single DNA transposon copy, and data values are aligned across all heatmaps to facilitate comparison. From left to right, a color-coded dendrogram indicating each element subfamily. Next, H3K9me3 enrichment values are shown as IP/input ratio of $\log_2$ CPM, from the start to the end of each copy and 50-kb upstream and downstream regions. The average of all the copies in each family is plotted in an H3K9me3 profile on top of each heatmap (bold red line shows the average and faded red background shows the standard deviation). Next heatmap to the right shows sRNA values normalized to $\log_2$ RPM, followed by a sense transcript heatmap (mRNA) in $\log_2$ CPM. The last heatmap on the right depicts RNA pol II enrichment values as IP/Input ratio of $\log_2$ CPM. **(B)** Tc1/Mariner-1 (blue arrowed block) representative genomic plot showing H3K9me3, sRNA, and mRNA data normalized as in **(A)** for canonical (*dcl1*Δ *dcl2*Δ, *ago1*Δ, *rdrp2*Δ, and *rdrp1*Δ) and alternative (*rdrp1*Δ and *r3b2*Δ) RNAi-deficient mutants. Transposition-related protein domains (colored rectangles) encoded by Tc1/Mariner-1_ML_224 ORF sequence (gray rectangle) and their position is shown. Other structural repeats (light blue rectangles) are plotted for reference.
