## Supplementary material for "Heterochromatin and RNAi act independently to ensure genome stability in Mucorales human fungal pathogens": Fig. S10

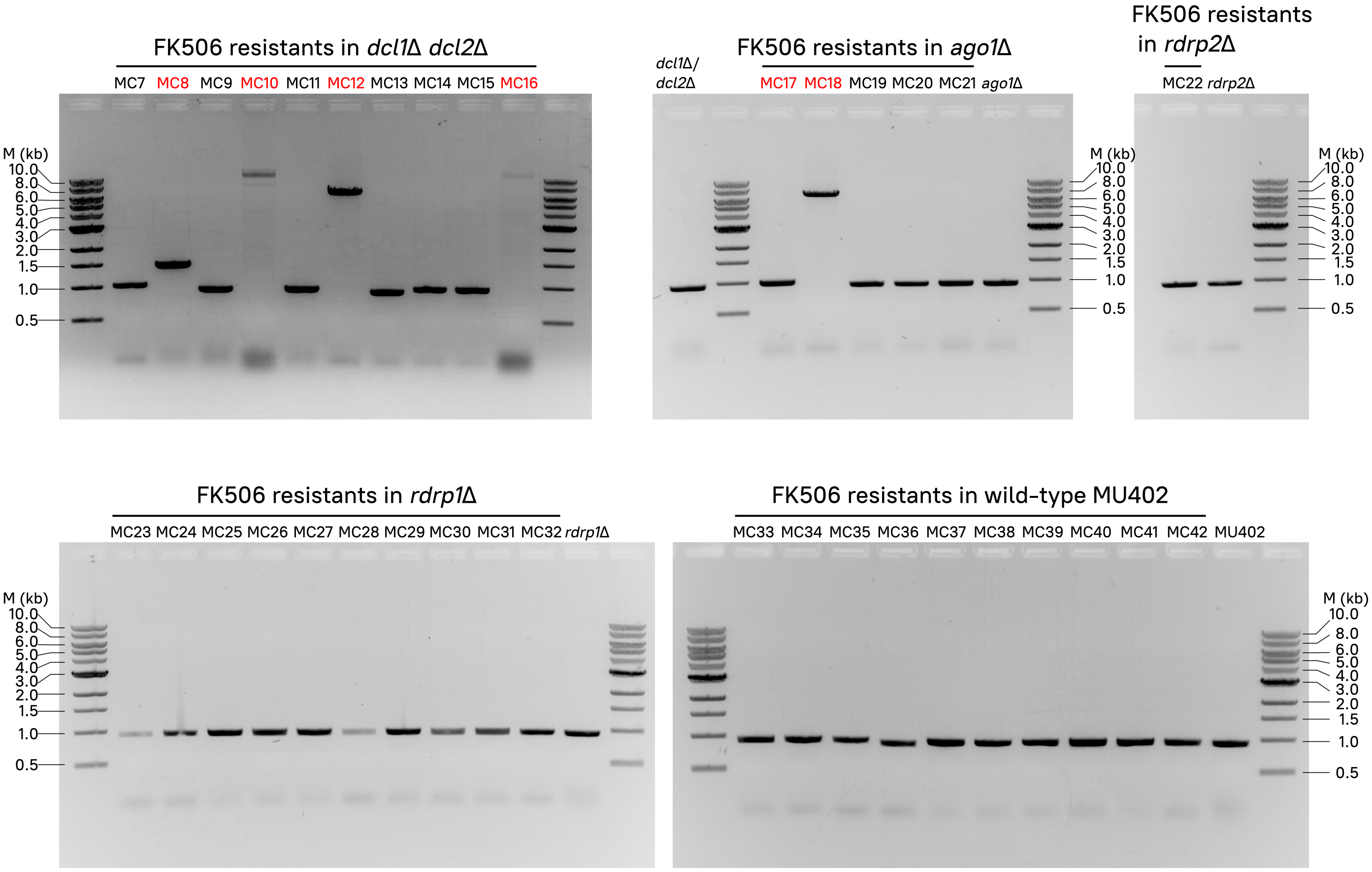


**Fig. S10.** Loss of canonical Dicer or Ago activities leads to large insertions and FK506 target disruption. The FKBP12-encoding *fkbA* locus was PCR-amplified in FK506-resistant isolates in different RNAi-deficient backgrounds (shown on top of each electrophoresis gel picture). *fkbA* sequences containing insertions (also confirmed by Sanger-sequencing) are highlighted in red, and were only found in *dcl1*Δ *dcl2*Δ and *ago1*Δ mutant strains. RNAi-deficient and wild-type strains not exposed to FK506 were also analyzed and included as FK506-sensitive controls. Lanes marked as M indicate DNA ladder and expected fragment sizes.
