## Supplementary material for "Heterochromatin and RNAi act independently to ensure genome stability in Mucorales human fungal pathogens": Dataset S2

**Dataset S2.** Fungal RNAi, KMT, DNMT, and KDM enzyme protein domain architecture. Full-length, scaled protein sequences of every identified protein homologs and their predicted, color-coded protein domains (rectangles) are shown. Identified homologs from JGI/Broad Institute show a JGI protein ID (or FungiDB gene ID if available), whereas those from Absidia glauca show an Ensembl gene ID, and experimentally curated proteins may show their gene name/symbol instead of their ID.

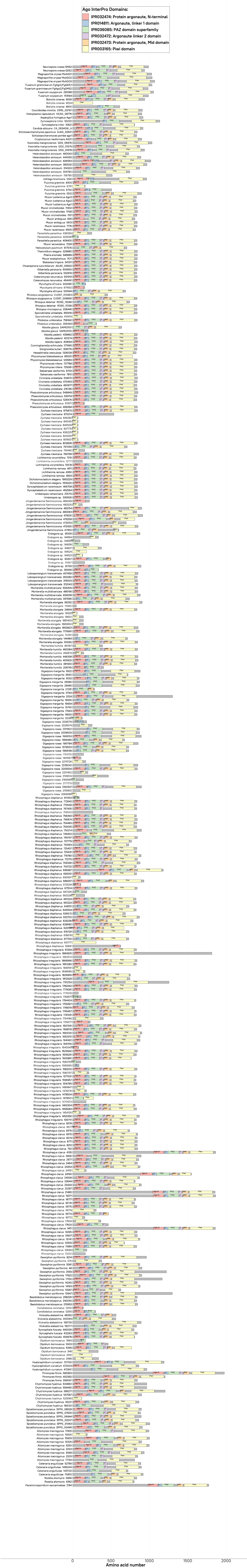

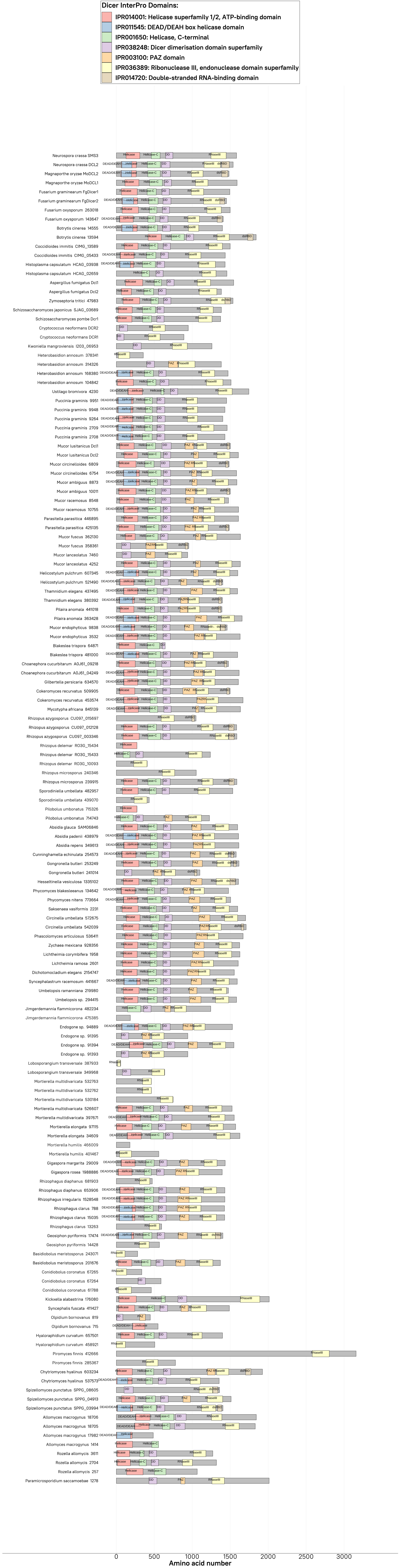

Dicer InterPro Domains:
IPR014001: Helicase superfamily 1/2, ATP-binding domain
IPR011545: DEAD/DEAH box helicase domain
IPR001650: Helicase, C-terminal
IPR038248: Dicer dimerisation domain superfamily
IPR003100: PAZ domain
IPR036389: Ribonuclease III, endonuclease domain superfamily
IPR014720: Double-stranded RNA-binding domain
Neurospora crassa SMS3
Neurospora crassa DCL2
Magnaporthe oryzae MoDCL2
Magnaporthe oryzae MoDCL1
Fusarium graminearum FgDicer1
Fusarium graminearum FgDicer2
Fusarium oxysporum 263018
Fusarium oxysporum 143647
Botrytis cinerea 14555
Botrytis cinerea 13594
Coccidioides immitis CIMG_13589
Coccidioides immitis CIMG_05433
Histoplasma capsulatum HCAG_03938
Histoplasma capsulatum HCAG_02659
Aspergillus fumigatus Dcl1
Aspergillus fumigatus Dcl2
Zymoseptoria tritici 47983
Schizosaccharomyces japonicus SJAG_03689
Schizosaccharomyces pombe Dcr1
Cryptococcus neoformans DCR2
Cryptococcus neoformans DCR1
Kwoniella mangroviensis I203_06953
Heterobasidion annosum 378341
Heterobasidion annosum 314326
Heterobasidion annosum 168380
Heterobasidion annosum 104842
Ustilago bromivora 4230
Puccinia graminis 9951
Puccinia graminis 9948
Puccinia graminis 9264
Puccinia graminis 2709
Puccinia graminis 2708
Mucor lusitanicus Dcl1
Mucor lusitanicus Dcl2
Mucor circinelloides 6809
Mucor circinelloides 6754
Mucor ambiguus 8873
Mucor ambiguus 10011
Mucor racemosus 8548
Mucor racemosus 10755
Parasitella parasitica 446895
Parasitella parasitica 425135
Mucor fuscus 362130
Mucor fuscus 358361
Mucor lanceolatus 7460
Mucor lanceolatus 4252
Helicostylum pulchrum 607345
Helicostylum pulchrum 521490
Thamnidium elegans 437495
Thamnidium elegans 380392
Pilaira anomala 441018
Pilaira anomala 363428
Mucor endophyticus 9838
Mucor endophyticus 3532
Blakeslea trispora 64871
Blakeslea trispora 481000
Choanephora cucurbitarum A0J61_09218
Choanephora cucurbitarum A0J61_04249
Gilbertella persicaria 643670
Cokeromyces recurvatus 509905
Cokeromyces recurvatus 453574
Mycotypha africana 845139
Rhizopus azygosporus CU097_015697
Rhizopus azygosporus CU097_012128
Rhizopus azygosporus CU097_003346
Rhizopus delemar RO3G_15434
Rhizopus delemar RO3G_15433
Rhizopus delemar RO3G_10093
Rhizopus microsporus 240346
Rhizopus microsporus 239915
Sporodiniella umbellata 482957
Sporodiniella umbellata 439070
Pilobolus umbonatus 715326
Pilobolus umbonatus 714743
Absidia glauca SAM06846
Absidia padenii 438979
Absidia repens 349613
Cunninghamella echinulata 254573
Gongronella butleri 253249
Gongronella butleri 241014
Hesseltinella vesiculosa 1335102
Phycomyces blakesleeanus 134642
Phycomyces nitens 773664
Saksenaea vasiformis 2231
Circinella umbellata 572675
Circinella umbellata 542039
Phascolomyces articulosus 536411
Zychaea mexicana 928356
Lichtheimia corymbifera 1958
Lichtheimia ramosa 2601
Dichotomocladium elegans 2154747
Syncephalastrum racemosum 441667
Umbelopsis ramanniana 219980
Umbelopsis sp. 294415
Jimgerdemannia flammicorona 482234
Jimgerdemannia flammicorona 475385
Endogone sp. 94889
Endogone sp. 91395
Endogone sp. 91394
Endogone sp. 91393
Lobosporangium transversale 387933
Lobosporangium transversale 349968
Mortierella multidivaricata 532763
Mortierella multidivaricata 532762
Mortierella multidivaricata 530184
Mortierella multidivaricata 526607
Mortierella multidivaricata 397671
Mortierella elongata 97115
Mortierella elongata 34609
Mortierella humilis 466009
Mortierella humilis 401467
Gigaspora margarita 29009
Gigaspora rosea 1988886
Rhizophagus diaphanus 681903
Rhizophagus diaphanus 653906
Rhizophagus irregularis 1528548
Rhizophagus clarus 788
Rhizophagus clarus 15035
Rhizophagus clarus 13263
Geosiphon pyriformis 17474
Geosiphon pyriformis 14428
Basidiobolus meristosporus 243071
Basidiobolus meristosporus 201676
Conidiobolus coronatus 67265
Conidiobolus coronatus 67264
Conidiobolus coronatus 61788
Kickxella alabastrina 176080
Syncephalis fuscata 411427
Olpidium bornovanus 819
Olpidium bornovanus 715
Hyaloraphidium curvatum 657501
Hyaloraphidium curvatum 458921
Piromyces finnis 412666
Piromyces finnis 285367
Chytriomyces hyalinus 603234
Chytriomyces hyalinus 537573
Spizellomyces punctatus SPPG_08605
Spizellomyces punctatus SPPG_04913
Spizellomyces punctatus SPPG_03994
Allomyces macrogynus 18706
Allomyces macrogynus 18705
Allomyces macrogynus 17982
Allomyces macrogynus 1414
Rozella allomycis 3611
Rozella allomycis 2704
Rozella allomycis 257
Paramicrosporidium saccamoebae 1278
0
500
1000
1500
2000
2500
3000
Amino acid number

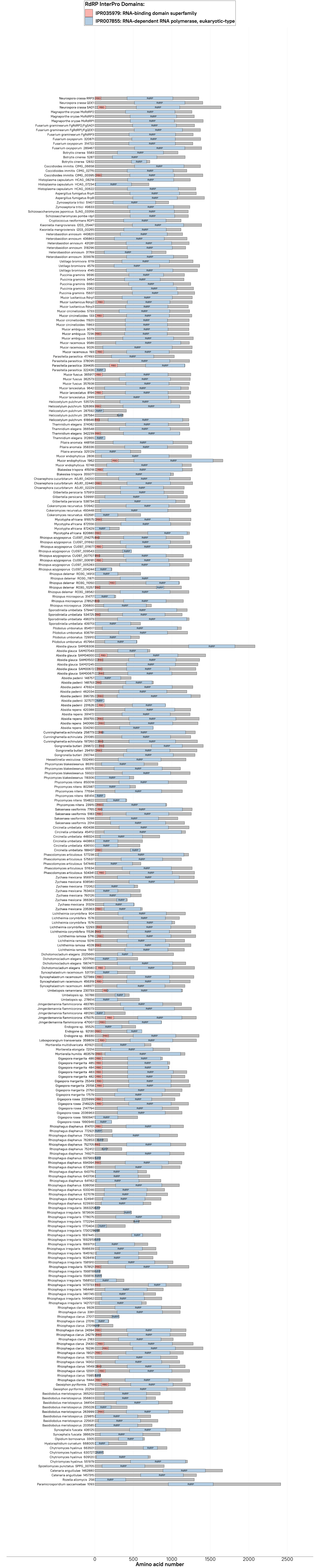
RdRP InterPro Domains:
IPR035979: RNA-binding domain superfamily
IPR007855: RNA-dependent RNA polymerase, eukaryotic-type
0
500
1000
1500
2000
2500
Amino acid number

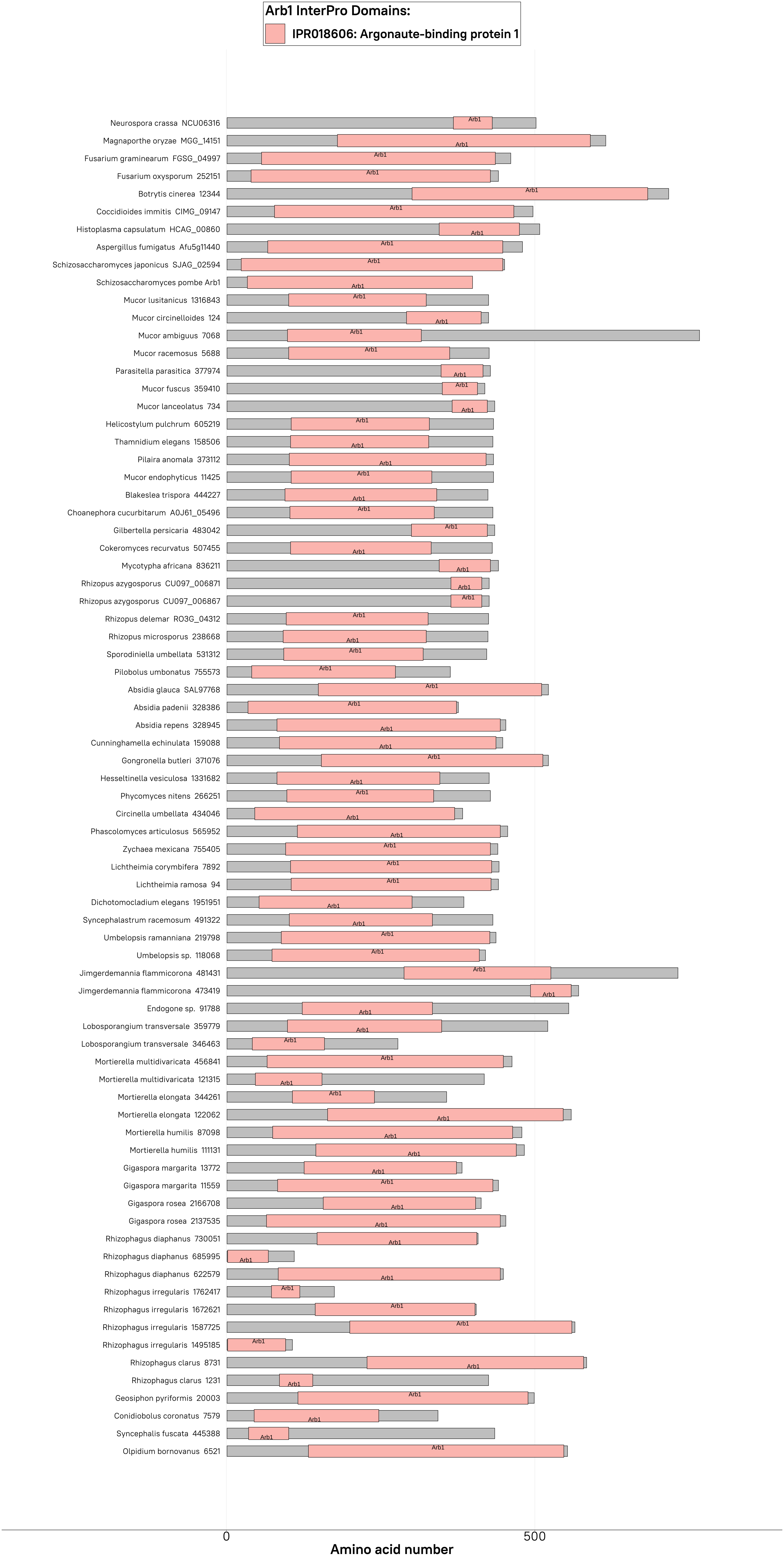
Arb1 InterPro Domains:
IPR018606: Argonaute-binding protein 1
Neurospora crassa NCU06316
Magnaporthe oryzae MGG_14151
Fusarium graminearum FGSG_04997
Fusarium oxysporum 252151
Botrytis cinerea 12344
Coccidioides immitis CIMG_09147
Histoplasma capsulatum HCAG_00860
Aspergillus fumigatus Afu5g11440
Schizosaccharomyces japonicus SJAG_02594
Schizosaccharomyces pombe Arb1
Mucor lusitanicus 1316843
Mucor circinelloides 124
Mucor ambiguus 7068
Mucor racemosus 5688
Parasitella parasitica 377974
Mucor fuscus 359410
Mucor lanceolatus 734
Helicostylum pulchrum 605219
Thamnidium elegans 158506
Pilaira anomala 373112
Mucor endophyticus 11425
Blakeslea trispora 444227
Choanephora cucurbitarum A0J61_05496
Gilbertella persicaria 483042
Cokeromyces recurvatus 507455
Mycotypha africana 836211
Rhizopus azygosporus CU097_006871
Rhizopus azygosporus CU097_006867
Rhizopus delemar RO3G_04312
Rhizopus microsporus 238668
Sporodiniella umbellata 531312
Pilobolus umbonatus 755573
Absidia glauca SAL97768
Absidia padenii 328386
Absidia repens 328945
Cunninghamella echinulata 159088
Gongronella butleri 371076
Hesseltinella vesiculosa 1331682
Phycomyces nitens 266251
Circinella umbellata 434046
Phascolomyces articulosus 565952
Zychaea mexicana 755405
Lichtheimia corymbifera 7892
Lichtheimia ramosa 94
Dichotomocladium elegans 1951951
Syncephalastrum racemosum 491322
Umbelopsis ramanniana 219798
Umbelopsis sp. 118068
Jimgerdemannia flammicorona 481431
Jimgerdemannia flammicorona 473419
Endogone sp. 91788
Lobosporangium transversale 359779
Lobosporangium transversale 346463
Mortierella multidivaricata 456841
Mortierella multidivaricata 121315
Mortierella elongata 344261
Mortierella elongata 122062
Mortierella humilis 87098
Mortierella humilis 111131
Gigaspora margarita 13772
Gigaspora margarita 11559
Gigaspora rosea 2166708
Gigaspora rosea 2137535
Rhizophagus diaphanus 730051
Rhizophagus diaphanus 685995
Rhizophagus diaphanus 622579
Rhizophagus irregularis 1762417
Rhizophagus irregularis 1672621
Rhizophagus irregularis 1587725
Rhizophagus irregularis 1495185
Rhizophagus clarus 8731
Rhizophagus clarus 1231
Geosiphon pyriformis 20003
Conidiobolus coronatus 7579
Syncephalis fuscata 445388
Olpidium bornovanus 6521
Arb1
0
500
Amino acid number

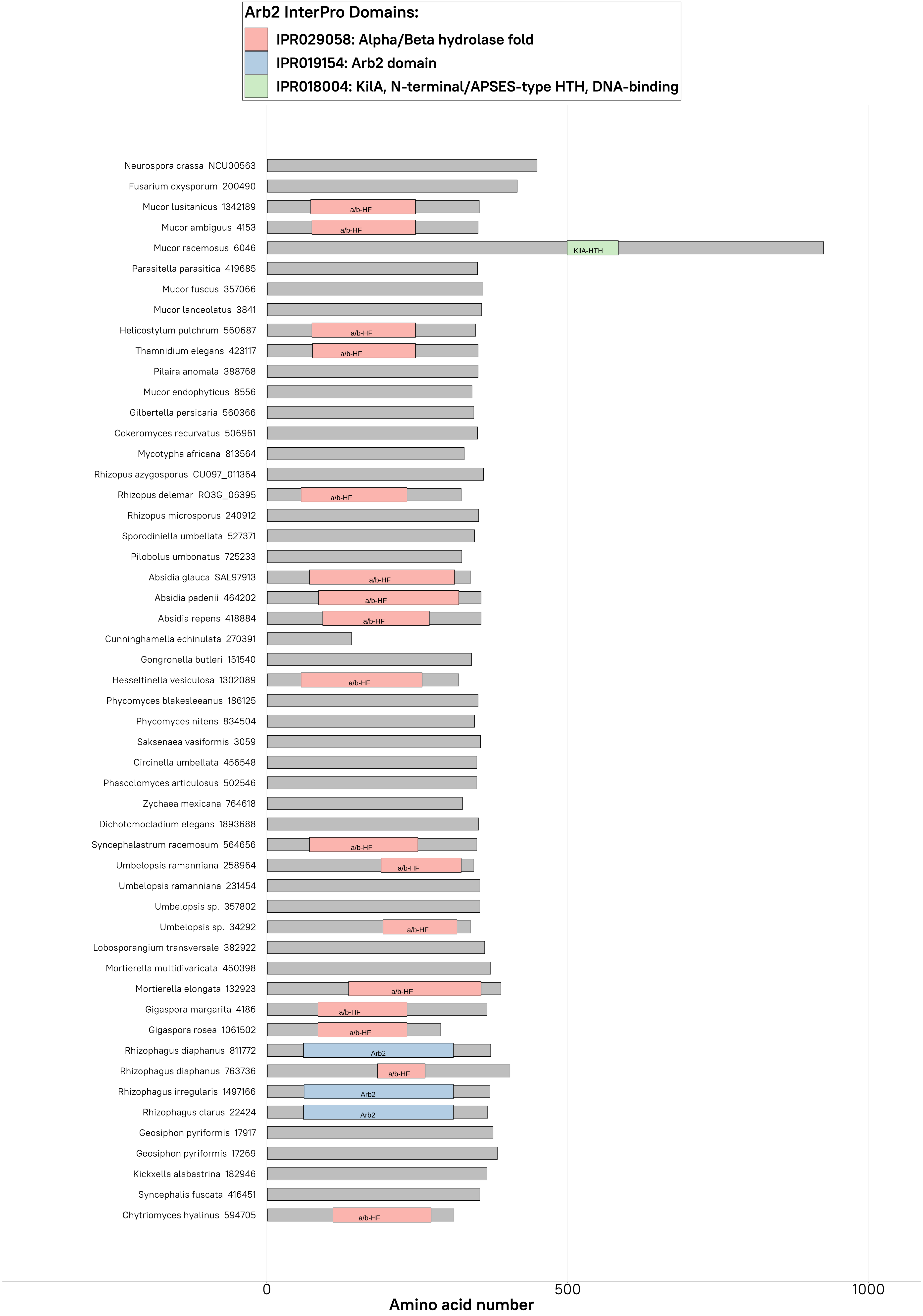
Arb2 InterPro Domains:
IPR029058: Alpha/Beta hydrolase fold
IPR019154: Arb2 domain
IPR018004: KilA, N-terminal/APSES-type HTH, DNA-binding
Neurospora crassa NCU00563
Fusarium oxysporum 200490
Mucor lusitanicus 1342189
Mucor ambiguus 4153
Mucor racemosus 6046
Parasitella parasitica 419685
Mucor fuscus 357066
Mucor lanceolatus 3841
Helicostylum pulchrum 560687
Thamnidium elegans 423117
Pilaira anomala 388768
Mucor endophyticus 8556
Gilbertella persicaria 560366
Cokeromyces recurvatus 506961
Mycotypha africana 813564
Rhizopus azygosporus CU097_011364
Rhizopus delemar RO3G_06395
Rhizopus microsporus 240912
Sporodiniella umbellata 527371
Pilobolus umbonatus 725233
Absidia glauca SAL97913
Absidia padenii 464202
Absidia repens 418884
Cunninghamella echinulata 270391
Gongronella butleri 151540
Hesseltinella vesiculosa 1302089
Phycomyces blakesleeanus 186125
Phycomyces nitens 834504
Saksenaea vasiformis 3059
Circinella umbellata 456548
Phascolomyces articulosus 502546
Zychaea mexicana 764618
Dichotomocladium elegans 1893688
Syncephalastrum racemosum 564656
Umbelopsis ramanniana 258964
Umbelopsis ramanniana 231454
Umbelopsis sp. 357802
Umbelopsis sp. 34292
Lobosporangium transversale 382922
Mortierella multidivaricata 460398
Mortierella elongata 132923
Gigaspora margarita 4186
Gigaspora rosea 1061502
Rhizophagus diaphanus 811772
Rhizophagus diaphanus 763736
Rhizophagus irregularis 1497166
Rhizophagus clarus 22424
Geosiphon pyriformis 17917
Geosiphon pyriformis 17269
Kickxella alabastrina 182946
Syncephalis fuscata 416451
Chytriomyces hyalinus 594705
a/b-HF
KilA-HTH
Arb2
0
500
1000
Amino acid number

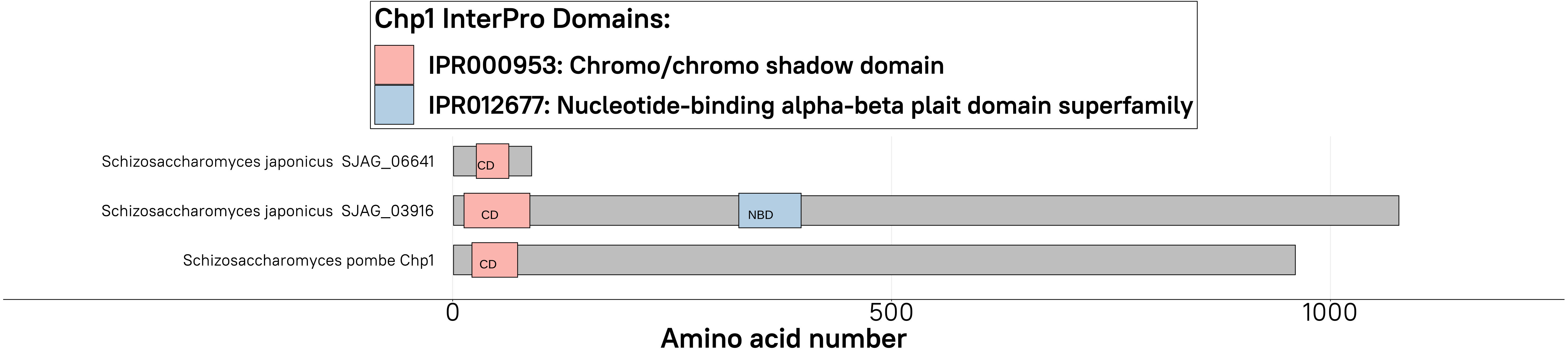

Chp1 InterPro Domains:
IPR000953: Chromo/chromo shadow domain
IPR012677: Nucleotide-binding alpha-beta plait domain superfamily
Schizosaccharomyces japonicus SJAG_06641
Schizosaccharomyces japonicus SJAG_03916
Schizosaccharomyces pombe Chp1
CD
CD
NBD
CD
0
500
1000
Amino acid number

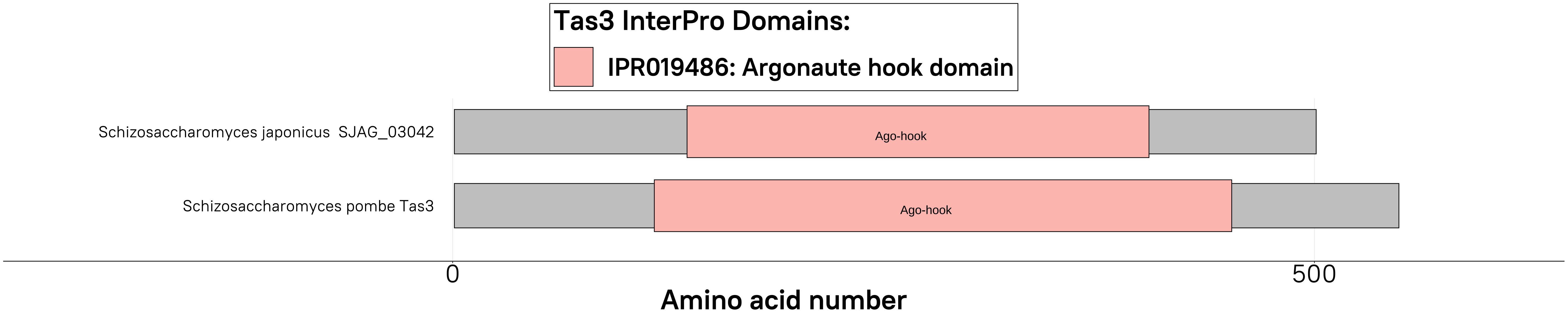
Tas3 InterPro Domains:
IPR019486: Argonaute hook domain
Schizosaccharomyces japonicus SJAG_03042
Ago-hook
Schizosaccharomyces pombe Tas3
Ago-hook
0
500
Amino acid number

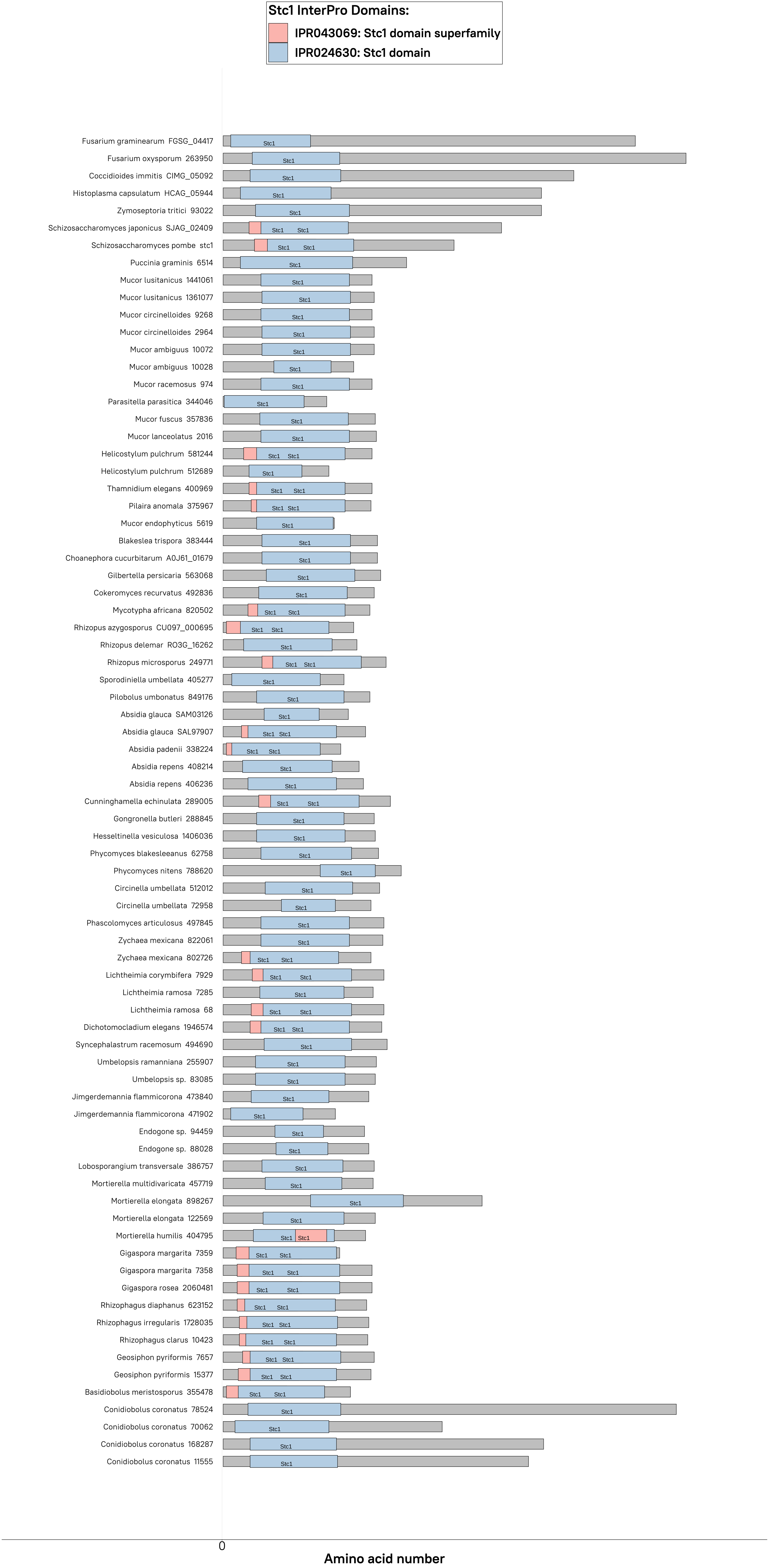
Stc1 InterPro Domains:
IPR043069: Stc1 domain superfamily
IPR024630: Stc1 domain
Fusarium graminearum FGSG_04417
Fusarium oxysporum 263950
Coccidioides immitis CIMG_05092
Histoplasma capsulatum HCAG_05944
Zymoseptoria tritici 93022
Schizosaccharomyces japonicus SJAG_02409
Schizosaccharomyces pombe stc1
Puccinia graminis 6514
Mucor lusitanicus 1441061
Mucor lusitanicus 1361077
Mucor circinelloides 9268
Mucor circinelloides 2964
Mucor ambiguus 10072
Mucor ambiguus 10028
Mucor racemosus 974
Parasitella parasitica 344046
Mucor fuscus 357836
Mucor lanceolatus 2016
Helicostylum pulchrum 581244
Helicostylum pulchrum 512689
Thamnidium elegans 400969
Pilaira anomala 375967
Mucor endophyticus 5619
Blakeslea trispora 383444
Choanephora cucurbitarum A0J61_01679
Gilbertella persicaria 563068
Cokeromyces recurvatus 492836
Mycotypha africana 820502
Rhizopus azygosporus CU097_000695
Rhizopus delemar RO3G_16262
Rhizopus microsporus 249771
Sporodiniella umbellata 405277
Pilobolus umbonatus 849176
Absidia glauca SAM03126
Absidia glauca SAL97907
Absidia padenii 338224
Absidia repens 408214
Absidia repens 406236
Cunninghamella echinulata 289005
Gongronella butleri 288845
Hesseltinella vesiculosa 1406036
Phycomyces blakesleeanus 62758
Phycomyces nitens 788620
Circinella umbellata 512012
Circinella umbellata 72958
Phascolomyces articulosus 497845
Zychaea mexicana 822061
Zychaea mexicana 802726
Lichtheimia corymbifera 7929
Lichtheimia ramosa 7285
Lichtheimia ramosa 68
Dichotomocladium elegans 1946574
Syncephalastrum racemosum 494690
Umbelopsis ramanniana 255907
Umbelopsis sp. 83085
Jimgerdemannia flammicorona 473840
Jimgerdemannia flammicorona 471902
Endogone sp. 94459
Endogone sp. 88028
Lobosporangium transversale 386757
Mortierella multidivaricata 457719
Mortierella elongata 898267
Mortierella elongata 122569
Mortierella humilis 404795
Gigaspora margarita 7359
Gigaspora margarita 7358
Gigaspora rosea 2060481
Rhizophagus diaphanus 623152
Rhizophagus irregularis 1728035
Rhizophagus clarus 10423
Geosiphon pyriformis 7657
Geosiphon pyriformis 15377
Basidiobolus meristosporus 355478
Conidiobolus coronatus 78524
Conidiobolus coronatus 70062
Conidiobolus coronatus 168287
Conidiobolus coronatus 11555
0
Amino acid number

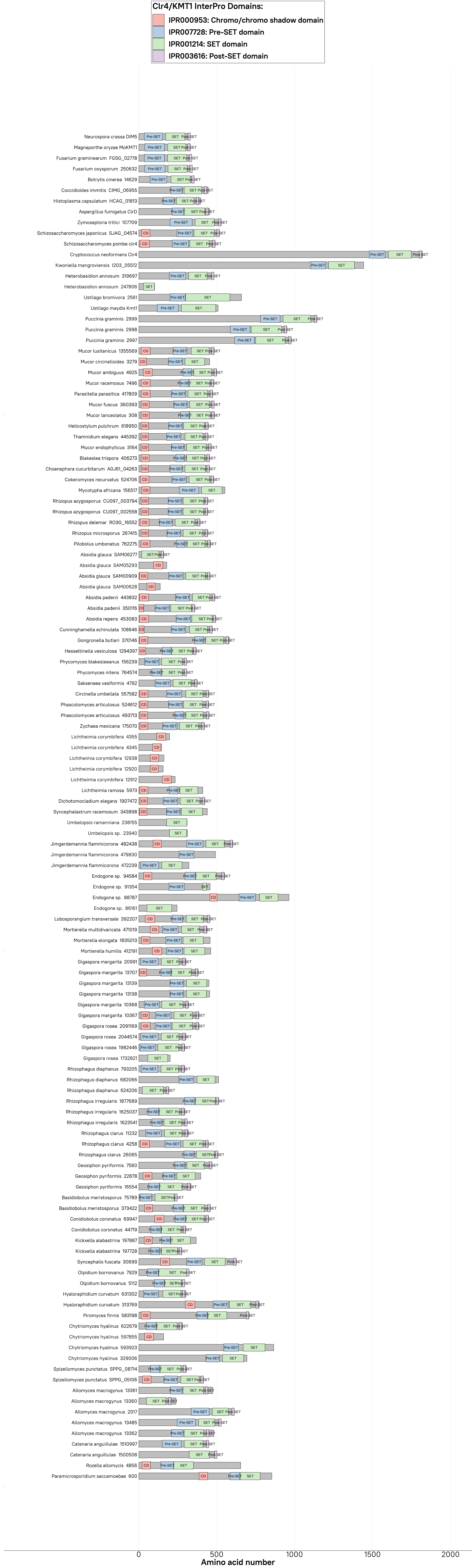

Clr4/KMT1 InterPro Domains:
IPR000953: Chromo/chromo shadow domain
IPR007728: Pre-SET domain
IPR001214: SET domain
IPR003616: Post-SET domain
Neurospora crassa DIM5
Magnaporthe oryzae MoKMT1
Fusarium graminearum FGSG_02778
Fusarium oxysporum 250632
Botrytis cinerea 14629
Coccidioides immitis CIMG_06955
Histoplasma capsulatum HCAG_01813
Aspergillus fumigatus ClrD
Zymoseptoria tritici 107709
Schizosaccharomyces japonicus SJAG_04574
Schizosaccharomyces pombe clr4
Cryptococcus neoformans Clr4
Kwoniella mangroviensis I203_05512
Heterobasidion annosum 319697
Heterobasidion annosum 247806
Ustilago bromivora 2561
Ustilago maydis Kmt1
Puccinia graminis 2999
Puccinia graminis 2998
Puccinia graminis 2997
Mucor lusitanicus 1355569
Mucor circinelloides 3279
Mucor ambiguus 4925
Mucor racemosus 7496
Parasitella parasitica 417809
Mucor fuscus 360393
Mucor lanceolatus 308
Helicostylum pulchrum 618950
Thamnidium elegans 445392
Mucor endophyticus 3164
Blakeslea trispora 406273
Choanephora cucurbitarum A0J61_04263
Cokeromyces recurvatus 524706
Mycotypha africana 156517
Rhizopus azygosporus CU097_003794
Rhizopus azygosporus CU097_002558
Rhizopus delemar RO3G_16552
Rhizopus microsporus 267415
Pilobolus umbonatus 762275
Absidia glauca SAM06277
Absidia glauca SAM05293
Absidia glauca SAM00909
Absidia glauca SAM00628
Absidia padenii 443832
Absidia padenii 350116
Absidia repens 453083
Cunninghamella echinulata 108646
Gongronella butleri 370146
Hesseltinella vesiculosa 1294397
Phycomyces blakesleeanus 156239
Phycomyces nitens 764574
Saksenaea vasiformis 4792
Circinella umbellata 557582
Phascolomyces articulosus 524612
Phascolomyces articulosus 493713
Zychaea mexicana 175070
Lichtheimia corymbifera 4355
Lichtheimia corymbifera 4345
Lichtheimia corymbifera 12938
Lichtheimia corymbifera 12920
Lichtheimia corymbifera 12912
Lichtheimia ramosa 5973
Dichotomocladium elegans 1907472
Syncephalastrum racemosum 343898
Umbelopsis ramanniana 238155
Umbelopsis sp. 23940
Jimgerdemannia flammicorona 482438
Jimgerdemannia flammicorona 479830
Jimgerdemannia flammicorona 472239
Endogone sp. 94584
Endogone sp. 91354
Endogone sp. 88787
Endogone sp. 86161
Lobosporangium transversale 392207
Mortierella multidivaricata 471019
Mortierella elongata 1835013
Mortierella humilis 412191
Gigaspora margarita 20991
Gigaspora margarita 13707
Gigaspora margarita 13139
Gigaspora margarita 13138
Gigaspora margarita 10368
Gigaspora margarita 10367
Gigaspora rosea 2091169
Gigaspora rosea 2044574
Gigaspora rosea 1982446
Gigaspora rosea 1732821
Rhizophagus diaphanus 793205
Rhizophagus diaphanus 682066
Rhizophagus diaphanus 624206
Rhizophagus irregularis 1877689
Rhizophagus irregularis 1625037
Rhizophagus irregularis 1623541
Rhizophagus clarus 11232
Rhizophagus clarus 4258
Rhizophagus clarus 26065
Geosiphon pyriformis 7560
Geosiphon pyriformis 22878
Geosiphon pyriformis 16554
Basidiobolus meristosporus 75789
Basidiobolus meristosporus 373422
Conidiobolus coronatus 69947
Conidiobolus coronatus 44719
Kickxella alabastrina 197887
Kickxella alabastrina 197728
Syncephalis fuscata 30699
Olpidium bornovanus 7929
Olpidium bornovanus 5112
Hyaloraphidium curvatum 631302
Hyaloraphidium curvatum 313769
Piromyces finnis 583198
Chytriomyces hyalinus 622679
Chytriomyces hyalinus 597855
Chytriomyces hyalinus 593923
Chytriomyces hyalinus 329006
Spizellomyces punctatus SPPG_08714
Spizellomyces punctatus SPPG_05106
Allomyces macrogynus 13361
Allomyces macrogynus 13360
Allomyces macrogynus 2017
Allomyces macrogynus 13485
Allomyces macrogynus 13362
Catenaria anguillulae 1510997
Catenaria anguillulae 1500508
Rozella allomycis 4856
Paramicrosporidium saccamoebae 600
0
500
1000
1500
2000
Amino acid number

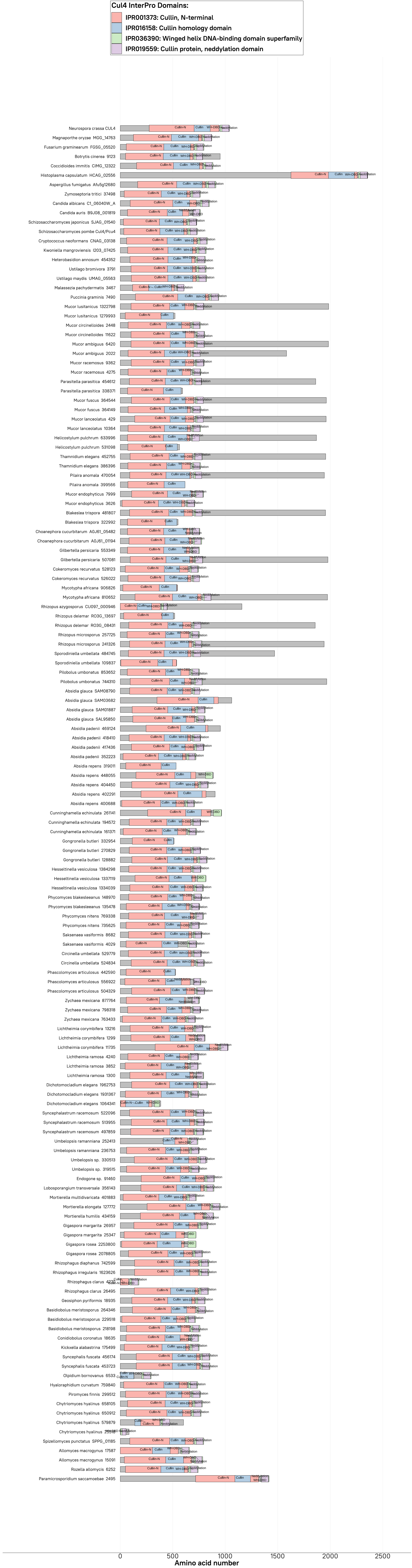
Cul4 InterPro Domains:
IPR001373: Cullin, N-terminal
IPR016158: Cullin homology domain
IPR036390: Winged helix DNA-binding domain superfamily
IPR019559: Cullin protein, neddylation domain
Neurospora crassa CUL4
Magnaporthe oryzae MGG_14763
Fusarium graminearum FGSG_05520
Botrytis cinerea 9123
Coccidioides immitis CIMG_12322
Histoplasma capsulatum HCAG_02556
Aspergillus fumigatus Afu5g12680
Zymoseptoria tritici 37498
Candida albicans C1_06040W_A
Candida auris B9J08_001819
Schizosaccharomyces japonicus SJAG_01540
Schizosaccharomyces pombe Cul4/Pcu4
Cryptococcus neoformans CNAG_03138
Kwoniella mangroviensis I203_07425
Heterobasidion annosum 454352
Ustilago bromivora 3791
Ustilago maydis UMAG_05563
Malassezia pachydermatis 3467
Puccinia graminis 7490
Mucor lusitanicus 1322798
Mucor lusitanicus 1279993
Mucor circinelloides 2448
Mucor circinelloides 11622
Mucor ambiguus 6420
Mucor ambiguus 2022
Mucor racemosus 9362
Mucor racemosus 4275
Parasitella parasitica 454612
Parasitella parasitica 338371
Mucor fuscus 364544
Mucor fuscus 364149
Mucor lanceolatus 429
Mucor lanceolatus 10364
Helicostylum pulchrum 633996
Helicostylum pulchrum 531098
Thamnidium elegans 452755
Thamnidium elegans 386396
Pilaira anomala 470054
Pilaira anomala 399566
Mucor endophyticus 7999
Mucor endophyticus 3626
Blakeslea trispora 481807
Blakeslea trispora 322992
Choanephora cucurbitarum A0J61_05482
Choanephora cucurbitarum A0J61_01194
Gilbertella persicaria 553349
Gilbertella persicaria 507081
Cokeromyces recurvatus 528123
Cokeromyces recurvatus 526022
Mycotypha africana 906826
Mycotypha africana 810652
Rhizopus azygosporus CU097_000946
Rhizopus delemar RO3G_13697
Rhizopus delemar RO3G_08431
Rhizopus microsporus 257725
Rhizopus microsporus 241326
Sporodiniella umbellata 484745
Sporodiniella umbellata 109837
Pilobolus umbonatus 853652
Pilobolus umbonatus 744310
Absidia glauca SAM08790
Absidia glauca SAM03682
Absidia glauca SAM01887
Absidia glauca SAL95850
Absidia padenii 469124
Absidia padenii 418410
Absidia padenii 417436
Absidia padenii 352223
Absidia repens 319011
Absidia repens 448055
Absidia repens 404450
Absidia repens 402291
Absidia repens 400688
Cunninghamella echinulata 261141
Cunninghamella echinulata 194572
Cunninghamella echinulata 161371
Gongronella butleri 332954
Gongronella butleri 270829
Gongronella butleri 128882
Hesseltinella vesiculosa 1384298
Hesseltinella vesiculosa 1337119
Hesseltinella vesiculosa 1334039
Phycomyces blakesleeanus 148970
Phycomyces blakesleeanus 135478
Phycomyces nitens 769338
Phycomyces nitens 735625
Saksenaea vasiformis 8682
Saksenaea vasiformis 4029
Circinella umbellata 529779
Circinella umbellata 524834
Phascolomyces articulosus 442590
Phascolomyces articulosus 556922
Phascolomyces articulosus 504329
Zychaea mexicana 877764
Zychaea mexicana 798318
Zychaea mexicana 763433
Lichtheimia corymbifera 13216
Lichtheimia corymbifera 1299
Lichtheimia corymbifera 11735
Lichtheimia ramosa 4240
Lichtheimia ramosa 3852
Lichtheimia ramosa 1300
Dichotomocladium elegans 1962753
Dichotomocladium elegans 1931367
Dichotomocladium elegans 1064341
Syncephalastrum racemosum 522096
Syncephalastrum racemosum 513955
Syncephalastrum racemosum 497859
Umbelopsis ramanniana 252413
Umbelopsis ramanniana 236753
Umbelopsis sp. 330513
Umbelopsis sp. 319515
Endogone sp. 91460
Lobosporangium transversale 356143
Mortierella multidivaricata 401883
Mortierella elongata 127772
Mortierella humilis 434159
Gigaspora margarita 26957
Gigaspora margarita 25347
Gigaspora rosea 2253800
Gigaspora rosea 2078805
Rhizophagus diaphanus 742599
Rhizophagus irregularis 1623626
Rhizophagus clarus 4277
Rhizophagus clarus 26495
Geosiphon pyriformis 18935
Basidiobolus meristosporus 264346
Basidiobolus meristosporus 229518
Basidiobolus meristosporus 218198
Conidiobolus coronatus 18635
Kickxella alabastrina 175499
Syncephalis fuscata 456174
Syncephalis fuscata 453723
Olpidium bornovanus 6533
Hyaloraphidium curvatum 759840
Piromyces finnis 299512
Chytriomyces hyalinus 658105
Chytriomyces hyalinus 650912
Chytriomyces hyalinus 579879
Chytriomyces hyalinus 25518
Spizellomyces punctatus SPPG_01185
Allomyces macrogynus 17587
Allomyces macrogynus 15091
Rozella allomycis 6252
Paramicrosporidium saccamoebae 2495
0
500
1000
1500
2000
2500
Amino acid number

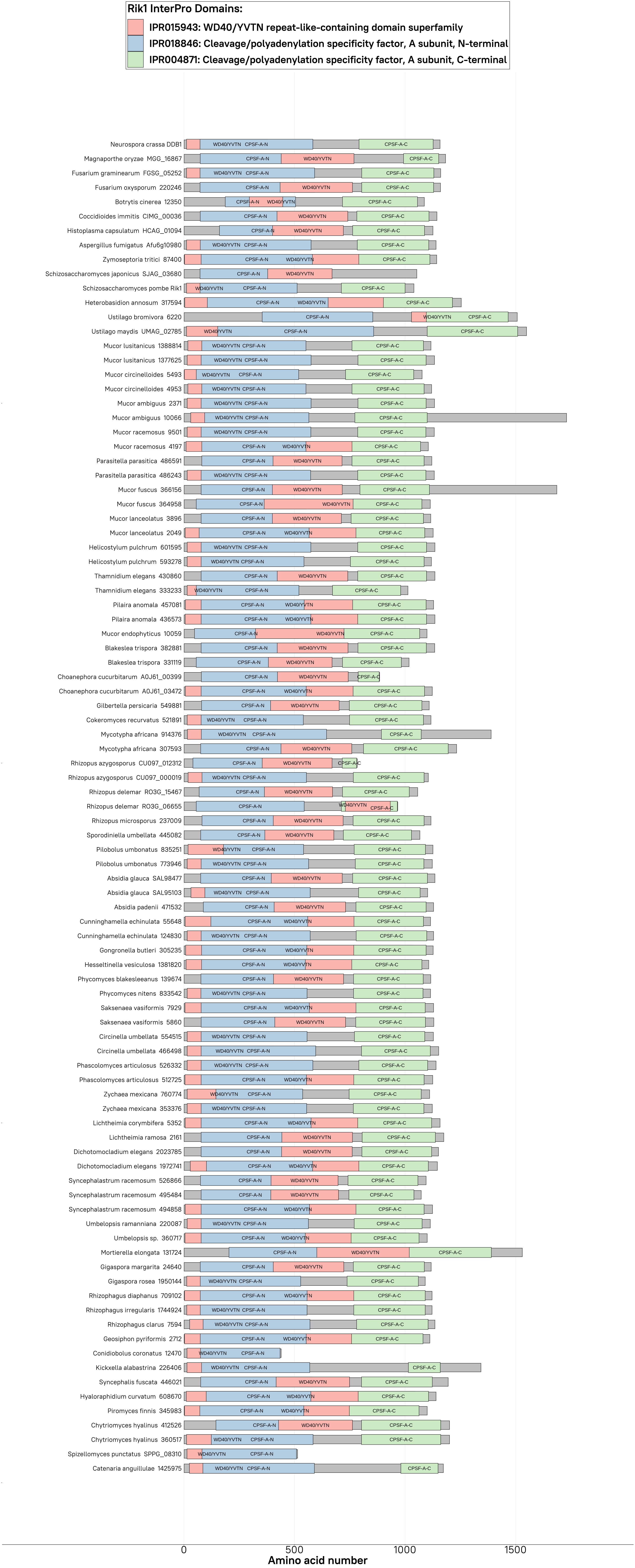
Rik1 InterPro Domains:
IPR015943: WD40/YVTN repeat-like-containing domain superfamily
IPR018846: Cleavage/polyadenylation specificity factor, A subunit, N-terminal
IPR004871: Cleavage/polyadenylation specificity factor, A subunit, C-terminal
Neurospora crassa DDB1
Magnaporthe oryzae MGG_16867
Fusarium graminearum FGSG_05252
Fusarium oxysporum 220246
Botrytis cinerea 12350
Coccidioides immitis CIMG_00036
Histoplasma capsulatum HCAG_01094
Aspergillus fumigatus Afu6g10980
Zymoseptoria tritici 87400
Schizosaccharomyces japonicus SJAG_03680
Schizosaccharomyces pombe Rik1
Heterobasidion annosum 317594
Ustilago bromivora 6220
Ustilago maydis UMAG_02785
Mucor lusitanicus 1388814
Mucor lusitanicus 1377625
Mucor circinelloides 5493
Mucor circinelloides 4953
Mucor ambiguus 2371
Mucor ambiguus 10066
Mucor racemosus 9501
Mucor racemosus 4197
Parasitella parasitica 486591
Parasitella parasitica 486243
Mucor fuscus 366156
Mucor fuscus 364958
Mucor lanceolatus 3896
Mucor lanceolatus 2049
Helicostylum pulchrum 601595
Helicostylum pulchrum 593278
Thamnidium elegans 430860
Thamnidium elegans 333233
Pilaira anomala 457081
Pilaira anomala 436573
Mucor endophyticus 10059
Blakeslea trispora 382881
Blakeslea trispora 331119
Choanephora cucurbitarum A0J61_00399
Choanephora cucurbitarum A0J61_03472
Gilbertella persicaria 549881
Cokeromyces recurvatus 521891
Mycotypha africana 914376
Mycotypha africana 307593
Rhizopus azygosporus CU097_012312
Rhizopus azygosporus CU097_000019
Rhizopus delemar RO3G_15467
Rhizopus delemar RO3G_06655
Rhizopus microsporus 237009
Sporodiniella umbellata 445082
Pilobolus umbonatus 835251
Pilobolus umbonatus 773946
Absidia glauca SAL98477
Absidia glauca SAL95103
Absidia padenii 471532
Cunninghamella echinulata 55648
Cunninghamella echinulata 124830
Gongronella butleri 305235
Hesseltinella vesiculosa 1381820
Phycomyces blakesleeanus 139674
Phycomyces nitens 833542
Saksenaea vasiformis 7929
Saksenaea vasiformis 5860
Circinella umbellata 554515
Circinella umbellata 466498
Phascolomyces articulosus 526332
Phascolomyces articulosus 512725
Zychaea mexicana 760774
Zychaea mexicana 353376
Lichtheimia corymbifera 5352
Lichtheimia ramosa 2161
Dichotomocladium elegans 2023785
Dichotomocladium elegans 1972741
Syncephalastrum racemosum 526866
Syncephalastrum racemosum 495484
Syncephalastrum racemosum 494858
Umbelopsis ramanniana 220087
Umbelopsis sp. 360717
Mortierella elongata 131724
Gigaspora margarita 24640
Gigaspora rosea 1950144
Rhizophagus diaphanus 709102
Rhizophagus irregularis 1744924
Rhizophagus clarus 7594
Geosiphon pyriformis 2712
Conidiobolus coronatus 12470
Kickxella alabastrina 226406
Syncephalis fuscata 446021
Hyaloraphidium curvatum 608670
Piromyces finnis 345983
Chytriomyces hyalinus 412526
Chytriomyces hyalinus 360517
Spizellomyces punctatus SPPG_08310
Catenaria anguillulae 1425975
WD40/YVTN
CPSF-A-N
CPSF-A-C
0
500
1000
1500
Amino acid number

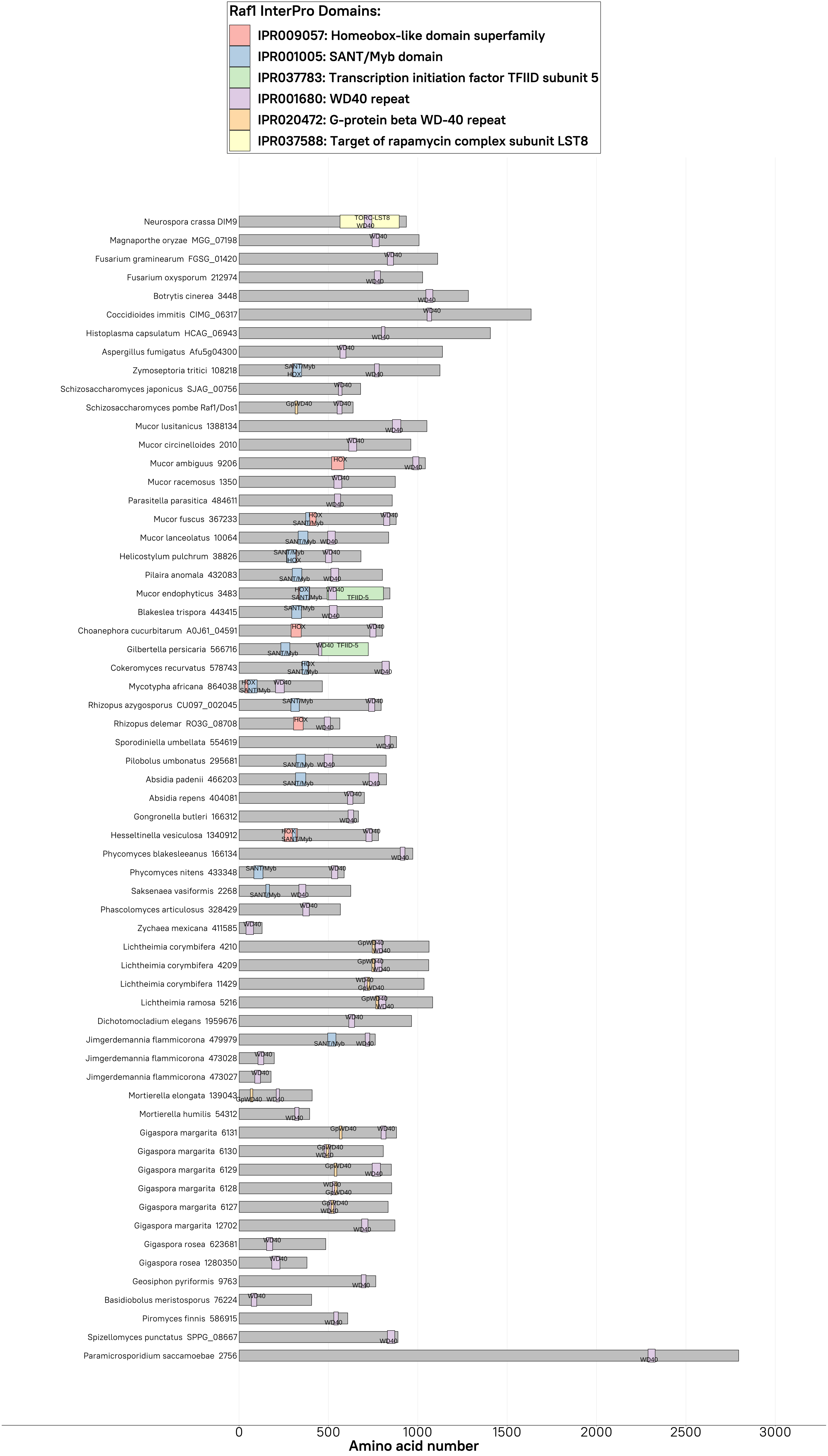

Raf1 InterPro Domains:
IPR009057: Homeobox-like domain superfamily
IPR001005: SANT/Myb domain
IPR037783: Transcription initiation factor TFIID subunit 5
IPR001680: WD40 repeat
IPR020472: G-protein beta WD-40 repeat
IPR037588: Target of rapamycin complex subunit LST8
Neurospora crassa DIM9
Magnaporthe oryzae MGG_07198
Fusarium graminearum FGSG_01420
Fusarium oxysporum 212974
Botrytis cinerea 3448
Coccidioides immitis CIMG_06317
Histoplasma capsulatum HCAG_06943
Aspergillus fumigatus Afu5g04300
Zymoseptoria tritici 108218
Schizosaccharomyces japonicus SJAG_00756
Schizosaccharomyces pombe Raf1/Dos1
Mucor lusitanicus 1388134
Mucor circinelloides 2010
Mucor ambiguus 9206
Mucor racemosus 1350
Parasitella parasitica 484611
Mucor fuscus 367233
Mucor lanceolatus 10064
Helicostylum pulchrum 38826
Pilaira anomala 432083
Mucor endophyticus 3483
Blakeslea trispora 443415
Choanephora cucurbitarum A0J61_04591
Gilbertella persicaria 566716
Cokeromyces recurvatus 578743
Mycotypha africana 864038
Rhizopus azygosporus CU097_002045
Rhizopus delemar RO3G_08708
Sporodiniella umbellata 554619
Pilobolus umbonatus 295681
Absidia padenii 466203
Absidia repens 404081
Gongronella butleri 166312
Hesseltinella vesiculosa 1340912
Phycomyces blakesleeanus 166134
Phycomyces nitens 433348
Saksenaea vasiformis 2268
Phascolomyces articulosus 328429
Zychaea mexicana 411585
Lichtheimia corymbifera 4210
Lichtheimia corymbifera 4209
Lichtheimia corymbifera 11429
Lichtheimia ramosa 5216
Dichotomocladium elegans 1959676
Jimgerdemannia flammicorona 479979
Jimgerdemannia flammicorona 473028
Jimgerdemannia flammicorona 473027
Mortierella elongata 139043
Mortierella humilis 54312
Gigaspora margarita 6131
Gigaspora margarita 6130
Gigaspora margarita 6129
Gigaspora margarita 6128
Gigaspora margarita 6127
Gigaspora margarita 12702
Gigaspora rosea 623681
Gigaspora rosea 1280350
Geosiphon pyriformis 9763
Basidiobolus meristosporus 76224
Piromyces finnis 586915
Spizellomyces punctatus SPPG_08667
Paramicrosporidium saccamoebae 2756
0
500
1000
1500
2000
2500
3000
Amino acid number

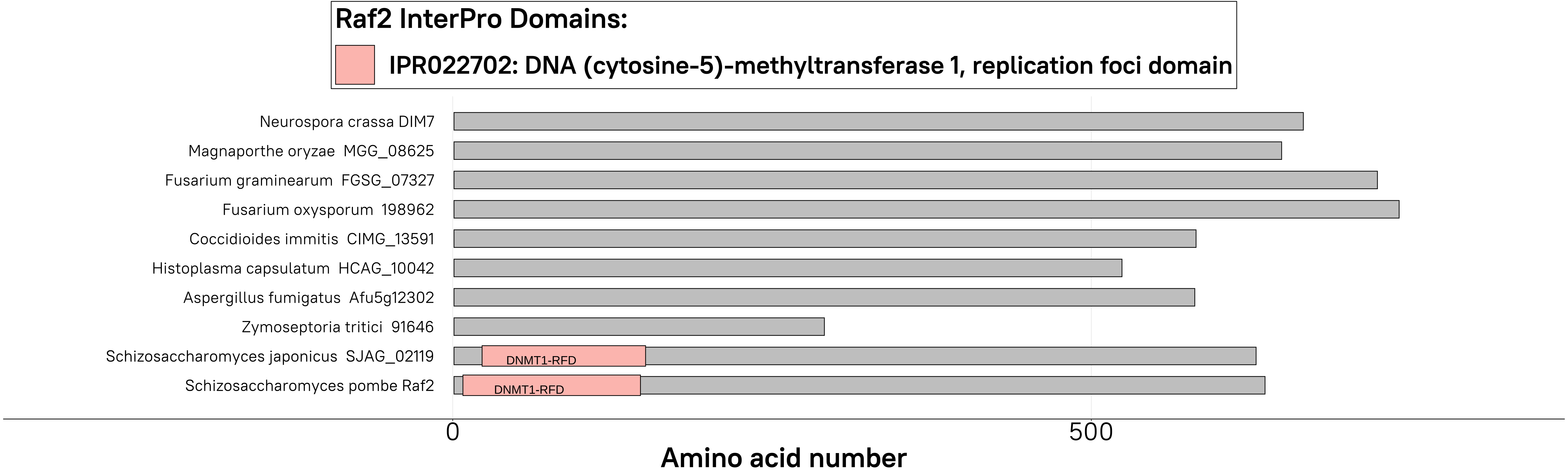

Raf2 InterPro Domains:
IPR022702: DNA (cytosine-5)-methyltransferase 1, replication foci domain
Neurospora crassa DIM7
Magnaporthe oryzae MGG_08625
Fusarium graminearum FGSG_07327
Fusarium oxysporum 198962
Coccidioides immitis CIMG_13591
Histoplasma capsulatum HCAG_10042
Aspergillus fumigatus Afu5g12302
Zymoseptoria tritici 91646
Schizosaccharomyces japonicus SJAG_02119
DNMT1-RFD
Schizosaccharomyces pombe Raf2
DNMT1-RFD
0
500
Amino acid number

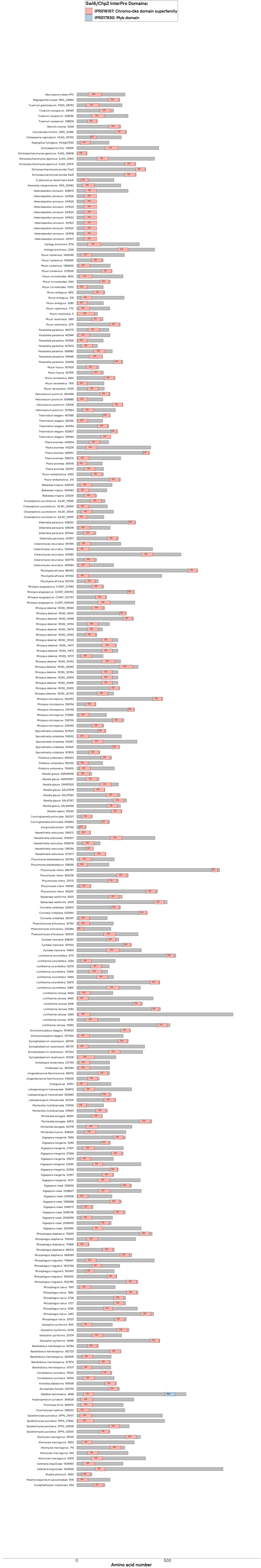
Swi6/Chp2 InterPro Domains:
IPR016197: Chromo-like domain superfamily
IPR017930: Myb domain
0
500
Amino acid number

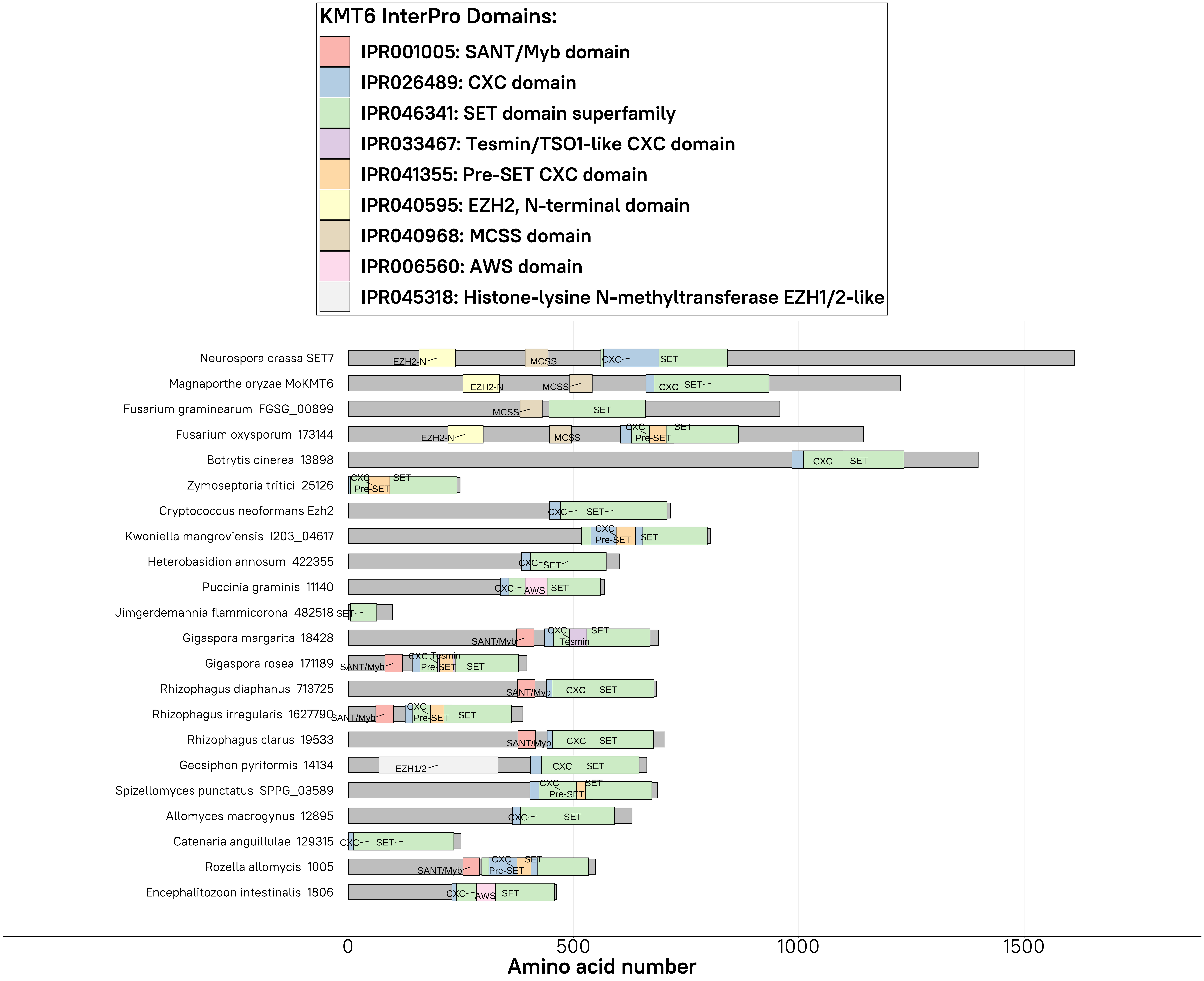

KMT6 InterPro Domains:
IPR001005: SANT/Myb domain
IPR026489: CXC domain
IPR046341: SET domain superfamily
IPR033467: Tesmin/TSO1-like CXC domain
IPR041355: Pre-SET CXC domain
IPR040595: EZH2, N-terminal domain
IPR040968: MCSS domain
IPR006560: AWS domain
IPR045318: Histone-lysine N-methyltransferase EZH1/2-like
Neurospora crassa SET7
Magnaporthe oryzae MoKMT6
Fusarium graminearum FGSG_00899
Fusarium oxysporum 173144
Botrytis cinerea 13898
Zymoseptoria tritici 25126
Cryptococcus neoformans Ezh2
Kwoniella mangroviensis I203_04617
Heterobasidion annosum 422355
Puccinia graminis 11140
Jimgerdemannia flammicorona 482518
Gigaspora margarita 18428
Gigaspora rosea 171189
Rhizophagus diaphanus 713725
Rhizophagus irregularis 1627790
Rhizophagus clarus 19533
Geosiphon pyriformis 14134
Spizellomyces punctatus SPPG_03589
Allomyces macrogynus 12895
Catenaria anguillulae 129315
Rozella allomycis 1005
Encephalitozoon intestinalis 1806
EZH2-N
MCSS
CXC
SET
Pre-SET
SANT/Myb
Tesmin
AWS
EZH1/2
0
500
1000
1500
Amino acid number

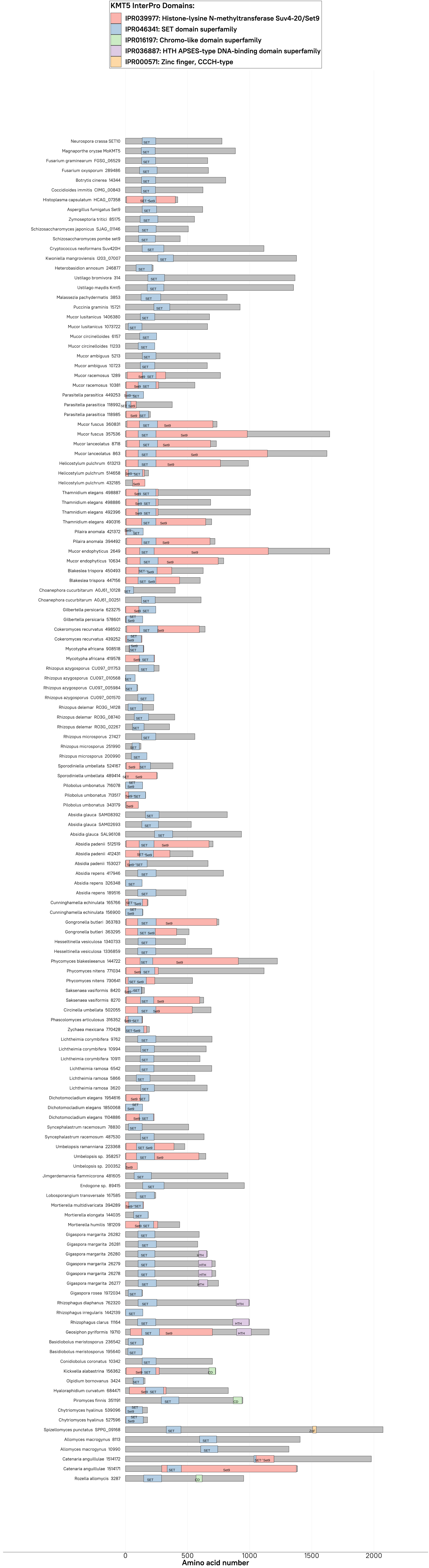

KMT5 InterPro Domains:
IPR039977: Histone-lysine N-methyltransferase Suv4-20/Set9
IPR046341: SET domain superfamily
IPR016197: Chromo-like domain superfamily
IPR036887: HTH APSES-type DNA-binding domain superfamily
IPR000571: Zinc finger, CCCH-type
Neurospora crassa SET10
Magnaporthe oryzae MoKMT5
Fusarium graminearum FGSG_06529
Fusarium oxysporum 289486
Botrytis cinerea 14344
Coccidioides immitis CIMG_00843
Histoplasma capsulatum HCAG_07358
Aspergillus fumigatus Set9
Zymoseptoria tritici 85175
Schizosaccharomyces japonicus SJAG_01146
Schizosaccharomyces pombe set9
Cryptococcus neoformans Suv420H
Kwoniella mangroviensis I203_07007
Heterobasidion annosum 246877
Ustilago bromivora 314
Ustilago maydis Kmt5
Malassezia pachydermatis 3853
Puccinia graminis 15721
Mucor lusitanicus 1406380
Mucor lusitanicus 1073722
Mucor circinelloides 6157
Mucor circinelloides 11233
Mucor ambiguus 5213
Mucor ambiguus 10723
Mucor racemosus 1289
Mucor racemosus 10381
Parasitella parasitica 449253
Parasitella parasitica 118992
Parasitella parasitica 118985
Mucor fuscus 360831
Mucor fuscus 357536
Mucor lanceolatus 8718
Mucor lanceolatus 863
Helicostylum pulchrum 613213
Helicostylum pulchrum 514658
Helicostylum pulchrum 432185
Thamnidium elegans 498887
Thamnidium elegans 498886
Thamnidium elegans 492396
Thamnidium elegans 490316
Pilaira anomala 421372
Pilaira anomala 394492
Mucor endophyticus 2649
Mucor endophyticus 10634
Blakeslea trispora 450493
Blakeslea trispora 447156
Choanephora cucurbitarum A0J61_10128
Choanephora cucurbitarum A0J61_00251
Gilbertella persicaria 623275
Gilbertella persicaria 578601
Cokeromyces recurvatus 498502
Cokeromyces recurvatus 439252
Mycotypha africana 908518
Mycotypha africana 419578
Rhizopus azygosporus CU097_011753
Rhizopus azygosporus CU097_010568
Rhizopus azygosporus CU097_005984
Rhizopus azygosporus CU097_001570
Rhizopus delemar RO3G_14128
Rhizopus delemar RO3G_08740
Rhizopus delemar RO3G_02267
Rhizopus microsporus 27427
Rhizopus microsporus 251990
Rhizopus microsporus 200990
Sporodiniella umbellata 524167
Sporodiniella umbellata 489414
Pilobolus umbonatus 716078
Pilobolus umbonatus 713517
Pilobolus umbonatus 343179
Absidia glauca SAM08392
Absidia glauca SAM02693
Absidia glauca SAL96108
Absidia padenii 512519
Absidia padenii 412431
Absidia padenii 153027
Absidia repens 417946
Absidia repens 326348
Absidia repens 189516
Cunninghamella echinulata 165766
Cunninghamella echinulata 156900
Gongronella butleri 363783
Gongronella butleri 363295
Hesseltinella vesiculosa 1340733
Hesseltinella vesiculosa 1336859
Phycomyces blakesleeanus 144722
Phycomyces nitens 771034
Phycomyces nitens 730641
Saksenaea vasiformis 8420
Saksenaea vasiformis 8270
Circinella umbellata 502055
Phascolomyces articulosus 316352
Zychaea mexicana 770428
Lichtheimia corymbifera 9762
Lichtheimia corymbifera 10994
Lichtheimia corymbifera 10911
Lichtheimia ramosa 6542
Lichtheimia ramosa 5866
Lichtheimia ramosa 3620
Dichotomocladium elegans 1954616
Dichotomocladium elegans 1850068
Dichotomocladium elegans 1104886
Syncephalastrum racemosum 78830
Syncephalastrum racemosum 487530
Umbelopsis ramanniana 223368
Umbelopsis sp. 358257
Umbelopsis sp. 200352
Jimgerdemannia flammicorona 481605
Endogone sp. 89415
Lobosporangium transversale 167585
Mortierella multidivaricata 394289
Mortierella elongata 144035
Mortierella humilis 181209
Gigaspora margarita 26282
Gigaspora margarita 26281
Gigaspora margarita 26280
Gigaspora margarita 26279
Gigaspora margarita 26278
Gigaspora margarita 26277
Gigaspora rosea 1972034
Rhizophagus diaphanus 762320
Rhizophagus irregularis 1442139
Rhizophagus clarus 11164
Geosiphon pyriformis 19710
Basidiobolus meristosporus 236542
Basidiobolus meristosporus 195640
Conidiobolus coronatus 10342
Kickxella alabastrina 156362
Olpidium bornovanus 3424
Hyaloraphidium curvatum 684471
Piromyces finnis 351191
Chytriomyces hyalinus 539096
Chytriomyces hyalinus 527596
Spizellomyces punctatus SPPG_09168
Allomyces macrogynus 8113
Allomyces macrogynus 10990
Catenaria anguillulae 1514172
Catenaria anguillulae 1514171
Rozella allomycis 3287
0
500
1000
1500
2000
Amino acid number

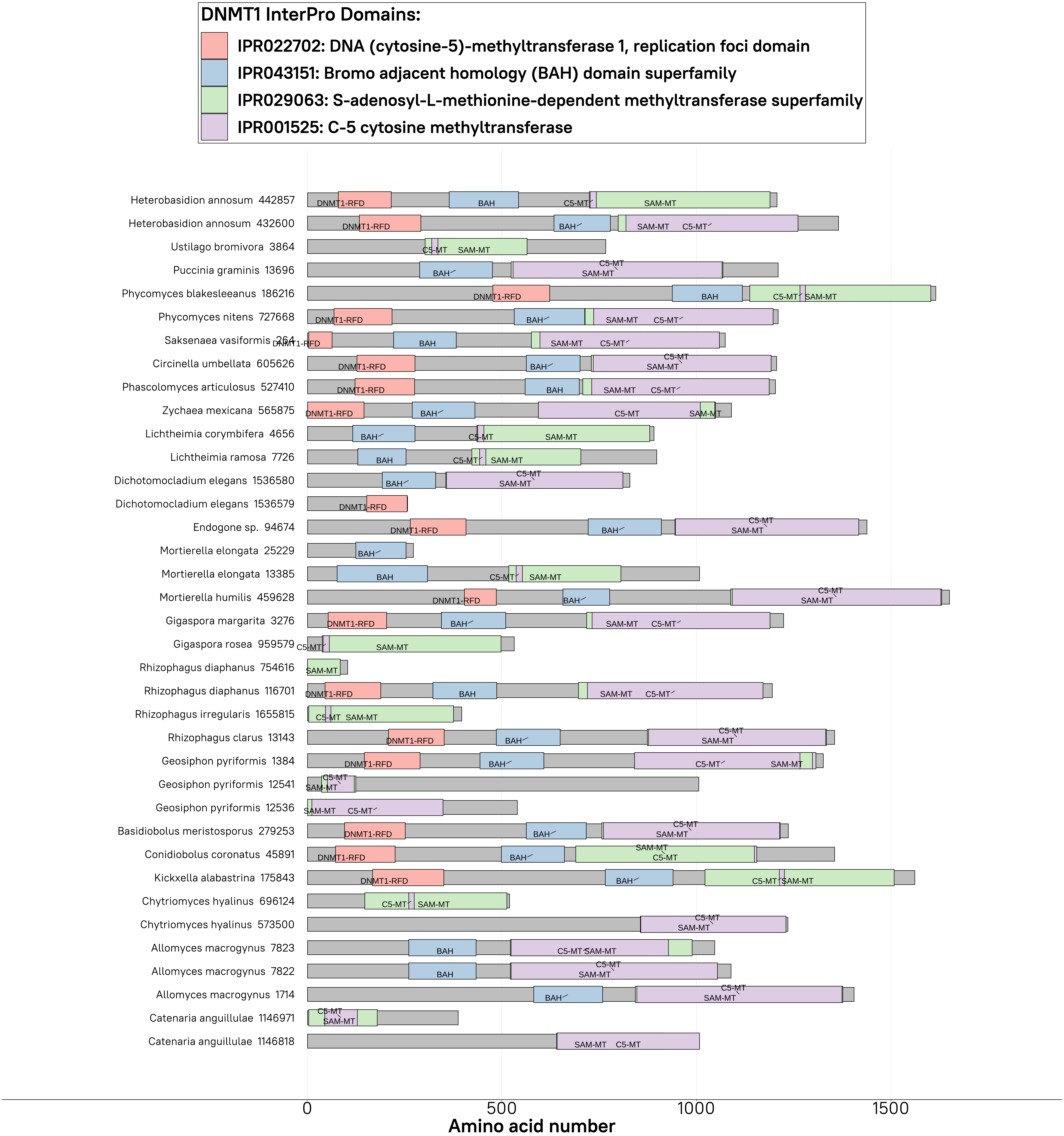
DNMT1 InterPro Domains:
IPR022702: DNA (cytosine-5)-methyltransferase 1, replication foci domain
IPR043151: Bromo adjacent homology (BAH) domain superfamily
IPR029063: S-adenosyl-L-methionine-dependent methyltransferase superfamily
IPR001525: C-5 cytosine methyltransferase
Heterobasidion annosum 442857
Heterobasidion annosum 432600
Ustilago bromivora 3864
Puccinia graminis 13696
Phycomyces blakesleeanus 186216
Phycomyces nitens 727668
Saksenaea vasiformis 264
Circinella umbellata 605626
Phascolomyces articulosus 527410
Zychaea mexicana 565875
Lichtheimia corymbifera 4656
Lichtheimia ramosa 7726
Dichotomocladium elegans 1536580
Dichotomocladium elegans 1536579
Endogone sp. 94674
Mortierella elongata 25229
Mortierella elongata 13385
Mortierella humilis 459628
Gigaspora margarita 3276
Gigaspora rosea 959579
Rhizophagus diaphanus 754616
Rhizophagus diaphanus 116701
Rhizophagus irregularis 1655815
Rhizophagus clarus 13143
Geosiphon pyriformis 1384
Geosiphon pyriformis 12541
Geosiphon pyriformis 12536
Basidiobolus meristosporus 279253
Conidiobolus coronatus 45891
Kickxella alabastrina 175843
Chytriomyces hyalinus 696124
Chytriomyces hyalinus 573500
Allomyces macrogynus 7823
Allomyces macrogynus 7822
Allomyces macrogynus 1714
Catenaria anguillulae 1146971
Catenaria anguillulae 1146818
DNMT1-RFD
BAH
SAM-MT
C5-MT
0
500
1000
1500
Amino acid number

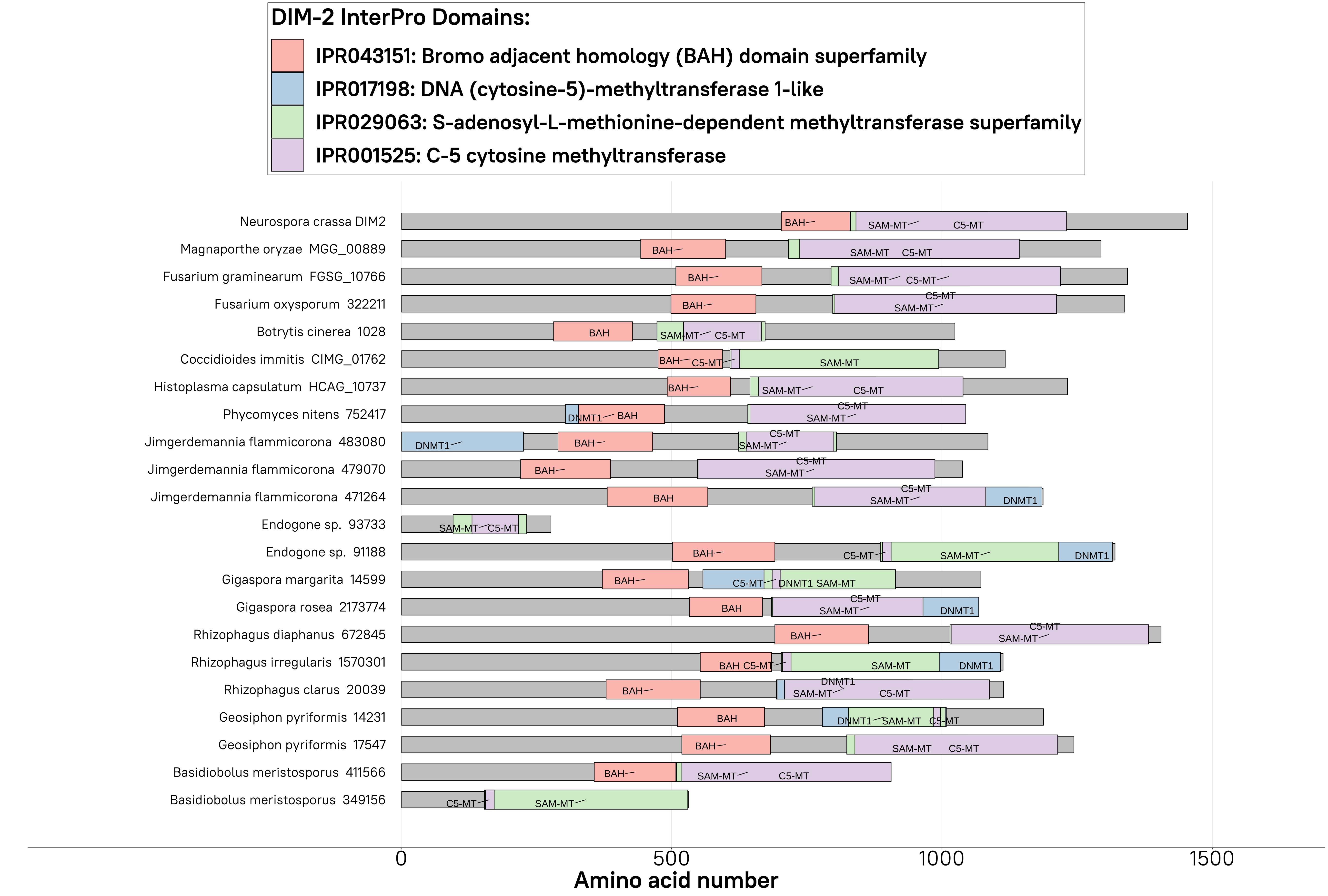
DIM-2 InterPro Domains:
IPR043151: Bromo adjacent homology (BAH) domain superfamily
IPR017198: DNA (cytosine-5)-methyltransferase 1-like
IPR029063: S-adenosyl-L-methionine-dependent methyltransferase superfamily
IPR001525: C-5 cytosine methyltransferase
Neurospora crassa DIM2
Magnaporthe oryzae MGG_00889
Fusarium graminearum FGSG_10766
Fusarium oxysporum 322211
Botrytis cinerea 1028
Coccidioides immitis CIMG_01762
Histoplasma capsulatum HCAG_10737
Phycomyces nitens 752417
Jimgerdemannia flammicorona 483080
Jimgerdemannia flammicorona 479070
Jimgerdemannia flammicorona 471264
Endogone sp. 93733
Endogone sp. 91188
Gigaspora margarita 14599
Gigaspora rosea 2173774
Rhizophagus diaphanus 672845
Rhizophagus irregularis 1570301
Rhizophagus clarus 20039
Geosiphon pyriformis 14231
Geosiphon pyriformis 17547
Basidiobolus meristosporus 411566
Basidiobolus meristosporus 349156
0
500
1000
1500
Amino acid number

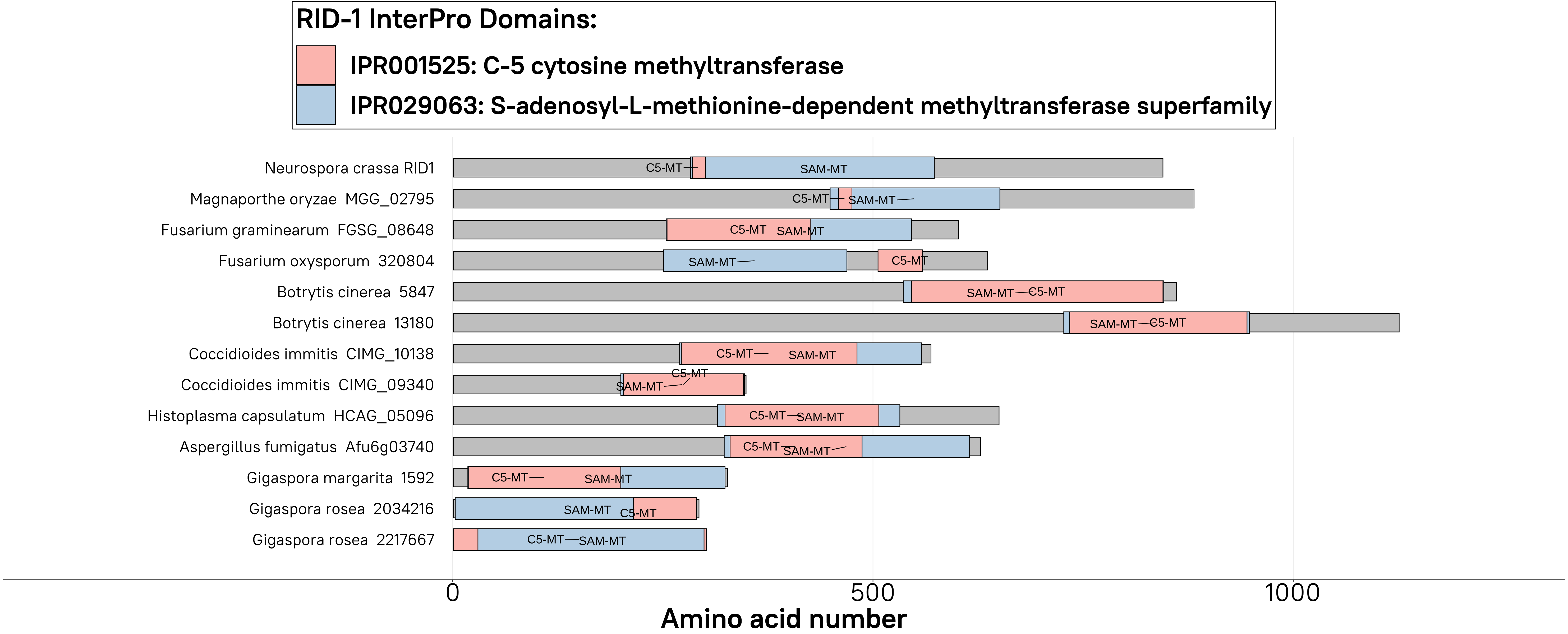

RID-1 InterPro Domains:
IPR001525: C-5 cytosine methyltransferase
IPR029063: S-adenosyl-L-methionine-dependent methyltransferase superfamily
Neurospora crassa RID1
Magnaporthe oryzae MGG_02795
Fusarium graminearum FGSG_08648
Fusarium oxysporum 320804
Botrytis cinerea 5847
Botrytis cinerea 13180
Coccidioides immitis CIMG_10138
Coccidioides immitis CIMG_09340
Histoplasma capsulatum HCAG_05096
Aspergillus fumigatus Afu6g03740
Gigaspora margarita 1592
Gigaspora rosea 2034216
Gigaspora rosea 2217667
C5-MT
SAM-MT
0
500
1000
Amino acid number

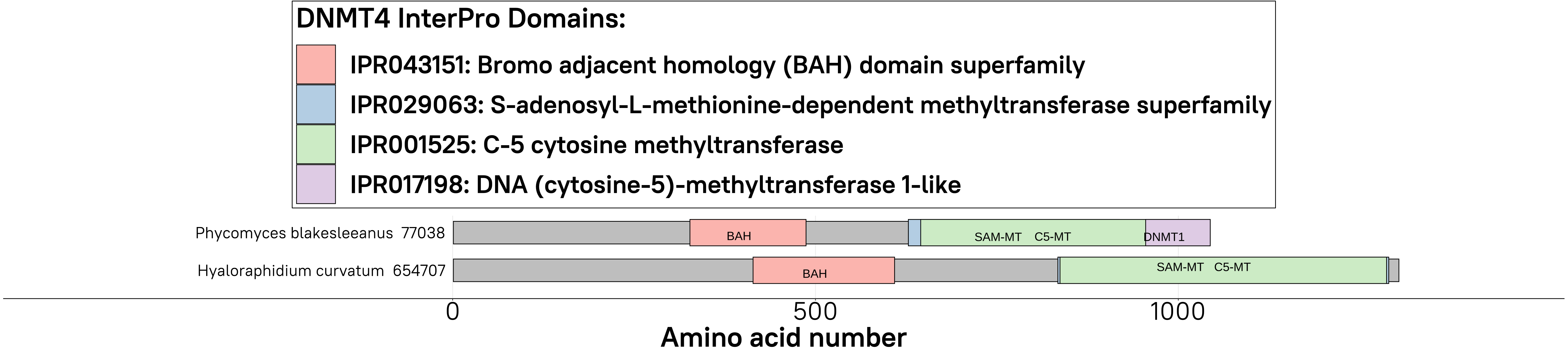
DNMT4 InterPro Domains:
IPR043151: Bromo adjacent homology (BAH) domain superfamily
IPR029063: S-adenosyl-L-methionine-dependent methyltransferase superfamily
IPR001525: C-5 cytosine methyltransferase
IPR017198: DNA (cytosine-5)-methyltransferase 1-like
Phycomyces blakesleeanus 77038
BAH
SAM-MT C5-MT
DNMT1
Hyaloraphidium curvatum 654707
BAH
SAM-MT C5-MT
0
500
1000
Amino acid number

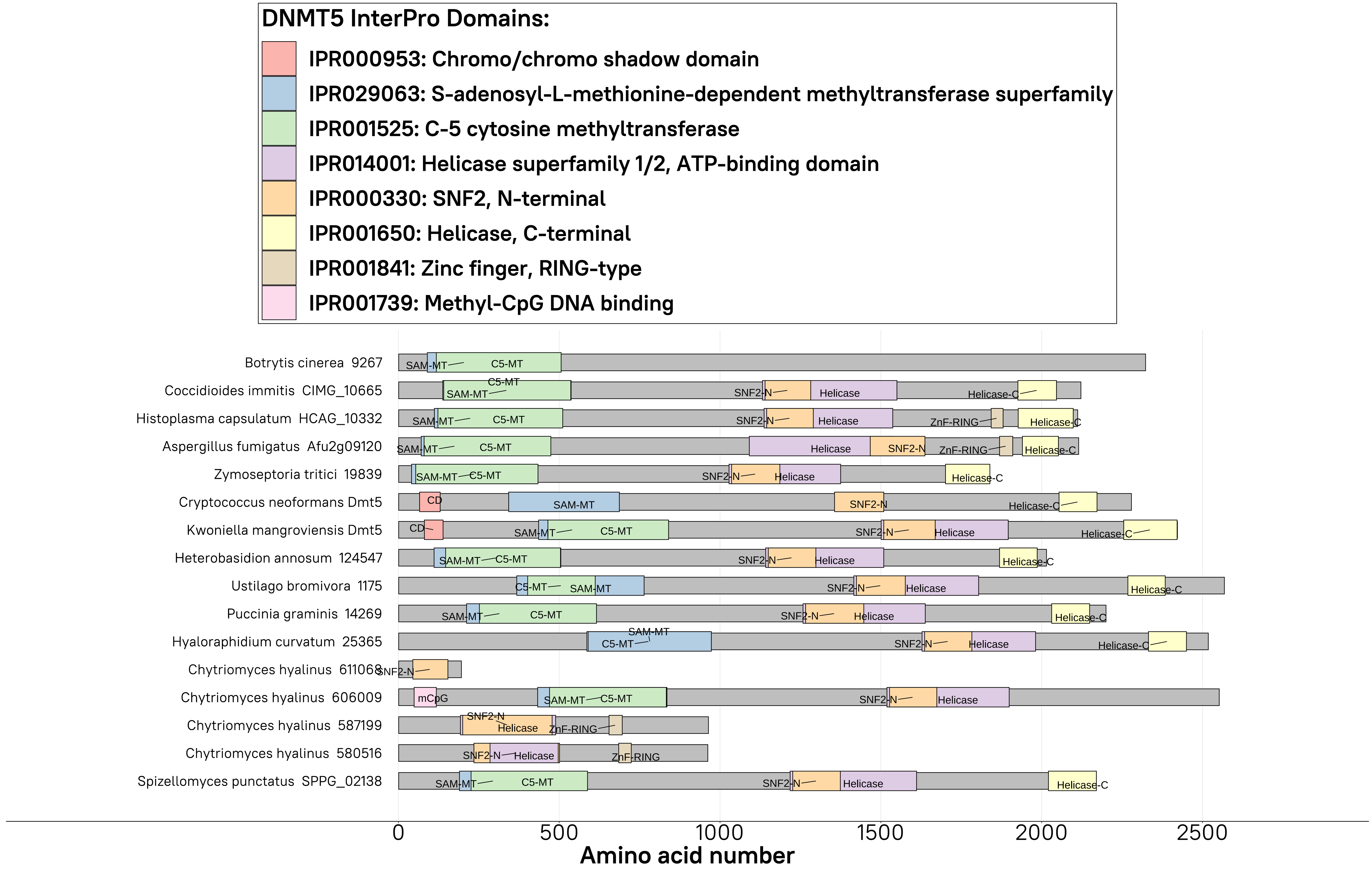
DNMT5 InterPro Domains:
IPR000953: Chromo/chromo shadow domain
IPR029063: S-adenosyl-L-methionine-dependent methyltransferase superfamily
IPR001525: C-5 cytosine methyltransferase
IPR014001: Helicase superfamily 1/2, ATP-binding domain
IPR000330: SNF2, N-terminal
IPR001650: Helicase, C-terminal
IPR001841: Zinc finger, RING-type
IPR001739: Methyl-CpG DNA binding
Botrytis cinerea 9267
Coccidioides immitis CIMG_10665
Histoplasma capsulatum HCAG_10332
Aspergillus fumigatus Afu2g09120
Zymoseptoria tritici 19839
Cryptococcus neoformans Dmt5
Kwoniella mangroviensis Dmt5
Heterobasidion annosum 124547
Ustilago bromivora 1175
Puccinia graminis 14269
Hyaloraphidium curvatum 25365
Chytriomyces hyalinus 611068
Chytriomyces hyalinus 606009
Chytriomyces hyalinus 587199
Chytriomyces hyalinus 580516
Spizellomyces punctatus SPPG_02138
SAM-MT
C5-MT
SNF2-N
Helicase
Helicase-C
ZnF-RING
CD
mCpG
0
500
1000
1500
2000
2500
Amino acid number

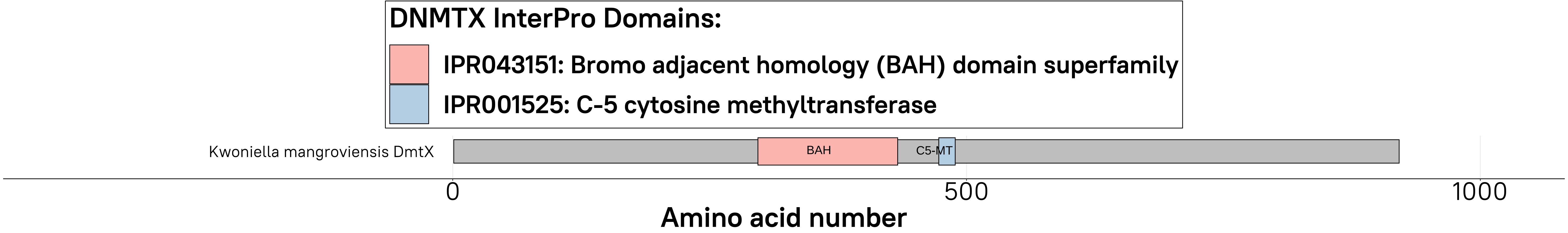
DNMTX InterPro Domains:
IPR043151: Bromo adjacent homology (BAH) domain superfamily
IPR001525: C-5 cytosine methyltransferase
Kwoniella mangroviensis DmtX
BAH
C5-MT
0
500
1000
Amino acid number

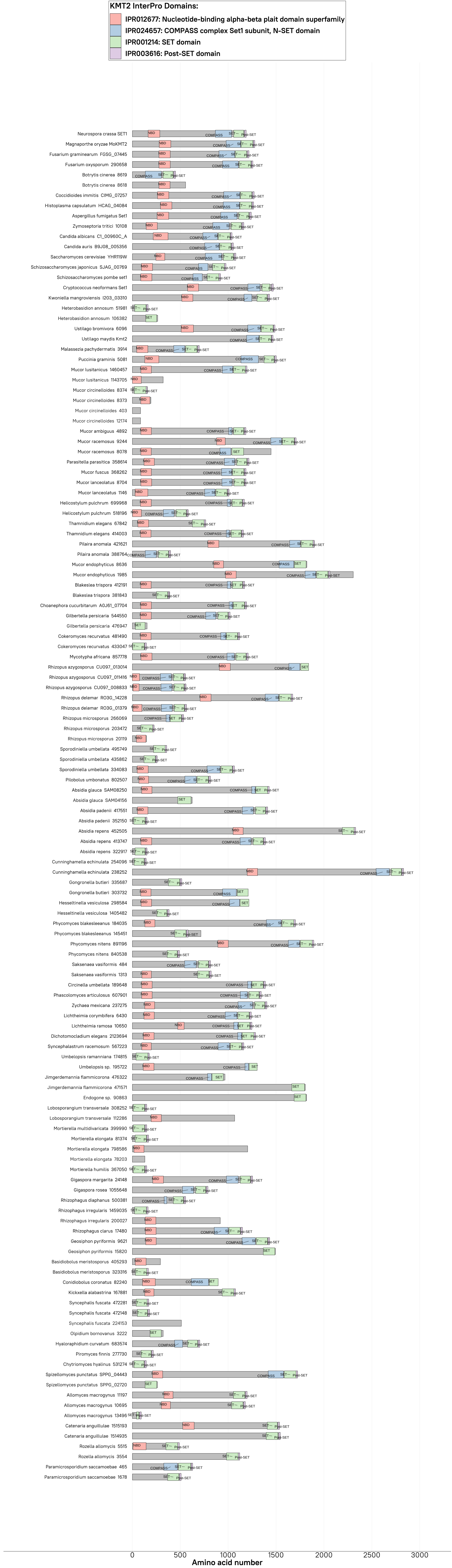
KMT2 InterPro Domains:
IPR012677: Nucleotide-binding alpha-beta plait domain superfamily
IPR024657: COMPASS complex Set1 subunit, N-SET domain
IPR001214: SET domain
IPR003616: Post-SET domain
Neurospora crassa SET1
Magnaporthe oryzae MoKMT2
Fusarium graminearum FGSG_07445
Fusarium oxysporum 290658
Botrytis cinerea 8619
Botrytis cinerea 8618
Coccidioides immitis CIMG_07257
Histoplasma capsulatum HCAG_04084
Aspergillus fumigatus Set1
Zymoseptoria tritici 10108
Candida albicans C1_00960C_A
Candida auris B9J08_005356
Saccharomyces cerevisiae YHR119W
Schizosaccharomyces japonicus SJAG_00769
Schizosaccharomyces pombe set1
Cryptococcus neoformans Set1
Kwoniella mangroviensis I203_03310
Heterobasidion annosum 51981
Heterobasidion annosum 106382
Ustilago bromivora 6096
Ustilago maydis Kmt2
Malassezia pachydermatis 3914
Puccinia graminis 5081
Mucor lusitanicus 1460457
Mucor lusitanicus 1143705
Mucor circinelloides 8374
Mucor circinelloides 8373
Mucor circinelloides 403
Mucor circinelloides 12174
Mucor ambiguus 4892
Mucor racemosus 9244
Mucor racemosus 8078
Parasitella parasitica 358614
Mucor fuscus 368262
Mucor lanceolatus 8704
Mucor lanceolatus 1146
Helicostylum pulchrum 699968
Helicostylum pulchrum 518196
Thamnidium elegans 67842
Thamnidium elegans 414003
Pilaira anomala 421621
Pilaira anomala 388764
Mucor endophyticus 8636
Mucor endophyticus 1985
Blakeslea trispora 412191
Blakeslea trispora 381843
Choanephora cucurbitarum A0J61_07704
Gilbertella persicaria 544550
Gilbertella persicaria 476947
Cokeromyces recurvatus 481490
Cokeromyces recurvatus 433047
Mycotypha africana 857778
Rhizopus azygosporus CU097_013014
Rhizopus azygosporus CU097_011416
Rhizopus azygosporus CU097_008833
Rhizopus delemar RO3G_14228
Rhizopus delemar RO3G_01379
Rhizopus microsporus 266069
Rhizopus microsporus 203472
Rhizopus microsporus 20119
Sporodiniella umbellata 495749
Sporodiniella umbellata 435862
Sporodiniella umbellata 334083
Pilobolus umbonatus 802507
Absidia glauca SAM08250
Absidia glauca SAM04156
Absidia padenii 417551
Absidia padenii 352150
Absidia repens 452505
Absidia repens 413747
Absidia repens 322917
Cunninghamella echinulata 254096
Cunninghamella echinulata 238252
Gongronella butleri 335687
Gongronella butleri 303732
Hesseltinella vesiculosa 298584
Hesseltinella vesiculosa 1405482
Phycomyces blakesleeanus 184035
Phycomyces blakesleeanus 145451
Phycomyces nitens 891196
Phycomyces nitens 840538
Saksenaea vasiformis 484
Saksenaea vasiformis 1313
Circinella umbellata 189648
Phascolomyces articulosus 607901
Zychaea mexicana 237275
Lichtheimia corymbifera 6430
Lichtheimia ramosa 10650
Dichotomocladium elegans 2123694
Syncephalastrum racemosum 567223
Umbelopsis ramanniana 174815
Umbelopsis sp. 195722
Jimgerdemannia flammicorona 476322
Jimgerdemannia flammicorona 471571
Endogone sp. 90863
Lobosporangium transversale 308252
Lobosporangium transversale 112286
Mortierella multidivaricata 399990
Mortierella elongata 81374
Mortierella elongata 798586
Mortierella elongata 78203
Mortierella humilis 367050
Gigaspora margarita 24148
Gigaspora rosea 1055648
Rhizophagus diaphanus 500381
Rhizophagus irregularis 1459035
Rhizophagus irregularis 200027
Rhizophagus clarus 17480
Geosiphon pyriformis 9621
Geosiphon pyriformis 15820
Basidiobolus meristosporus 405293
Basidiobolus meristosporus 323316
Conidiobolus coronatus 82240
Kickxella alabastrina 167881
Syncephalis fuscata 472281
Syncephalis fuscata 472148
Syncephalis fuscata 224153
Olpidium bornovanus 3222
Hyaloraphidium curvatum 683574
Piromyces finnis 277730
Chytriomyces hyalinus 531274
Spizellomyces punctatus SPPG_04443
Spizellomyces punctatus SPPG_02720
Allomyces macrogynus 11197
Allomyces macrogynus 10695
Allomyces macrogynus 13496
Catenaria anguillulae 1515193
Catenaria anguillulae 1514935
Rozella allomycis 5515
Rozella allomycis 3554
Paramicrosporidium saccamoebae 465
Paramicrosporidium saccamoebae 1678
0
500
1000
1500
2000
2500
3000
Amino acid number

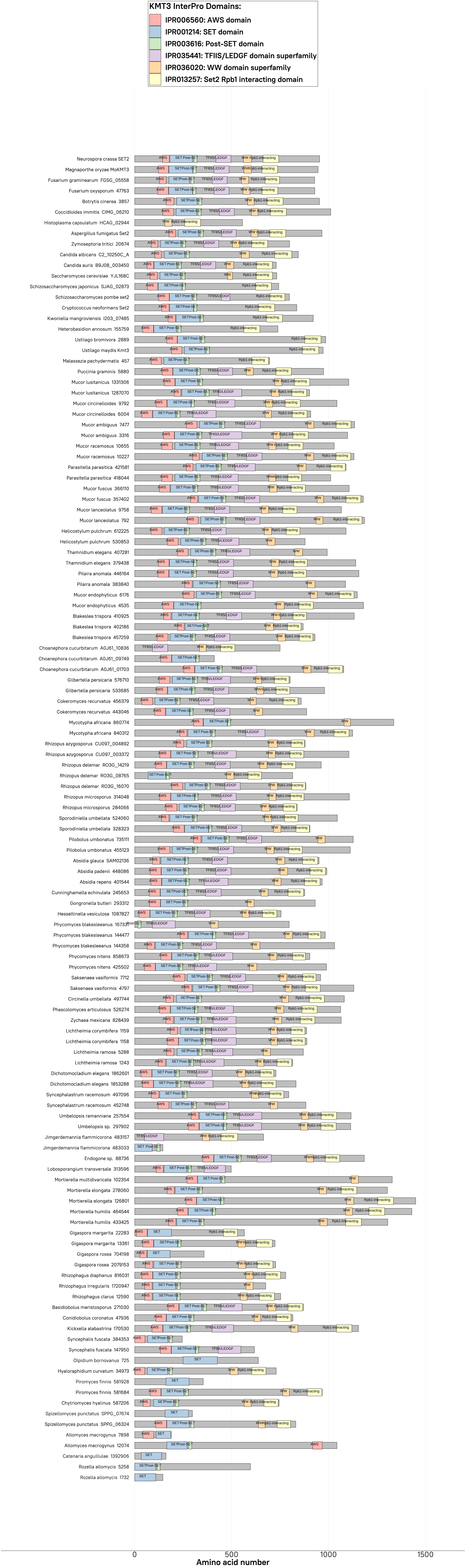
KMT3 InterPro Domains:
IPR006560: AWS domain
IPR001214: SET domain
IPR003616: Post-SET domain
IPR035441: TFIIS/LEDGF domain superfamily
IPR036020: WW domain superfamily
IPR013257: Set2 Rpb1 interacting domain
Neurospora crassa SET2
Magnaporthe oryzae MoKMT3
Fusarium graminearum FGSG_05558
Fusarium oxysporum 47763
Botrytis cinerea 3857
Coccidioides immitis CIMG_06210
Histoplasma capsulatum HCAG_02944
Aspergillus fumigatus Set2
Zymoseptoria tritici 20674
Candida albicans C2_10250C_A
Candida auris B9J08_003450
Saccharomyces cerevisiae YJL168C
Schizosaccharomyces japonicus SJAG_02873
Schizosaccharomyces pombe set2
Cryptococcus neoformans Set2
Kwoniella mangroviensis I203_07485
Heterobasidion annosum 155759
Ustilago bromivora 2889
Ustilago maydis Kmt3
Malassezia pachydermatis 457
Puccinia graminis 5880
Mucor lusitanicus 1331306
Mucor lusitanicus 1287070
Mucor circinelloides 9792
Mucor circinelloides 6004
Mucor ambiguus 7477
Mucor ambiguus 3316
Mucor racemosus 10655
Mucor racemosus 10227
Parasitella parasitica 421581
Parasitella parasitica 418044
Mucor fuscus 366110
Mucor fuscus 357402
Mucor lanceolatus 9758
Mucor lanceolatus 792
Helicostylum pulchrum 612225
Helicostylum pulchrum 530853
Thamnidium elegans 407281
Thamnidium elegans 379438
Pilaira anomala 446164
Pilaira anomala 383840
Mucor endophyticus 6176
Mucor endophyticus 4535
Blakeslea trispora 410925
Blakeslea trispora 402186
Blakeslea trispora 457259
Choanephora cucurbitarum A0J61_10836
Choanephora cucurbitarum A0J61_09749
Choanephora cucurbitarum A0J61_01703
Gilbertella persicaria 576710
Gilbertella persicaria 533685
Cokeromyces recurvatus 456379
Cokeromyces recurvatus 443046
Mycotypha africana 860774
Mycotypha africana 840312
Rhizopus azygosporus CU097_004892
Rhizopus azygosporus CU097_003372
Rhizopus delemar RO3G_14219
Rhizopus delemar RO3G_08765
Rhizopus delemar RO3G_16070
Rhizopus microsporus 314048
Rhizopus microsporus 284066
Sporodiniella umbellata 524060
Sporodiniella umbellata 328323
Pilobolus umbonatus 735111
Pilobolus umbonatus 455123
Absidia glauca SAM02136
Absidia padenii 448086
Absidia repens 401544
Cunninghamella echinulata 245653
Gongronella butleri 293312
Hesseltinella vesiculosa 1087827
Phycomyces blakesleeanus 167337
Phycomyces blakesleeanus 144477
Phycomyces blakesleeanus 144358
Phycomyces nitens 858673
Phycomyces nitens 425502
Saksenaea vasiformis 7712
Saksenaea vasiformis 4797
Circinella umbellata 497744
Phascolomyces articulosus 526274
Zychaea mexicana 828439
Lichtheimia corymbifera 1159
Lichtheimia corymbifera 1158
Lichtheimia ramosa 5288
Lichtheimia ramosa 1243
Dichotomocladium elegans 1862601
Dichotomocladium elegans 1853288
Syncephalastrum racemosum 497096
Syncephalastrum racemosum 452748
Umbelopsis ramanniana 257554
Umbelopsis sp. 297902
Jimgerdemannia flammicorona 483157
Jimgerdemannia flammicorona 483033
Endogone sp. 88736
Lobosporangium transversale 313596
Mortierella multidivaricata 102354
Mortierella elongata 278060
Mortierella elongata 126801
Mortierella humilis 464544
Mortierella humilis 433425
Gigaspora margarita 22283
Gigaspora margarita 13361
Gigaspora rosea 704198
Gigaspora rosea 2079153
Rhizophagus diaphanus 816031
Rhizophagus irregularis 1720947
Rhizophagus clarus 12590
Basidiobolus meristosporus 271030
Conidiobolus coronatus 47936
Kickxella alabastrina 170530
Syncephalis fuscata 384353
Syncephalis fuscata 147950
Olpidium bornovanus 725
Hyaloraphidium curvatum 34973
Piromyces finnis 581928
Piromyces finnis 581684
Chytriomyces hyalinus 587206
Spizellomyces punctatus SPPG_07674
Spizellomyces punctatus SPPG_06324
Allomyces macrogynus 7898
Allomyces macrogynus 12074
Catenaria anguillulae 1392906
Rozella allomycis 5258
Rozella allomycis 1732
0
500
1000
1500
Amino acid number

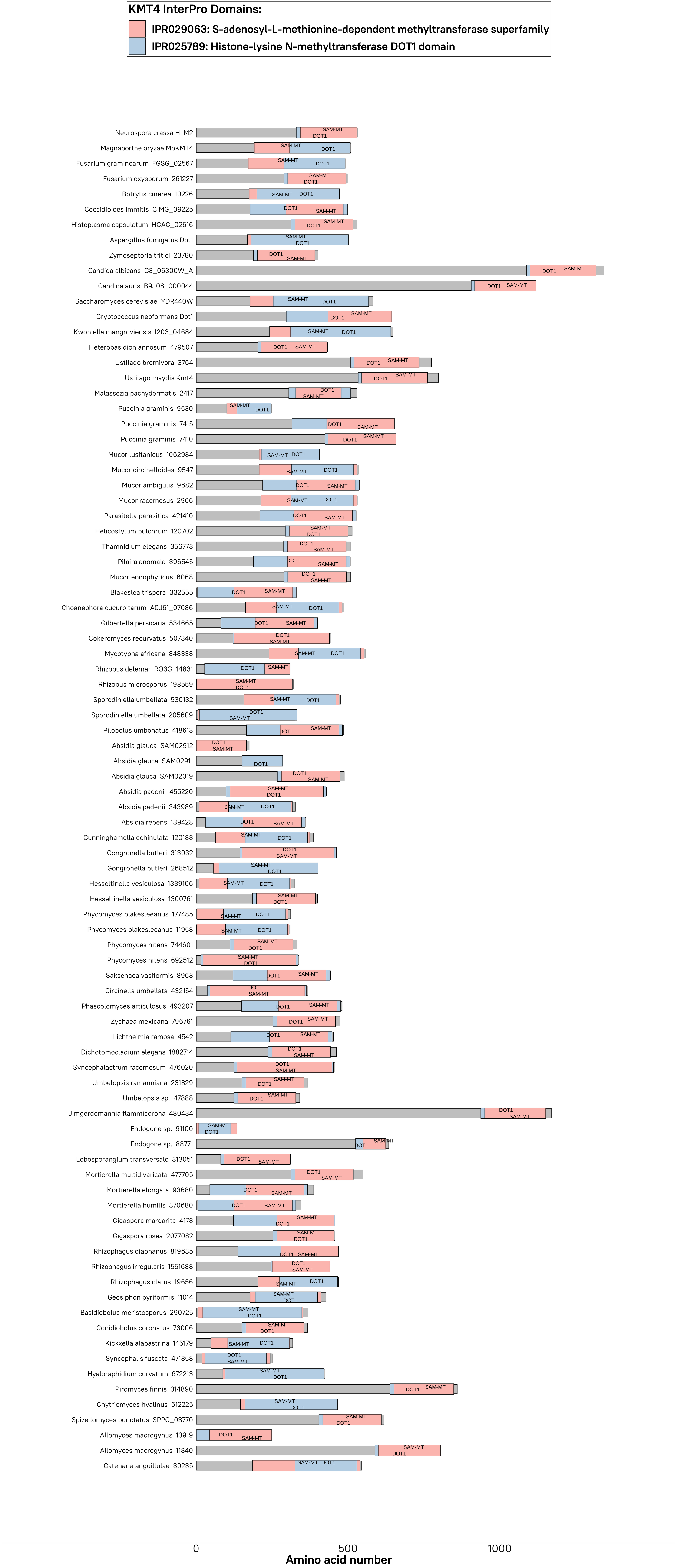
KMT4 InterPro Domains:
IPR029063: S-adenosyl-L-methionine-dependent methyltransferase superfamily
IPR025789: Histone-lysine N-methyltransferase DOT1 domain
Neurospora crassa HLM2
Magnaporthe oryzae MoKMT4
Fusarium graminearum FGSG_02567
Fusarium oxysporum 261227
Botrytis cinerea 10226
Coccidioides immitis CIMG_09225
Histoplasma capsulatum HCAG_02616
Aspergillus fumigatus Dot1
Zymoseptoria tritici 23780
Candida albicans C3_06300W_A
Candida auris B9J08_000044
Saccharomyces cerevisiae YDR440W
Cryptococcus neoformans Dot1
Kwoniella mangroviensis I203_04684
Heterobasidion annosum 479507
Ustilago bromivora 3764
Ustilago maydis Kmt4
Malassezia pachydermatis 2417
Puccinia graminis 9530
Puccinia graminis 7415
Puccinia graminis 7410
Mucor lusitanicus 1062984
Mucor circinelloides 9547
Mucor ambiguus 9682
Mucor racemosus 2966
Parasitella parasitica 421410
Helicostylum pulchrum 120702
Thamnidium elegans 356773
Pilaira anomala 396545
Mucor endophyticus 6068
Blakeslea trispora 332555
Choanephora cucurbitarum A0J61_07086
Gilbertella persicaria 534665
Cokeromyces recurvatus 507340
Mycotypha africana 848338
Rhizopus delemar RO3G_14831
Rhizopus microsporus 198559
Sporodiniella umbellata 530132
Sporodiniella umbellata 205609
Pilobolus umbonatus 418613
Absidia glauca SAM02912
Absidia glauca SAM02911
Absidia glauca SAM02019
Absidia padenii 455220
Absidia padenii 343989
Absidia repens 139428
Cunninghamella echinulata 120183
Gongronella butleri 313032
Gongronella butleri 268512
Hesseltinella vesiculosa 1339106
Hesseltinella vesiculosa 1300761
Phycomyces blakesleeanus 177485
Phycomyces blakesleeanus 11958
Phycomyces nitens 744601
Phycomyces nitens 692512
Saksenaea vasiformis 8963
Circinella umbellata 432154
Phascolomyces articulosus 493207
Zychaea mexicana 796761
Lichtheimia ramosa 4542
Dichotomocladium elegans 1882714
Syncephalastrum racemosum 476020
Umbelopsis ramanniana 231329
Umbelopsis sp. 47888
Jimgerdemannia flammicorona 480434
Endogone sp. 91100
Endogone sp. 88771
Lobosporangium transversale 313051
Mortierella multidivaricata 477705
Mortierella elongata 93680
Mortierella humilis 370680
Gigaspora margarita 4173
Gigaspora rosea 2077082
Rhizophagus diaphanus 819635
Rhizophagus irregularis 1551688
Rhizophagus clarus 19656
Geosiphon pyriformis 11014
Basidiobolus meristosporus 290725
Conidiobolus coronatus 73006
Kickxella alabastrina 145179
Syncephalis fuscata 471858
Hyaloraphidium curvatum 672213
Piromyces finnis 314890
Chytriomyces hyalinus 612225
Spizellomyces punctatus SPPG_03770
Allomyces macrogynus 13919
Allomyces macrogynus 11840
Catenaria anguillulae 30235
SAM-MT
DOT1
0
500
1000
Amino acid number

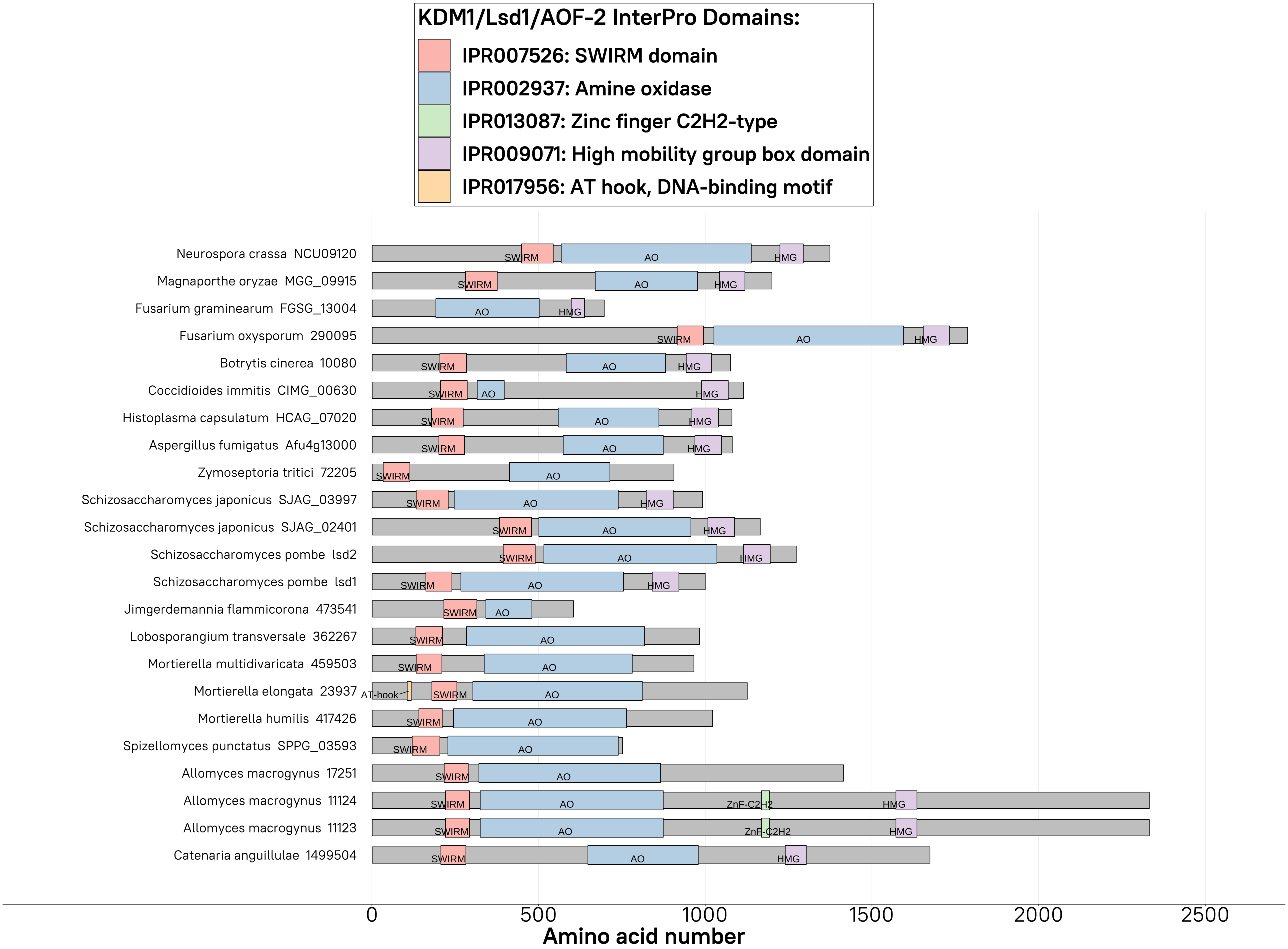

KDM1/Lsd1/AOF-2 InterPro Domains:
IPR007526: SWIRM domain
IPR002937: Amine oxidase
IPR013087: Zinc finger C2H2-type
IPR009071: High mobility group box domain
IPR017956: AT hook, DNA-binding motif
Neurospora crassa NCU09120
Magnaporthe oryzae MGG_09915
Fusarium graminearum FGSG_13004
Fusarium oxysporum 290095
Botrytis cinerea 10080
Coccidioides immitis CIMG_00630
Histoplasma capsulatum HCAG_07020
Aspergillus fumigatus Afu4g13000
Zymoseptoria tritici 72205
Schizosaccharomyces japonicus SJAG_03997
Schizosaccharomyces japonicus SJAG_02401
Schizosaccharomyces pombe lsd2
Schizosaccharomyces pombe lsd1
Jimgerdemannia flammicorona 473541
Lobosporangium transversale 362267
Mortierella multidivaricata 459503
Mortierella elongata 23937
Mortierella humilis 417426
Spizellomyces punctatus SPPG_03593
Allomyces macrogynus 17251
Allomyces macrogynus 11124
Allomyces macrogynus 11123
Catenaria anguillulae 1499504
SWIRM
AO
HMG
ZnF-C2H2
AT-hook
0
500
1000
1500
2000
2500
Amino acid number

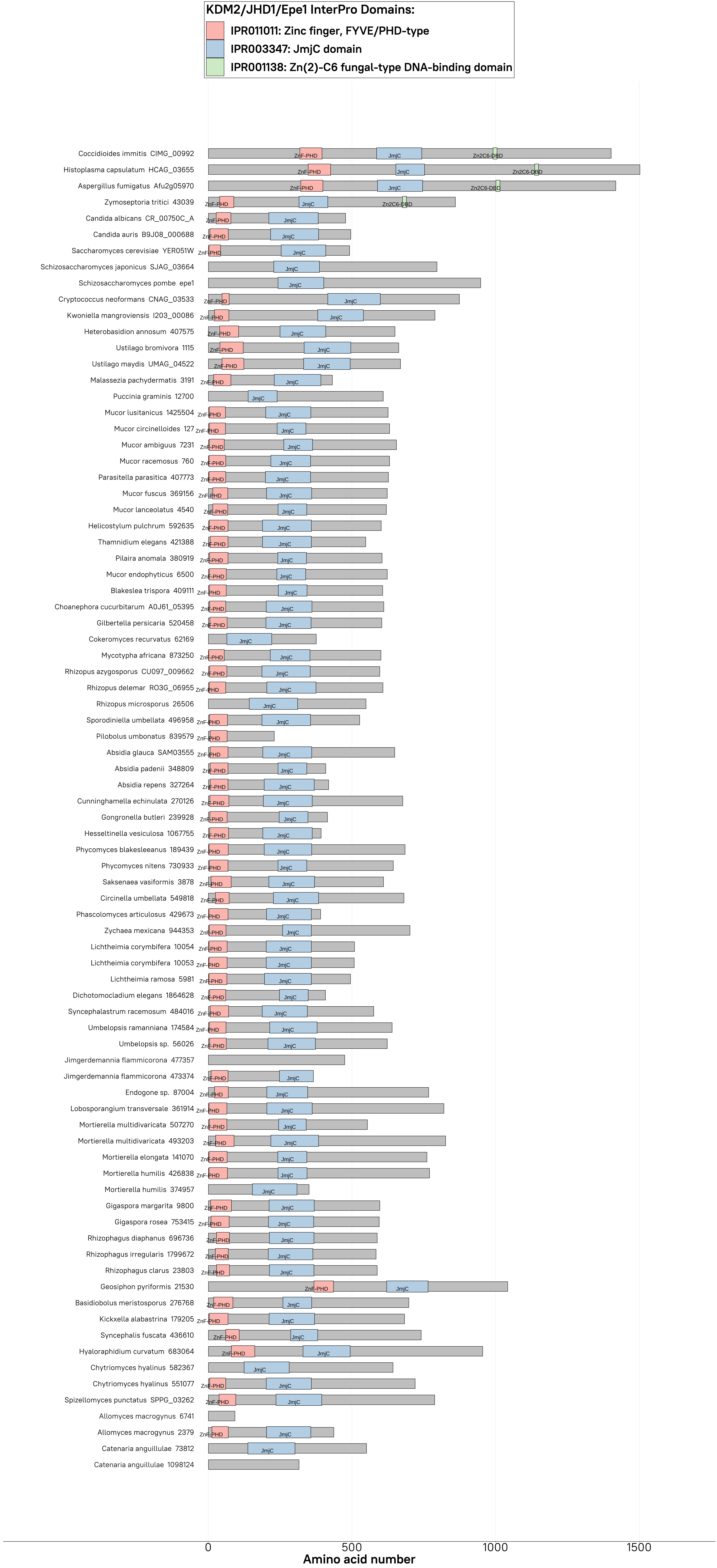
KDM2/JHD1/Epe1 InterPro Domains:
IPR011011: Zinc finger, FYVE/PHD-type
IPR003347: JmjC domain
IPR001138: Zn(2)-C6 fungal-type DNA-binding domain
Coccidioides immitis CIMG_00992
Histoplasma capsulatum HCAG_03655
Aspergillus fumigatus Afu2g05970
Zymoseptoria tritici 43039
Candida albicans CR_00750C_A
Candida auris B9J08_000688
Saccharomyces cerevisiae YER051W
Schizosaccharomyces japonicus SJAG_03664
Schizosaccharomyces pombe epe1
Cryptococcus neoformans CNAG_03533
Kwoniella mangroviensis I203_00086
Heterobasidion annosum 407575
Ustilago bromivora 1115
Ustilago maydis UMAG_04522
Malassezia pachydermatis 3191
Puccinia graminis 12700
Mucor lusitanicus 1425504
Mucor circinelloides 127
Mucor ambiguus 7231
Mucor racemosus 760
Parasitella parasitica 407773
Mucor fuscus 369156
Mucor lanceolatus 4540
Helicostylum pulchrum 592635
Thamnidium elegans 421388
Pilaira anomala 380919
Mucor endophyticus 6500
Blakeslea trispora 409111
Choanephora cucurbitarum A0J61_05395
Gilbertella persicaria 520458
Cokeromyces recurvatus 62169
Mycotypha africana 873250
Rhizopus azygosporus CU097_009662
Rhizopus delemar RO3G_06955
Rhizopus microsporus 26506
Sporodiniella umbellata 496958
Pilobolus umbonatus 839579
Absidia glauca SAM03555
Absidia padenii 348809
Absidia repens 327264
Cunninghamella echinulata 270126
Gongronella butleri 239928
Hesseltinella vesiculosa 1067755
Phycomyces blakesleeanus 189439
Phycomyces nitens 730933
Saksenaea vasiformis 3878
Circinella umbellata 549818
Phascolomyces articulosus 429673
Zychaea mexicana 944353
Lichtheimia corymbifera 10054
Lichtheimia corymbifera 10053
Lichtheimia ramosa 5981
Dichotomocladium elegans 1864628
Syncephalastrum racemosum 484016
Umbelopsis ramanniana 174584
Umbelopsis sp. 56026
Jimgerdemannia flammicorona 477357
Jimgerdemannia flammicorona 473374
Endogone sp. 87004
Lobosporangium transversale 361914
Mortierella multidivaricata 507270
Mortierella multidivaricata 493203
Mortierella elongata 141070
Mortierella humilis 426838
Mortierella humilis 374957
Gigaspora margarita 9800
Gigaspora rosea 753415
Rhizophagus diaphanus 696736
Rhizophagus irregularis 1799672
Rhizophagus clarus 23803
Geosiphon pyriformis 21530
Basidiobolus meristosporus 276768
Kickxella alabastrina 179205
Syncephalis fuscata 436610
Hyaloraphidium curvatum 683064
Chytriomyces hyalinus 582367
Chytriomyces hyalinus 551077
Spizellomyces punctatus SPPG_03262
Allomyces macrogynus 6741
Allomyces macrogynus 2379
Catenaria anguillulae 73812
Catenaria anguillulae 1098124
ZnF-PHD
JmjC
Zn2C6-DBD
0
500
1000
1500
Amino acid number

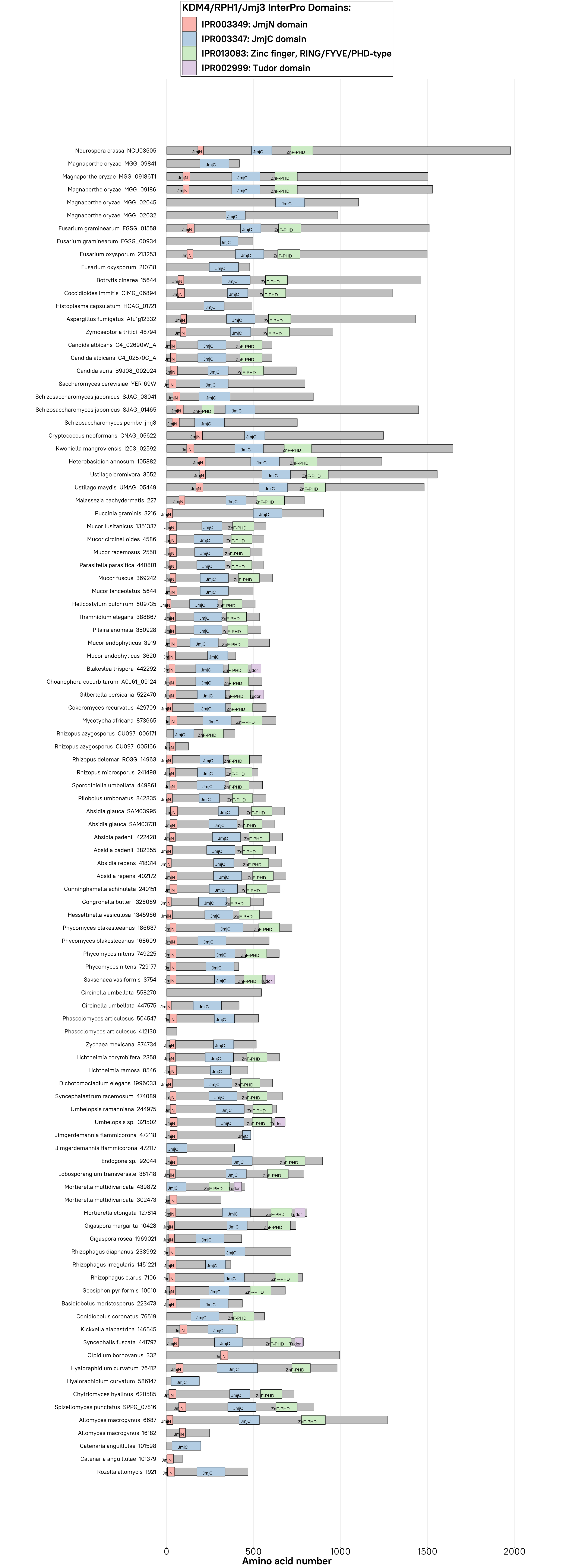
KDM4/RPH1/Jmj3 InterPro Domains:
IPR003349: JmjN domain
IPR003347: JmjC domain
IPR013083: Zinc finger, RING/FYVE/PHD-type
IPR002999: Tudor domain
Neurospora crassa NCU03505
Magnaporthe oryzae MGG_09841
Magnaporthe oryzae MGG_09186T1
Magnaporthe oryzae MGG_09186
Magnaporthe oryzae MGG_02045
Magnaporthe oryzae MGG_02032
Fusarium graminearum FGSG_01558
Fusarium graminearum FGSG_00934
Fusarium oxysporum 213253
Fusarium oxysporum 210718
Botrytis cinerea 15644
Coccidioides immitis CIMG_06894
Histoplasma capsulatum HCAG_01721
Aspergillus fumigatus Afu1g12332
Zymoseptoria tritici 48794
Candida albicans C4_02690W_A
Candida albicans C4_02570C_A
Candida auris B9J08_002024
Saccharomyces cerevisiae YER169W
Schizosaccharomyces japonicus SJAG_03041
Schizosaccharomyces japonicus SJAG_01465
Schizosaccharomyces pombe jmj3
Cryptococcus neoformans CNAG_05622
Kwoniella mangroviensis I203_02592
Heterobasidion annosum 105882
Ustilago bromivora 3652
Ustilago maydis UMAG_05449
Malassezia pachydermatis 227
Puccinia graminis 3216
Mucor lusitanicus 1351337
Mucor circinelloides 4586
Mucor racemosus 2550
Parasitella parasitica 440801
Mucor fuscus 369242
Mucor lanceolatus 5644
Helicostylum pulchrum 609735
Thamnidium elegans 388867
Pilaira anomala 350928
Mucor endophyticus 3919
Mucor endophyticus 3620
Blakeslea trispora 442292
Choanephora cucurbitarum A0J61_09124
Gilbertella persicaria 522470
Cokeromyces recurvatus 429709
Mycotypha africana 873665
Rhizopus azygosporus CU097_006171
Rhizopus azygosporus CU097_005166
Rhizopus delemar RO3G_14963
Rhizopus microsporus 241498
Sporodiniella umbellata 449861
Pilobolus umbonatus 842835
Absidia glauca SAM03995
Absidia glauca SAM03731
Absidia padenii 422428
Absidia padenii 382355
Absidia repens 418314
Absidia repens 402172
Cunninghamella echinulata 240151
Gongronella butleri 326069
Hesseltinella vesiculosa 1345966
Phycomyces blakesleeanus 186637
Phycomyces blakesleeanus 168609
Phycomyces nitens 749225
Phycomyces nitens 729177
Saksenaea vasiformis 3754
Circinella umbellata 558270
Circinella umbellata 447575
Phascolomyces articulosus 504547
Phascolomyces articulosus 412130
Zychaea mexicana 874734
Lichtheimia corymbifera 2358
Lichtheimia ramosa 8546
Dichotomocladium elegans 1996033
Syncephalastrum racemosum 474089
Umbelopsis ramanniana 244975
Umbelopsis sp. 321502
Jimgerdemannia flammicorona 472118
Jimgerdemannia flammicorona 472117
Endogone sp. 92044
Lobosporangium transversale 361718
Mortierella multidivaricata 439872
Mortierella multidivaricata 302473
Mortierella elongata 127814
Gigaspora margarita 10423
Gigaspora rosea 1969021
Rhizophagus diaphanus 233992
Rhizophagus irregularis 1451221
Rhizophagus clarus 7106
Geosiphon pyriformis 10010
Basidiobolus meristosporus 223473
Conidiobolus coronatus 76519
Kickxella alabastrina 146545
Syncephalis fuscata 441797
Olpidium bornovanus 332
Hyaloraphidium curvatum 76412
Hyaloraphidium curvatum 586147
Chytriomyces hyalinus 620585
Spizellomyces punctatus SPPG_07816
Allomyces macrogynus 6687
Allomyces macrogynus 16182
Catenaria anguillulae 101598
Catenaria anguillulae 101379
Rozella allomycis 1921
0
500
1000
1500
2000
Amino acid number

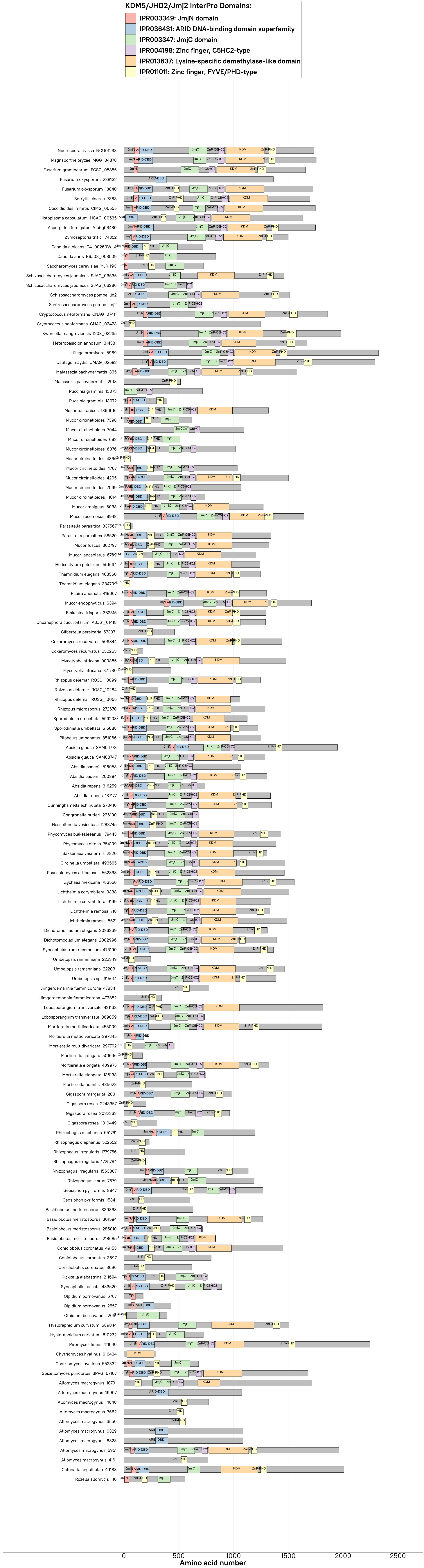

KDM5/JHD2/Jmj2 InterPro Domains:
IPR003349: JmjN domain
IPR036431: ARID DNA-binding domain superfamily
IPR003347: JmjC domain
IPR004198: Zinc finger, C5HC2-type
IPR013637: Lysine-specific demethylase-like domain
IPR011011: Zinc finger, FYVE/PHD-type
Neurospora crassa NCU01238
Magnaporthe oryzae MGG_04878
Fusarium graminearum FGSG_05855
Fusarium oxysporum 238132
Fusarium oxysporum 18840
Botrytis cinerea 7388
Coccidioides immitis CIMG_06555
Histoplasma capsulatum HCAG_00535
Aspergillus fumigatus Afu5g03430
Zymoseptoria tritici 74352
Candida albicans C4_00260W_A
Candida auris B9J08_003509
Saccharomyces cerevisiae YJR119C
Schizosaccharomyces japonicus SJAG_03635
Schizosaccharomyces japonicus SJAG_03286
Schizosaccharomyces pombe lid2
Schizosaccharomyces pombe jmj2
Cryptococcus neoformans CNAG_07411
Cryptococcus neoformans CNAG_03423
Kwoniella mangroviensis I203_02265
Heterobasidion annosum 314581
Ustilago bromivora 5989
Ustilago maydis UMAG_02582
Malassezia pachydermatis 335
Malassezia pachydermatis 2918
Puccinia graminis 13073
Puccinia graminis 13072
Mucor lusitanicus 1396016
Mucor circinelloides 7398
Mucor circinelloides 7044
Mucor circinelloides 693
Mucor circinelloides 6876
Mucor circinelloides 4866
Mucor circinelloides 4707
Mucor circinelloides 4205
Mucor circinelloides 2069
Mucor circinelloides 11014
Mucor ambiguus 6038
Mucor racemosus 8948
Parasitella parasitica 337567
Parasitella parasitica 58520
Mucor fuscus 362797
Mucor lanceolatus 6795
Helicostylum pulchrum 551694
Thamnidium elegans 463560
Thamnidium elegans 334709
Pilaira anomala 419067
Mucor endophyticus 6394
Blakeslea trispora 382515
Choanephora cucurbitarum A0J61_01418
Gilbertella persicaria 573071
Cokeromyces recurvatus 506344
Cokeromyces recurvatus 250263
Mycotypha africana 909885
Mycotypha africana 871780
Rhizopus delemar RO3G_13099
Rhizopus delemar RO3G_10284
Rhizopus delemar RO3G_10055
Rhizopus microsporus 272670
Sporodiniella umbellata 559203
Sporodiniella umbellata 515088
Pilobolus umbonatus 851066
Absidia glauca SAM04778
Absidia glauca SAM03747
Absidia padenii 516053
Absidia padenii 200384
Absidia repens 316259
Absidia repens 137177
Cunninghamella echinulata 270410
Gongronella butleri 236100
Hesseltinella vesiculosa 1283745
Phycomyces blakesleeanus 179443
Phycomyces nitens 754109
Saksenaea vasiformis 2820
Circinella umbellata 493565
Phascolomyces articulosus 562333
Zychaea mexicana 783556
Lichtheimia corymbifera 9338
Lichtheimia corymbifera 9199
Lichtheimia ramosa 718
Lichtheimia ramosa 5621
Dichotomocladium elegans 2033269
Dichotomocladium elegans 2002996
Syncephalastrum racemosum 476190
Umbelopsis ramanniana 222349
Umbelopsis ramanniana 222031
Umbelopsis sp. 315614
Jimgerdemannia flammicorona 478341
Jimgerdemannia flammicorona 473852
Lobosporangium transversale 421168
Lobosporangium transversale 369059
Mortierella multidivaricata 453009
Mortierella multidivaricata 297845
Mortierella multidivaricata 297792
Mortierella elongata 501696
Mortierella elongata 409975
Mortierella elongata 136138
Mortierella humilis 435623
Gigaspora margarita 2001
Gigaspora rosea 2243357
Gigaspora rosea 2032333
Gigaspora rosea 1010449
Rhizophagus diaphanus 651761
Rhizophagus diaphanus 522552
Rhizophagus irregularis 1779756
Rhizophagus irregularis 1725784
Rhizophagus irregularis 1563307
Rhizophagus clarus 7879
Geosiphon pyriformis 8847
Geosiphon pyriformis 15341
Basidiobolus meristosporus 339863
Basidiobolus meristosporus 301594
Basidiobolus meristosporus 285010
Basidiobolus meristosporus 218685
Conidiobolus coronatus 49153
Conidiobolus coronatus 3697
Conidiobolus coronatus 3696
Kickxella alabastrina 211694
Syncephalis fuscata 433520
Olpidium bornovanus 6767
Olpidium bornovanus 2557
Olpidium bornovanus 2091
Hyaloraphidium curvatum 689844
Hyaloraphidium curvatum 610232
Piromyces finnis 411040
Chytriomyces hyalinus 616434
Chytriomyces hyalinus 552332
Spizellomyces punctatus SPPG_07107
Allomyces macrogynus 18791
Allomyces macrogynus 16907
Allomyces macrogynus 14640
Allomyces macrogynus 7662
Allomyces macrogynus 6550
Allomyces macrogynus 6329
Allomyces macrogynus 6328
Allomyces macrogynus 5951
Allomyces macrogynus 4161
Catenaria anguillulae 49188
Rozella allomycis 110
0
500
1000
1500
2000
2500
Amino acid number
