## Supplementary material for "Heterochromatin and RNAi act independently to ensure genome stability in Mucorales human fungal pathogens": Dataset S3

**Dataset S3.** Major transposable element families and subfamilies, their ORFs, and protein domain configurations. The full-length, scaled nucleotide sequences of the major transposable element families and subfamilies identified in *M. lusitanicus* are depicted. Each family shows ORF prediction (gray arrowed blocks), as well as encoded InterPro protein domains (color-coded rectangles).

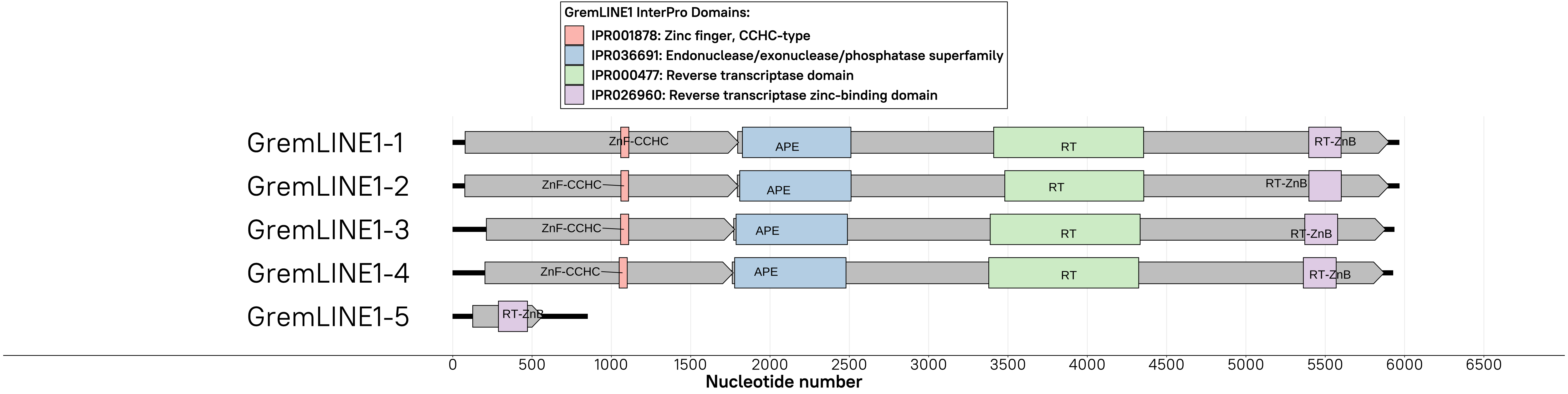

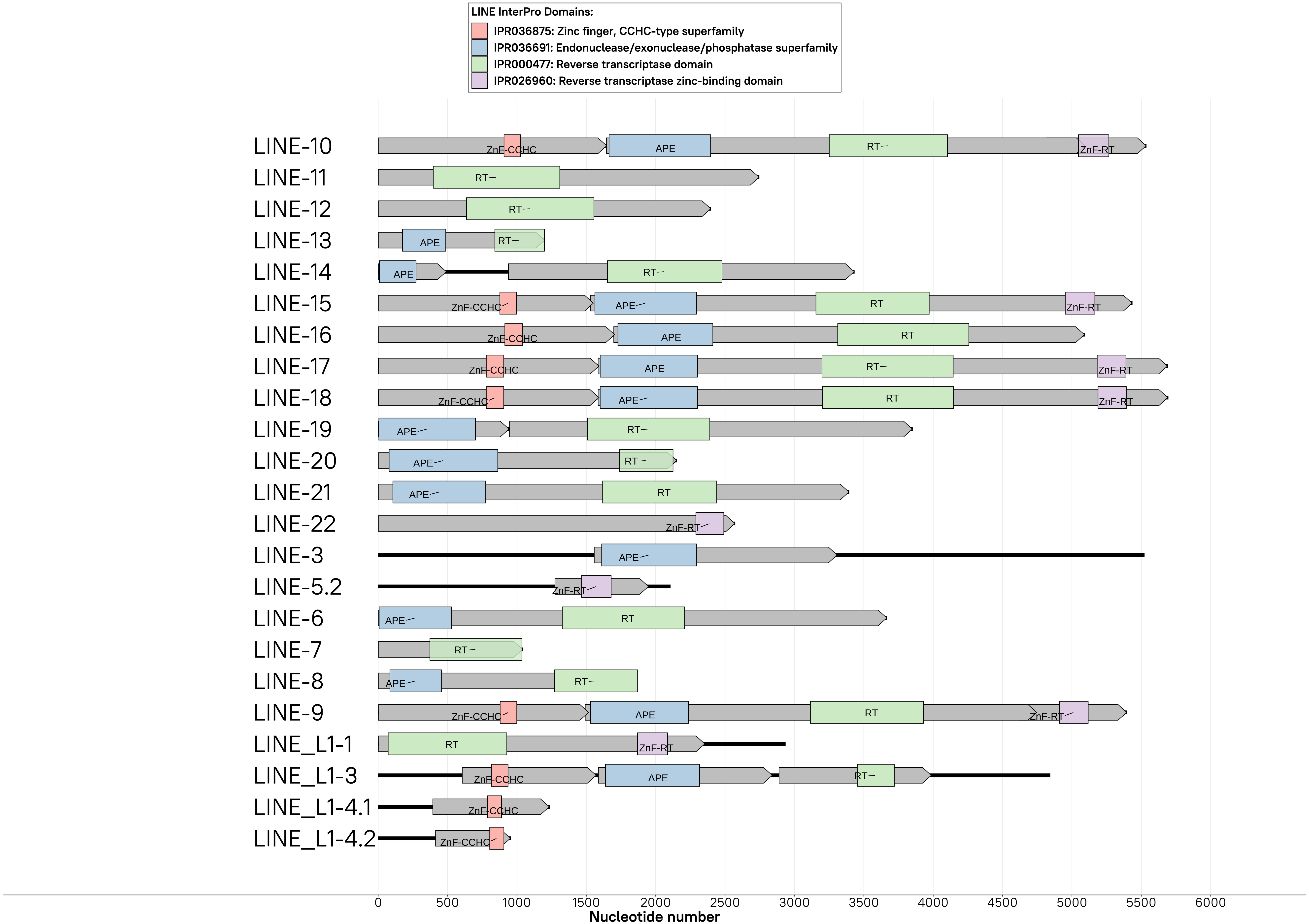
LINE InterPro Domains:
IPR036875: Zinc finger, CCHC-type superfamily
IPR036691: Endonuclease/exonuclease/phosphatase superfamily
IPR000477: Reverse transcriptase domain
IPR026960: Reverse transcriptase zinc-binding domain
LINE-10
LINE-11
LINE-12
LINE-13
LINE-14
LINE-15
LINE-16
LINE-17
LINE-18
LINE-19
LINE-20
LINE-21
LINE-22
LINE-3
LINE-5.2
LINE-6
LINE-7
LINE-8
LINE-9
LINE_L1-1
LINE_L1-3
LINE_L1-4.1
LINE_L1-4.2
ZnF-CCHC
APE
RT
ZnF-RT
0
500
1000
1500
2000
2500
3000
3500
4000
4500
5000
5500
6000
Nucleotide number

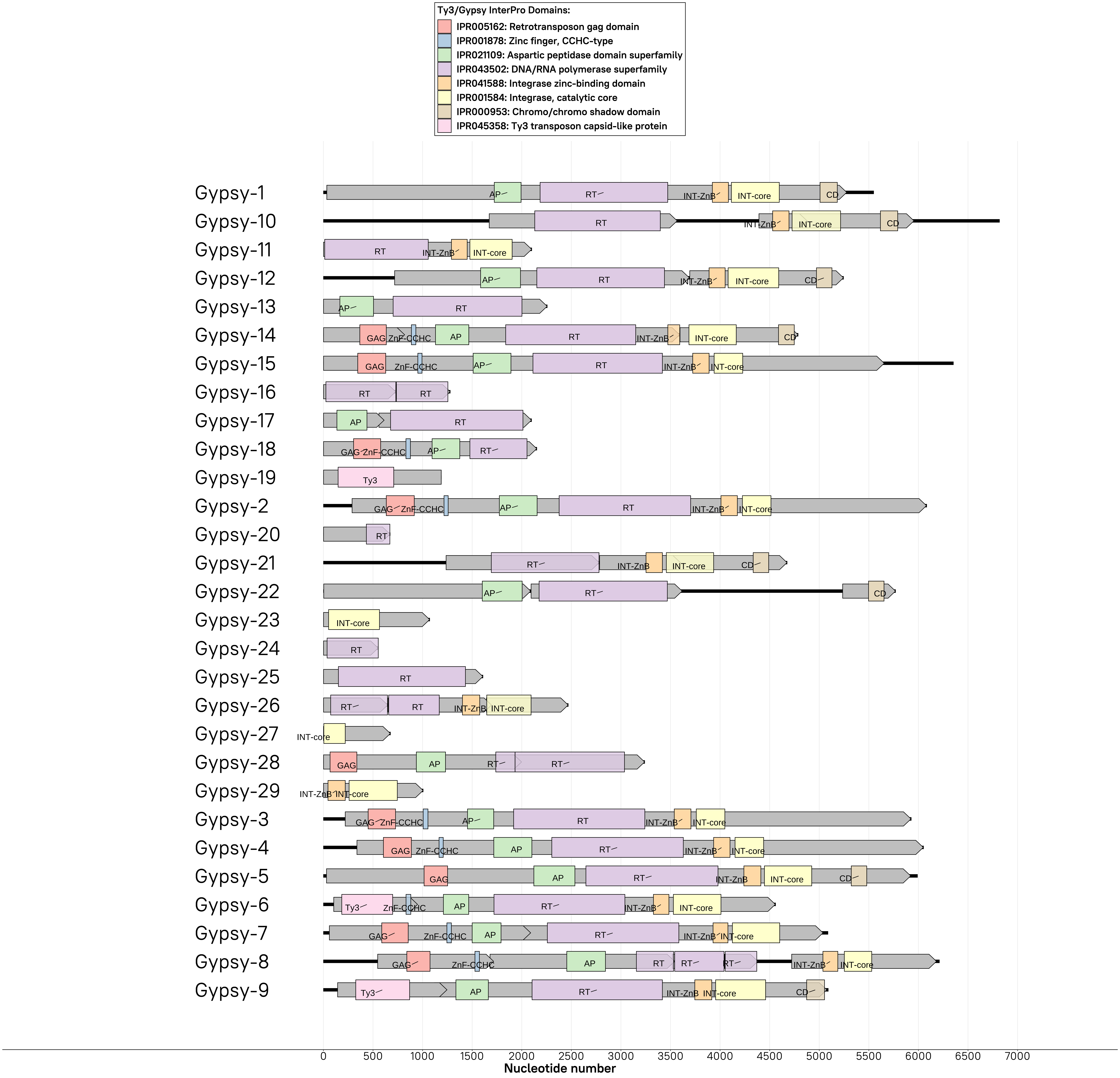

Ty3/Gypsy InterPro Domains:
IPR005162: Retrotransposon gag domain
IPR001878: Zinc finger, CCHC-type
IPR021109: Aspartic peptidase domain superfamily
IPR043502: DNA/RNA polymerase superfamily
IPR041588: Integrase zinc-binding domain
IPR001584: Integrase, catalytic core
IPR000953: Chromo/chromo shadow domain
IPR045358: Ty3 transposon capsid-like protein
Gypsy-1
Gypsy-10
Gypsy-11
Gypsy-12
Gypsy-13
Gypsy-14
Gypsy-15
Gypsy-16
Gypsy-17
Gypsy-18
Gypsy-19
Gypsy-2
Gypsy-20
Gypsy-21
Gypsy-22
Gypsy-23
Gypsy-24
Gypsy-25
Gypsy-26
Gypsy-27
Gypsy-28
Gypsy-29
Gypsy-3
Gypsy-4
Gypsy-5
Gypsy-6
Gypsy-7
Gypsy-8
Gypsy-9
0
500
1000
1500
2000
2500
3000
3500
4000
4500
5000
5500
6000
6500
7000
Nucleotide number

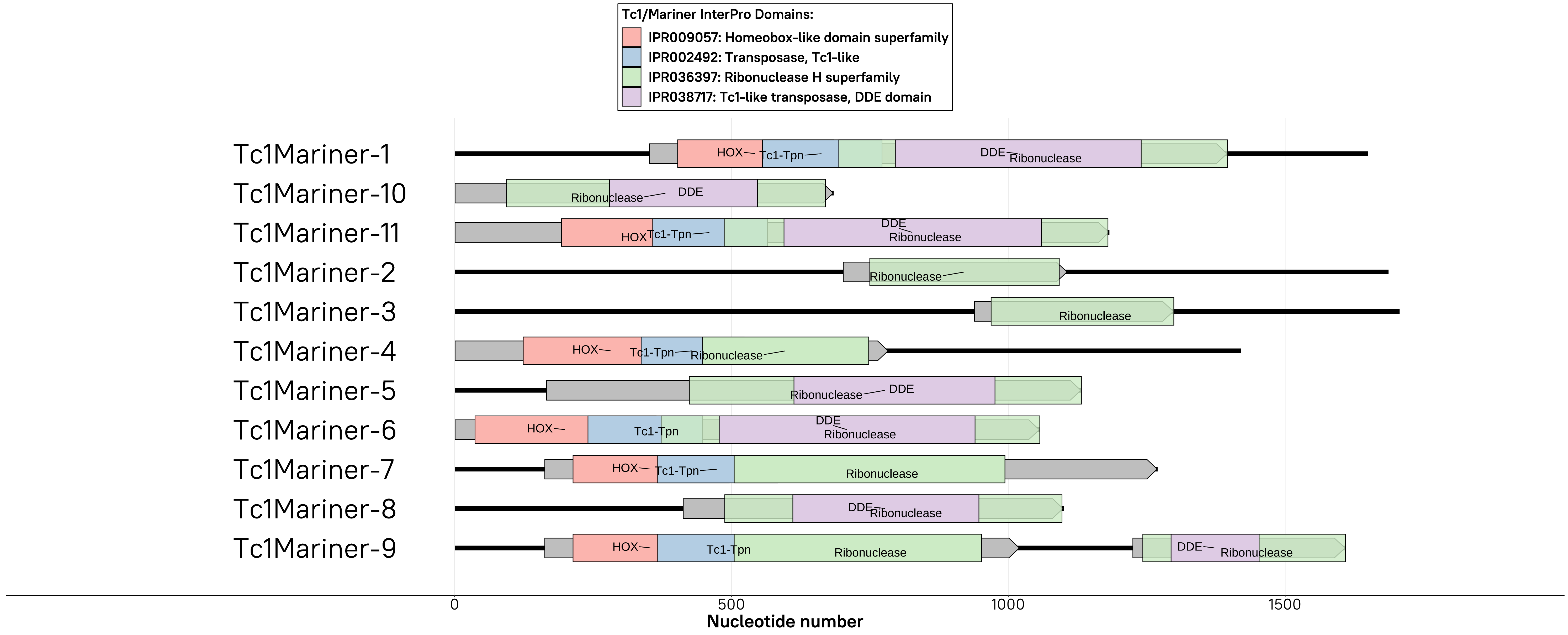
Tc1/Mariner InterPro Domains:
IPR009057: Homeobox-like domain superfamily
IPR002492: Transposase, Tc1-like
IPR036397: Ribonuclease H superfamily
IPR038717: Tc1-like transposase, DDE domain
Tc1Mariner-1
Tc1Mariner-10
Tc1Mariner-11
Tc1Mariner-2
Tc1Mariner-3
Tc1Mariner-4
Tc1Mariner-5
Tc1Mariner-6
Tc1Mariner-7
Tc1Mariner-8
Tc1Mariner-9
HOX
Tc1-Tpn
DDE
Ribonuclease
0
500
1000
1500
Nucleotide number

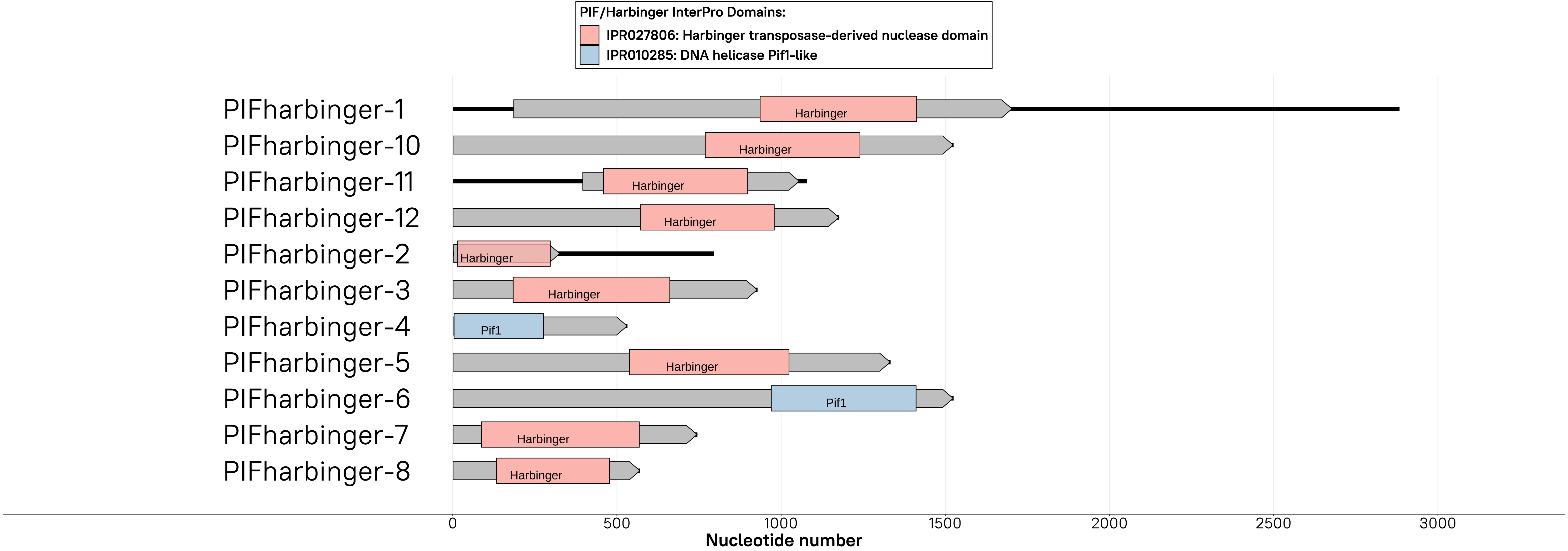
PIF/Harbinger InterPro Domains:
IPR027806: Harbinger transposase-derived nuclease domain
IPR010285: DNA helicase Pif1-like
PIFharbinger-1
PIFharbinger-10
PIFharbinger-11
PIFharbinger-12
PIFharbinger-2
PIFharbinger-3
PIFharbinger-4
PIFharbinger-5
PIFharbinger-6
PIFharbinger-7
PIFharbinger-8
Harbinger
Pif1
0
500
1000
1500
2000
2500
3000
Nucleotide number

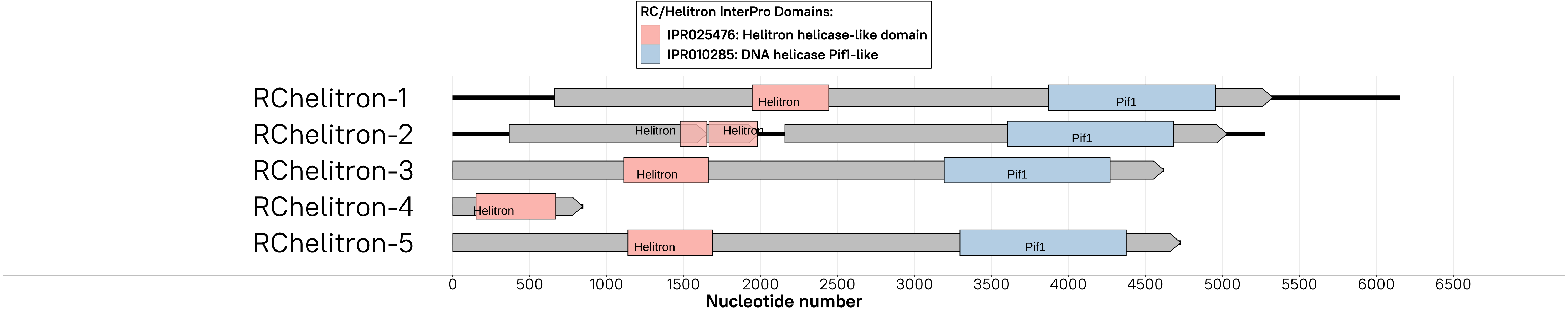
RC/Helitron InterPro Domains:
IPR025476: Helitron helicase-like domain
IPR010285: DNA helicase Pif1-like
RChelitron-1
RChelitron-2
RChelitron-3
RChelitron-4
RChelitron-5
Helitron
Pif1
0
500
1000
1500
2000
2500
3000
3500
4000
4500
5000
5500
6000
6500
Nucleotide number
